# Spatial mapping of RNA turnover kinetics and regulatory landscapes of mRNA stability in the mammalian brain

**DOI:** 10.64898/2026.01.29.702431

**Authors:** Qi Qiu, Hongjie Zhang, Zijie Xia, William Gao, Julie Leu, Dongming Liang, Ying Li, Yijing Su, Emily Feierman, Erin Van Horn, Guo-li Ming, Erica Korb, Hongjun Song, Zhaolan Zhou, Hao Wu

## Abstract

Spatial and activity-dependent gene regulation in the mammalian brain requires coordinated control of RNA synthesis and degradation ^1,2^, yet spatially resolved measurement of RNA turnover kinetics in complex tissues remains technically challenging ^3^. Here, we present *spatial NT-seq*, an approach that integrates transgenesis-free metabolic RNA labeling with *in situ* chemical recoding to spatially co-map RNA abundance and turnover kinetics in the mouse brain. By distinguishing newly synthesized from pre-existing RNAs, this method reveals spatially resolved transcriptional and post-transcriptional responses to electroconvulsive stimulation (ECS), a treatment for refractory depression. We uncover pronounced spatial heterogeneity in RNA turnover, with the dentate gyrus (DG) exhibiting elevated basal RNA turnover and robust ECS-induced responses. These findings reveal a “kinetics scaling” mechanism of coordinated regulation of RNA synthesis and decay, by which DG cells can rapidly remodel their transcript pools in responses to external stimuli or differentiation signals ^4,5^. Machine learning applied to *in vivo* RNA kinetics landscapes further identifies sequence features and post-transcriptional regulators underlying region-and cell-type-specific control of mRNA stability. Together, this integrated experimental and computational framework, *in vivo Timescope*, enables transcriptome-wide mapping of RNA turnover kinetics and the regulatory architecture of RNA stability across spatial and cellular contexts, providing new insights into the spatiotemporal regulation of RNA dynamics in brain function and disease.

## Main

The complex functions of the mammalian brain arise from diverse cell types organized into spatially distinct circuits and regions ^6^. This intricate architecture demands precise spatiotemporal control of gene expression, regulated at both transcriptional and post-transcriptional levels ^7^. RNA turnover kinetics, which governs how rapidly transcripts are produced and degraded, critically shapes steady-state transcriptomes and governs the temporal precision of gene expression responses ^2,8^. Rapid RNA turnover enables cells to quickly adapt to developmental cues ^9,10^ and environmental stimuli ^4^ such as neuronal activation. In contrast, slow turnover supports long-range RNA transport for localized protein synthesis in distal neuronal compartments ^11^ and stabilizes long-lived non-coding RNAs involved in maintaining chromatin integrity in the brain ^12^. Dysregulation of RNA turnover and stability has been implicated in a range of neurodevelopmental, neuropsychiatric and neurodegenerative diseases ^7,10,13–15^, highlighting the need to map RNA turnover kinetics and stability regulation *in vivo* across the spatial and cellular context of the brain.

Standard single-cell and spatial transcriptomics technologies provide only total RNA abundance, a static snapshot reflecting the equilibrium between RNA synthesis and degradation, thereby obscuring the underlying kinetic regulation ^3^. Metabolic RNA labeling can distinguish newly transcribed RNAs from the pre-existing pool, enabling quantitative measurement of RNA turnover kinetics through ‘single-pulse’ or ‘pulse-chase’ labeling strategies ^16^. While recent approaches combining metabolic labeling with single-cell RNA sequencing (scRNA-seq) or imaging methods have enabled cell-type specific RNA turnover kinetics measurements ^17–22^, they remain largely restricted to analysis of *in vitro* or *ex vivo* systems. Thus, the spatially resolved landscape of *in vivo* RNA kinetics, and the regulatory impact of sequence features or post-transcriptional regulators on RNA stability, remain largely unexplored.

Cell-permeable nucleoside analogs such as 4-thiouracil (4tU) ^23–25^, 4-thiouridine (4sU) ^26,27^ and 5-ethynyluridine (5-EU) ^28^ have been used to label newly transcribed RNAs *in vivo*, yet their utility for studying RNA turnover and stability is not well established. The most widely used *in vivo* RNA labeling method, ‘TU tagging’, employs transgenic expression of *Toxoplasma gondii* uracil phosphoribosyltransferase (UPRT) ^23,24^ to convert 4tU to 4-thiouridine monophosphate (4-thio-UMP), enabling its incorporation into nascent RNAs in specific cell-types. However, the requirement for transgenic mouse models limits its scalability and broader application.

In this study, we optimized and validated a transgenesis-free *in vivo* metabolic RNA labeling strategy using 4sU to quantify cell-type-specific RNA turnover in both embryonic and postnatal mouse brains. We then combined this *in vivo* labeling approach with *in situ* chemical recoding to develop spatial <u>N</u>ew RNA <u>T</u>agging <u>Seq</u>uencing (spatial NT-Seq), a first-in-class method that enables simultaneous spatial mapping of RNA abundance and turnover kinetics in intact tissues. Applied to mouse brains following electroconvulsive stimulation (ECS), a clinically effective treatment for refractory depression, spatial NT-seq disentangled the distinct contributions from RNA synthesis and degradation to ECS-induced gene expression changes across brain regions, providing new mechanistic insights into molecular effects of ECS. Moreover, spatial NT-seq revealed striking regional heterogeneity in brain RNA turnover kinetics and identified the dentate gyrus (DG) as a spatial hotspot of RNA turnover. The coordinated up-regulation of RNA synthesis and decay in this brain region likely facilitates DG cells to rapidly adjust protein output from existing transcript pools, supporting their intrinsic plasticity in responses to continuous neurogenesis and external stimuli such as learning and ECS. Together, by integrating transgenesis-free RNA labeling, spatial multimodal transcriptomics (i.e., co-detection of newly-synthesized, pre-existing, and total RNAs), and computational modeling, our “*in vivo Timescope*” framework provides a powerful platform to dissect how transcriptional and post-transcriptional regulation jointly shape gene expression landscapes in the brain.

### Benchmarking transgenesis-free metabolic RNA labeling in mouse brains

The lack of systematic validation has limited the application of *in vivo* metabolic RNA labeling for studying RNA turnover and stability in tissues. Although 4sU has been shown to enable direct RNA labeling in non-neural tissues ^26,27^, its labeling efficiency and potential toxicity in the brain remain uncertain. To address these gaps and enable analysis of more challenging brain cell types ^29^, we further optimized droplet-based single-cell metabolically labeled new RNA tagging sequencing (scNT-seq) ^21^ to enhance both the chemical conversion efficiency of 4sU in labeled new RNAs and overall library complexity (scNT-seq2 in **Fig. 1a**, **Extended Data Fig. 1a-c, Supplementary Fig. 1 and note 1**). Using scNT-seq2, we systematically validated the utility of transgenesis-free 4sU labeling for profiling RNA turnover in embryonic and postnatal mouse brains.

**Fig 1.**
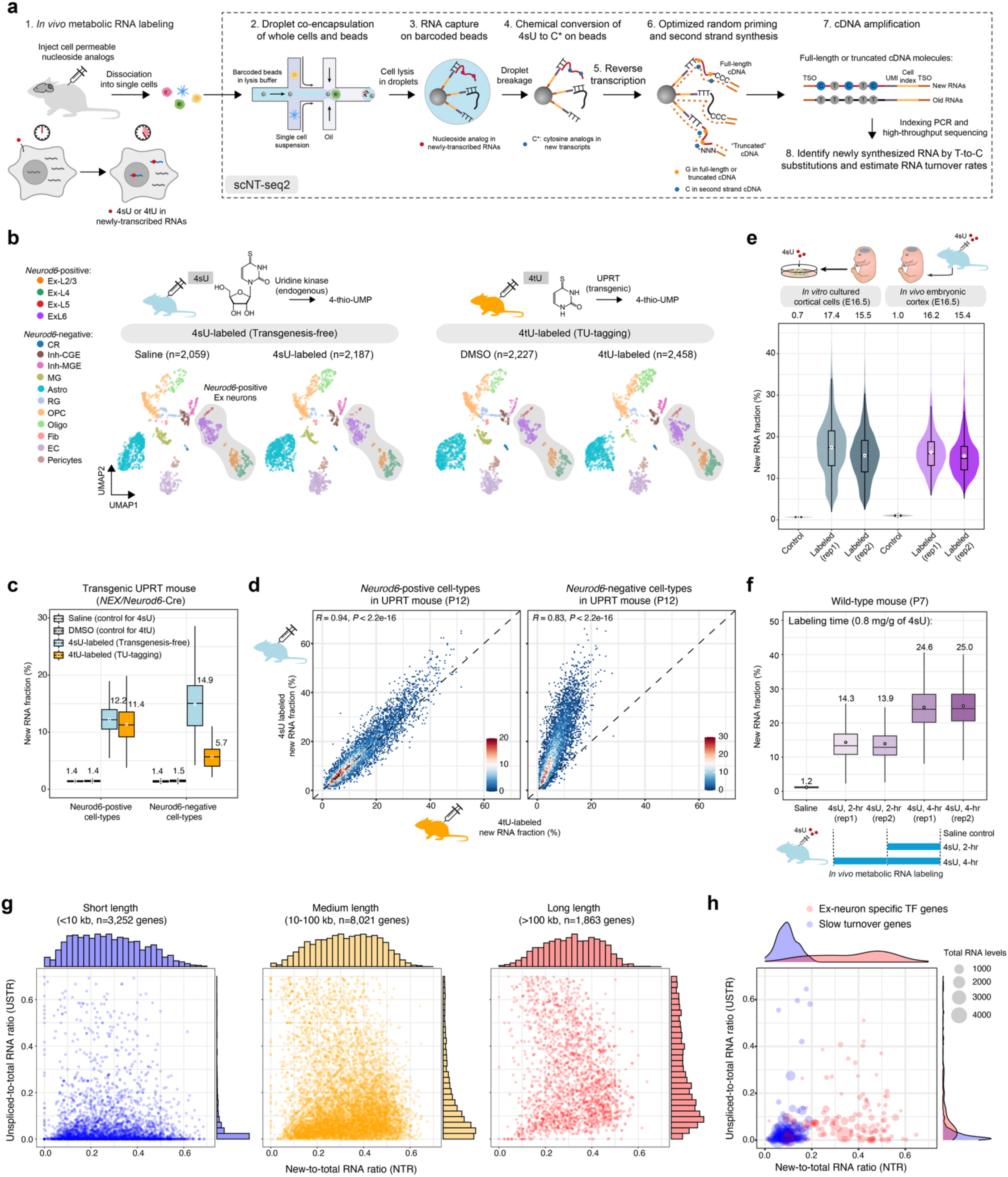
Benchmark and validation of *in vivo* transgenesis-free metabolic RNA labeling for cell-type specific RNA turnover analysis in mouse brains. **a.** Schematic depiction of the integrated workflow of *in vivo* metabolic RNA labeling followed by scNT-seq2. 4sU, 4-thiouridine; 4tU, 4-thiouracil; TSO, template switch oligo. **b.** UMAP visualization of cortical cells from four intraperitoneally injected *UPRT* transgenic mice colored by annotated cell types. DMSO injection serves as the control for the 4tU-labeled mouse, while saline injection is the control for the 4sU-labeled mouse. The cell number for each sample is indicated on the top. 4sU, 4-thiouridine; 4tU, 4-thiouracil; UPRT, Uracil phosphoribosyltransferase; 4-thio-UMP, 4-thiouridine monophosphate. CR, Cajal-Retzius cells; Ex-L2/3/4/5/6, cortical layer specific excitatory neurons; Inh-CGE, caudal ganglionic eminence (CGE)-derived interneurons; Inh-MGE, medial ganglionic eminence (MGE)-derived interneurons; MG, microglia; Astro, astrocytes; RG, radial glial cells; OPC, oligodendrocyte precursor cells; Oligo, oligodendrocytes; Fib, meningeal fibroblast; EC, endothelial cells. **c.** Box plot showing the 4sU-or 4tU-labeled new RNA fraction per cell for both *Neurod6*-positive and-negative cell-types (as defined in Fig. 1b). The boxes show the median (center line) and interquartile range (IQR, spanning the 25th to 75th percentiles). Whiskers extend to 1.5 times the IQR. The mean is shown as a white dot, with its value indicated above the box. **d.** Scatterplots comparing transcriptome-wide gene-level new RNA fractions between 4sU-(y-axis) and 4tU-labeled (x-axis) UPRT mice (P12 cortex) in Neurod6/NEX-cre targeted (*Neurod6*-positive cortical Ex neurons in the left panel) or non-targeted (*Neurod6*-negative cortical cells in the right panel) cell-types. **e.** Violin plot illustrating the proportion of 4sU-labeled new transcripts per cell from *in vivo* labeled embryonic day 16.5 (E16.5) mouse cortical tissues compared to *in vitro* labeled cultured E16.5 cortical cells ^1^. See ‘Data visualization’ in the Methods for definitions of violin plot and box plot elements. Mean new transcript fracctions for each sample is presented by a white dot, and the value is displayed above. Cell numbers for the conditions: control (media for *in vitro* culture): 1,999; *in vitro* labeled (rep1): 2,450; *in vitro* labeled (rep2): 4,295; control (saline injection for *in vivo* labeling): 3,129; *in vivo* labeled (rep1): 3,823; *in vivo* labeled (rep2): 3,305. **f.** Box plot (top panel) showing the 4sU-labeled new transcript fraction per cell for wild-type postnatal day 7 (P7) mouse cortical samples in labeling time evaluation experiments. A schematic depiction of 4sU labeling time experiment is shown (bottom panel). Cell numbers for the samples: saline control, 2,340; 4sU-2hr-rep1, 3,413; 4sU-2hr-rep2, 4,384; 4sU-4hr-rep1, 2,652; 4sU-4hr-rep2, 3,617. **g.** Scatter plots showing NTRs (x-axis, from RNA labeling) and USTRs (y-axis, from RNA splicing) for gene groups with distinct genes lengths (short – blue, <10 kb; medium – yellow, 10-100 kb; long – red, >100 kb) in cortical Ex neurons. Only expressed genes (transcript counts>5, CPM) are shown, and the number of genes for each group is indicated. The marginal histogram plots depict the distributions of NTRs (x-axis) and USTRs (y-axis) for each gene group. **h.** Scatter plots showing new-to-total RNA ratio (NTR derived from RNA labeling) and unspliced-to-total RNA ratio (USTR from RNA splicing) for cell-type specific TFs (red, n=124) and broadly expressed slow turnover genes (blue, n=269) in cortical Ex neurons. The size of circles represent total RNA levels for each gene (CPM: counts per million reads). The marginal density plots depict the distributions of NTRs (x-axis) and UTRs (y-axis) for two gene groups.

First, using the ‘TU tagging’ ^23,24,30^ as a benchmark, we evaluated the *in vivo* RNA labeling efficiency with 4sU in the brain. We generated “Ex-UPRT” mice by crossing a *Nex/Neurod6-Cre* driver ^31,32^ with a conditional *UPRT* line (*CA>GFPstop>UPRT*) ^23^, achieving *Neurod6-*positive excitatory (Ex) neuron-specific *Uprt* expression. “Ex-UPRT mice” at postnatal day 12 (P12) were injected intraperitoneally with 4tU or 4sU at equimolar doses ^33^, and scNT-seq2 analysis identified 15 major cortical cell-types (**Fig. 1b** and **Extended Data Fig. 1d**) and confirmed *Uprt* transgene expression restricted to *Neurod6*-expressing Ex neurons (**Extended Data Fig. 1e**). Both 4sU and 4tU labeling minimally altered cell-type specific transcriptomic states, as shown by closely matched distributions of cortical cell-types in treated and control mice (**Fig. 1b** and **Extended Data Fig. 1d**). Quantification of labeled new RNA fractions indicated robust labeling in *Neurond6*-postive Ex neurons with both 4sU and 4tU (**Fig. 1c**). Transcriptome-wide gene level analysis from 4sU and 4tU labeling were highly concordant in cortical Ex neurons (left in **Fig. 1d**). Notably, 4sU also efficiently labeled new RNAs in *Neurod6*-negative cell types (**Fig. 1c**), independent of UPRT expression, whereas 4tU labeling in these cell-types was substantially lower (**Fig. 1c-d**), likely due to low endogenous UPRT-like enzymatic activity ^34^. Thus, 4sU enables efficient, broad *in vivo* RNA labeling in the brain, matching the labeling efficiency of TU-tagging.

Second, we further evaluated transgenesis-free RNA labeling with 4sU in wild-type animals. Time-pregnant females were injected 4sU, and embryonic day 16.5 (E16.5) embryonic cortices were analyzed alongside primary E16.5 cortical cells labeled *in vitro.* This direct comparison showed similar increases in 4sU-labeled new RNA fractions (*in vitro*: 16.5%, *in vivo*: 15.8%) compared to saline controls (<1.0%), with comparable cell-to-cell variability (**Fig. 1e**), likely reflecting intrinsic differences in RNA metabolism at individual cell levels ^35^. Transcriptome-wide comparisons of gene-level new-to-total RNA ratios (NTRs) showed high concordance between *in vitro* and *in vivo* labeling (**Extended Data Fig. 1f**), demonstrating that 4sU efficiently crosses both fetal-maternal and blood-brain barriers to label new RNAs in developing embryonic brains.

Third, we systematically evaluated the effects of 4sU labeling duration and dosage on RNA labeling across 26,640 cortical cells from nine wild-type postnatal day 7 (P7) mice (**Extended Data Fig. 2a-b**). Consistent with “Ex-UPRT” mice, 4sU labeling minimally affected cell-type composition (**Extended Data Fig. 2c**). Liquid chromatography-tandem mass spectrometry analysis showed that cortical 4sU levels rose for two hours post-injection and stabilized thereafter (**Extended Data Fig. 2d**). Labeled RNA fractions increased in a nearly linear fashion with time (2-hr: 14.1%, 4-hr: 24.8% in **Fig. 1f**), consistent across biological replicates and cortical cell-types (**Extended Data Fig. 2e**). Higher 4sU dose further increased labeled new RNA fractions, plateauing at ∼1 mg/g (**Extended Data Fig. 2f**). Importantly, 4sU labeling did not affect scNT-seq2 performance (mean mapping rates – saline: 79.8%, 4sU: 79.4%; mean library complexity [detected at 50,000 reads per cell] – saline: 5,826 genes and 19,912 transcripts, 4sU: 5,683 genes and 19,246 transcripts), and aggregated single-cell new and old transcriptomic profiles showed high reproducibility across biological replicates (**Extended Data Fig. 2g**).

Lastly, we assessed the impact of 4sU labeling on global gene expression using an established approach ^36^. Transgenesis-free *in vivo* 4sU labeling showed minimal transcriptional perturbation across developmental stages (E16.5, P7 and P12), labeling durations (2h vs 4h), and 4sU dosage (**Supplementary Fig. 2a**), compared to *in vitro* 4sU or *in vivo* 4tU (TU-tagging) labeling. Spliced-to-total RNA ratios were highly concordant between 4sU-labeled and control samples, indicating little effect of *in vivo* labeling on mRNA splicing (**Supplementary Fig. 2b**). Notably, the unspliced-to-total RNA ratio (USTR) provides a labeling-free estimate of global RNA processing and turnover rates ^37^. To validate *in vivo* 4sU labeling for estimating global RNA processing and turnover, we compared NTRs (labeling-based) with USTRs (splicing-based) across cortical cell-types in two P7 mice. Significant correlations between NTRs and USTRs across replicates (**Supplementary Fig. 2c**) suggests that labeling-based measurements reflect intrinsic cell-type-specific RNA processing and turnover rather than local 4sU variability under saturating conditions.

Together, these results demonstrate that integrating 4sU-based RNA labeling with droplet-based scNT-seq2 enables quantitative single-cell profiling of new and old transcriptomes across diverse cell-types in embryonic and postnatal mouse brains. Compared to TU-tagging, this transgenesis-free approach achieves similar *in vivo* labeling efficiency with broad applicability and minimal cytotoxicity.

### Transcriptome-wide RNA turnover analysis across cortical cell-types

To examine gene-level RNA turnover across cortical cell types, we identified cell-type-specific genes (**Supplementary Fig. 3a**) and determined RNA turnover rates for different gene groups by computing their NTRs ^16,38^. This analysis confirmed that cell-type-specific transcription factors (TFs) exhibited faster turnover than housekeeping genes (in Ex neurons – TFs: 0.35, housekeeping genes: 0.095; **Supplementary Fig. 3b**). Unlike splicing-based estimates (USTRs), which are confounded by gene length and structure (y-axis in **Fig. 1g** and **Supplementary Fig. 3c**), RNA labeling-based analysis (NTRs) quantitatively capture turnover differences across gene groups with distinct functions (e.g. x-axis in **Fig. 1h** and **Supplementary Fig. 3d**). This is likely because shorter or intron less genes are sparsely covered in splicing-based analysis.

Gene set enrichment analysis further showed that fast-turnover genes are enriched for cell-type-specific TFs (**Supplementary Fig. 4a**), such as *Neurod6* (NTR: 0.459) and *Rorb* (NTR: 0.456) in Ex neurons, while broadly expressed genes like *Eef2* (NTR: 0.092) and *Eif3k* (NTR: 0.102) show slower turnover (**Supplementary Fig. 4b-c**). Notably, non-TF cell-type-specific genes exhibit diverse RNA turnover linked to different functions (**Supplementary Fig. 3b** and **4d**), highlighting that RNA turnover regulation is optimized for gene function and cell identity.

### Development and validation of spatial NT-seq

Current sequencing-based spatial transcriptomics (ST) methods measure total RNA without distinguishing newly transcribed from pre-existing RNAs ^39–41^. A recent approach combined 5-EU labeling with *in situ* sequencing of a pre-defined gene panel to assess RNA turnover in cultured cells ^17^, but lacked transcriptome-wide coverage and required pulse-chase labeling involving multi-timepoint measurements to infer population-level kinetics.

To enable transcriptome-wide spatial RNA turnover analysis *in vivo*, we leveraged SLAM chemistry ^42^, which modifies 4sU with iodoacetamide (IAA) to induce T-to-C substitutions in new RNAs during reverse transcription (**Fig. 2a**). As the SLAM reaction requires organic solvents, we reasoned that it could be adapted for *in situ* chemical conversion in methanol-fixed cells or tissue sections and made compatible with various sequencing-based ST platforms. Previous applications of SLAM reaction reported relatively low conversion efficiency (∼0.5-1% T-to-C substitutions) in cultured cells ^20,43^. Using on-bead TimeLapse chemistry used in scNT-seq2 as a benchmark, we systematically optimized key parameters of *in situ* SLAM reaction (e.g. IAA concentration, reaction temperature/time) (**Supplementary Fig. 5a-c**), achieving markedly higher conversion rates (*in situ* SLAM vs. on-bead Timelapse based 4sU conversion rates - 5.35% vs. 5.71%) without compromising performance (**Supplementary Fig. 5c-e**).

**Fig. 2.**
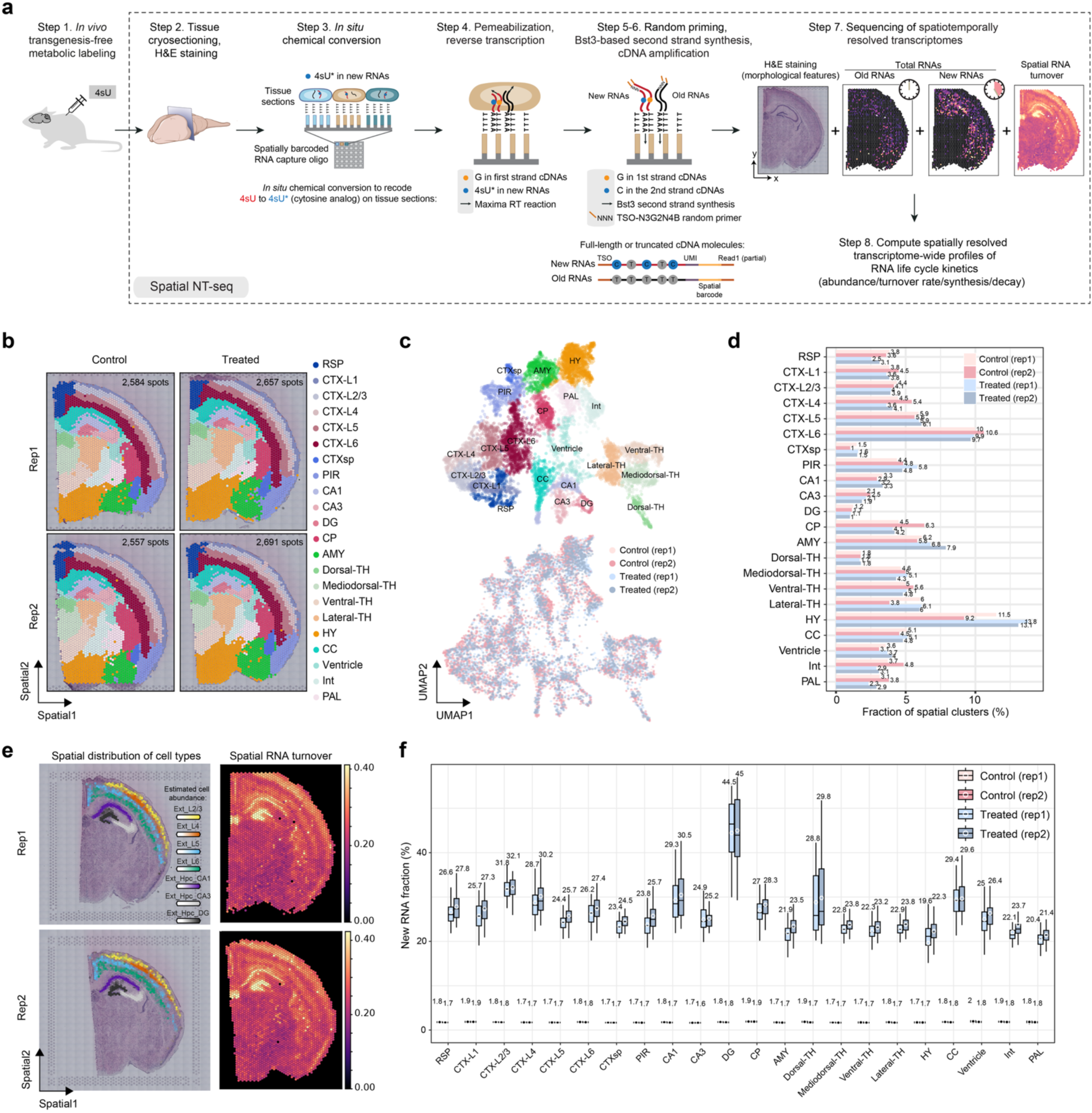
Development and validation of *in vivo* spatial NT-seq for spatially resolved profiling of transcriptome-wide RNA turnover. **a.** Schematic diagram of Spatial NT-seq. Specifically, we developed *in situ* chemical recoding within tissue sections via SLAM reaction, which covalently attaches a carboxyamidomethyl group from iodoacetamide (IAA) to the thiol group in 4sU. This modified base (4sU*) is then transformed into a cytosine analog to induce T-to-C substitutions in new RNAs during subsequent reverse transcription reaction. Spatial patterns of “new” and “old” RNAs for the *Bcl11a* gene are shown as an example. Some components of the diagram were adapted from previous papers ^2,3^. **b.** Transcriptome based annotation of spatial clusters within four coronal sections from the P7 wild-type mouse brain. Treated: sections subjected to the *in situ* chemical conversion; Control: sections without conversion. Numbers of spatial spots per section: control-rep1, 2,584; control-rep2, 2,557; treated-rep1, 2,657; treated-rep2, 2,691. Abbreviations for brain regi^4,5^ons: RSP, retrosplenial area; CTX, cortex; CTXsp, cortical subplate; PIR, piriform area; CA, cornu ammonis; DG, dentate gyrus; CP, caudoputamen; AMY, amygdalar nucleus; TH, thalamus; HY, hypothalamus; CC, corpus callosum; Int, internal capsule; PAL, pallidum. **c.** UMAP visualization of spatial spots from the four brain sections, colored by annotated spatially resolved brain regions (top, corresponding to **b**) or by sample names (bottom). **d.** Relative composition of spatial spots assigned to each spatial region within the four brain sections. **e.** Visualization of cell abundance score (indicated by color intensity) for seven representative excitatory (Ext) neuronal subtypes within cortical and hippocampal (Hpc) regions across two treated tissue sections in Fig. 2b (left panel). Spatial mapping of global RNA turnover by corrected NTRs per spot (right panel). **f.** Box plot illustrating the corrected new RNA fraction per spot over 22 brain regions within each section. Mean fraction is represented as a white dot, with its value displayed on the top.

To enable spatially resolved co-mapping of RNA abundance and turnover in the brain, we developed spatial NT-Seq (**Fig. 2a**). This method integrates transgenesis-free *in vivo* 4sU labeling, the preparation of fresh frozen tissue sections, methanol fixation, and histological staining. The critical step of chemically recoding 4sU-labeled new mRNAs is then performed directly through *in situ* SLAM reaction within tissue sections on Visium spatial capture slides, a widely adopted sequencing-based ST platform that contain spatially barcoded mRNA capture oligonucleotides ^44^. Following optimized tissue permeabilization, modified RT and second strand synthesis reactions (see methods) were performed to capture newly both synthesized (“new”) and pre-existing (“old”) transcriptomes within the same spatial spot.

To benchmark spatial NT-seq, we applied it to metabolically labeled P7 mouse brains (**Extended Data Fig. 3a**). Compared to control (i.e. standard Visium), spatial NT-seq showed comparable library complexity and alignment rates (**Extended Data Fig. 3b-c**). Clustering of 10,489 spatial spots from control and treated coronal brain sections identified 22 distinct spatial clusters (**Fig. 2b-c**), which were annotated using the Allen Common Coordinate Framework (CCF) reference ^6^ and further validated by *Cell2location*-based integration with reference whole brain single-nucleus RNA-seq (snRNA-seq) data ^45^ (**Supplementary Fig. 6a-c**). Importantly, spatial cluster distributions (**Fig. 2b-c** and **Supplementary Fig. 6b**) and proportions (**Fig. 2d**) were highly concordant between control and treated groups, with minimal impact of labeling on global gene expression (**Extended Data Fig. 3d**). Notably, spatial NT-seq achieved a 14.6-fold increase in T-to-C conversion compared to control (**Extended Data Fig. 3e**). Thus, this method enables efficient *in situ* chemical conversion while preserving tissue architecture, spatial resolution, and detection sensitivity, allowing spatial co-profiling of RNA abundance and turnover in the brain.

### Spatial profiling of RNA turnover in the mouse brain

To map spatial RNA turnover *in vivo*, we quantified labeled new RNA fractions across P7 mouse brain regions (**Fig. 2e-f**). Most regions showed similar new RNA fractions (mean±s.d. = 25.7±3.1% across 21 domains excluding DG), but specific regions (DG: 44.8%, CTX_L2/3: 32.0%, CA1: 29.9%) exhibited substantially higher levels (**Fig. 2e-f**). Spatial NT-seq analysis of postnatal day 56 (P56) adult mouse brains further confirmed elevated RNA turnover in DG and CA1 across animals (**Extended Data Fig. 4a-b**). Notably, these differences are unlikely to be driven by local 4sU availability, as vascular density is comparable (**Extended Data Fig. 4c**) or even lower in DG and CA1 relative to other regions, based on brain-wide quantitative mapping of the vascular network including arteries, capillaries and veins ^46^. Moreover, we computed the probability of T-to-C substitutions in newly synthesized transcripts, which primarily reflects 4sU incorporation efficiency ^47^, across all P7 brain regions. This incorporation rate depends on the relative local concentrations of triphosphorylated 4sU and U available for incorporation into nascent RNAs ^47^. Importantly, 4sU incorporation efficiency is relatively comparable across most brain regions (**Extended Data Fig. 4d**). Thus, the observed differences in RNA turnover across brain regions largely reflect intrinsic regulation of RNA metabolism.

Next, we investigated potential mechanisms underlying elevated RNA turnover in specific brain regions by examining the expression level of genes related to global regulation of RNA synthesis and stability. This analysis revealed that *Uck1*, a uridine/cytidine kinase in the pyrimidine salvage pathway ^48^, is expressed at significantly higher levels in the DG and CA1 compared to other brain regions (**Supplementary Fig. 7**). This enzyme catalyzes the phosphorylation of uridine and related analogs (e.g. 4sU), a rate limiting step for recycling pyrimidine nucleosides and activation of 4sU for new RNA synthesis ^49^. Notably, key components of major RNA degradation pathways, including *Upf1* (essential for nonsense-mediated mRNA decay (NMD) pathway), *Eif4a3* (a exon junction complex (EJC) factor required for NMD activity) ^7^, *Mettl3* (an enzyme in the RNA *N*^6^-methyladenosine (m^6^A) modification pathway), *Ythdf2* (a m6A reader promoting RNA decay) ^50,51^, *Cnot1* (a scaffold protein of the CCR4-NOT deadenylase complex) ^52^, and *Ago2* and *Dicer1* (key regulators of microRNA biogenesis and processing) ^53^, were also consistently up-regulated in the DG relative to other brain regions (**Supplementary Fig. 7**). Thus, enhanced pyrimidine nucleoside salvage and increased RNA degradation likely contribute to the elevated RNA turnover observed in the DG.

Finally, consistent with *in vivo* single-cell analysis, cell-type-specific TFs exhibited substantially higher turnover rates than housekeeping genes (**Extended Data Fig. 3f**). Brain-region-specific TFs such as *Rorb* (enriched in CTX-L4/TH) and *Bcl11b* (enriched in CTX-L6/CP/PIR/CA1/DG) showed markedly elevated turnover rates compared to housekeeping genes like *Actb* and *Mapt* (left panels, **Extended Data Fig. 3g**). In contrast, splicing-based analysis of these genes failed to accurately capture spatial profiles of new (unspliced reads) and old (spliced reads) RNAs (right panels, **Extended Data Fig. 3g**), likely due to sparse coverage and/or gene structure limitations. Together, these results establish *in vivo* spatial NT-seq as a robust approach for spatial profiling of RNA turnover and uncovering regional regulation of RNA metabolism.

### Spatiotemporally resolved profiling of ECS-induced RNA dynamics

Elevated RNA turnover in regions such as the DG may help maintain cellular plasticity, enabling rapid responses to learning, memory formation, and external stimuli ^4,28^. To investigate how spatially distinct RNA turnover contributes to activity-dependent gene regulation, we applied spatial NT-seq to profile RNA dynamics in response to synchronous neuronal activation induced by ECS, a procedure currently used to treat patients with severe mood and psychotic disorders, including depression, bipolar disorder, and schizophrenia ^54^. Despite its clinical efficacy, the molecular mechanisms underlying ECS remain incompletely understood ^55^. ECS induces widespread neural activity and induces multi-phase expression of activity-regulated genes (ARGs) ^56^, particularly in the hippocampus, with lasting effects on synaptic plasticity, neurogenesis, and neuronal morphology ^57,58^. While previous studies identified cell-type-specific gene expression changes post-ECS ^59,60^, they could not disentangle the distinct contributions from RNA synthesis and decay or capture spatially resolved RNA turnover across brain regions.

To spatiotemporally profile the ECS-induced ARG dynamics *in vivo*, we performed RNA labeling in two groups of adult male mice (P56) and conducted spatial NT-seq at 0 (control), 30-, and 120-min post-stimulation (**Fig. 3a**). Specifically, ECS was administered using a transient electrical stimulus (1.0 s total duration, see Methods) ^60^. Integrated analysis of 44,434 high-quality spatial spots across 16 brain sections identified 18 major spatial clusters based on whole-transcriptome similarity (**Fig. 3b** and **Extended Data Fig. 5a-c**). For instance, the CTX-L2/3 region was enriched for upper-layer excitatory neurons (Ext_L2/3, red), whereas CA1 and DG regions were associated with hippocampal excitatory CA1 (Ext_Hpc_CA1, purple) and DG (Ext_Hpc_DG1/2, black) neurons, respectively (**Extended Data Fig. 5b**). Quantification of labeled new RNA fractions confirmed robust *in vivo* labeling and high chemical conversion efficiency across time points (**Extended Data Fig. 4b**), with DG consistently exhibiting elevated RNA turnover compared to other regions, mirroring observations in P7 mouse brains.

**Fig. 3.**
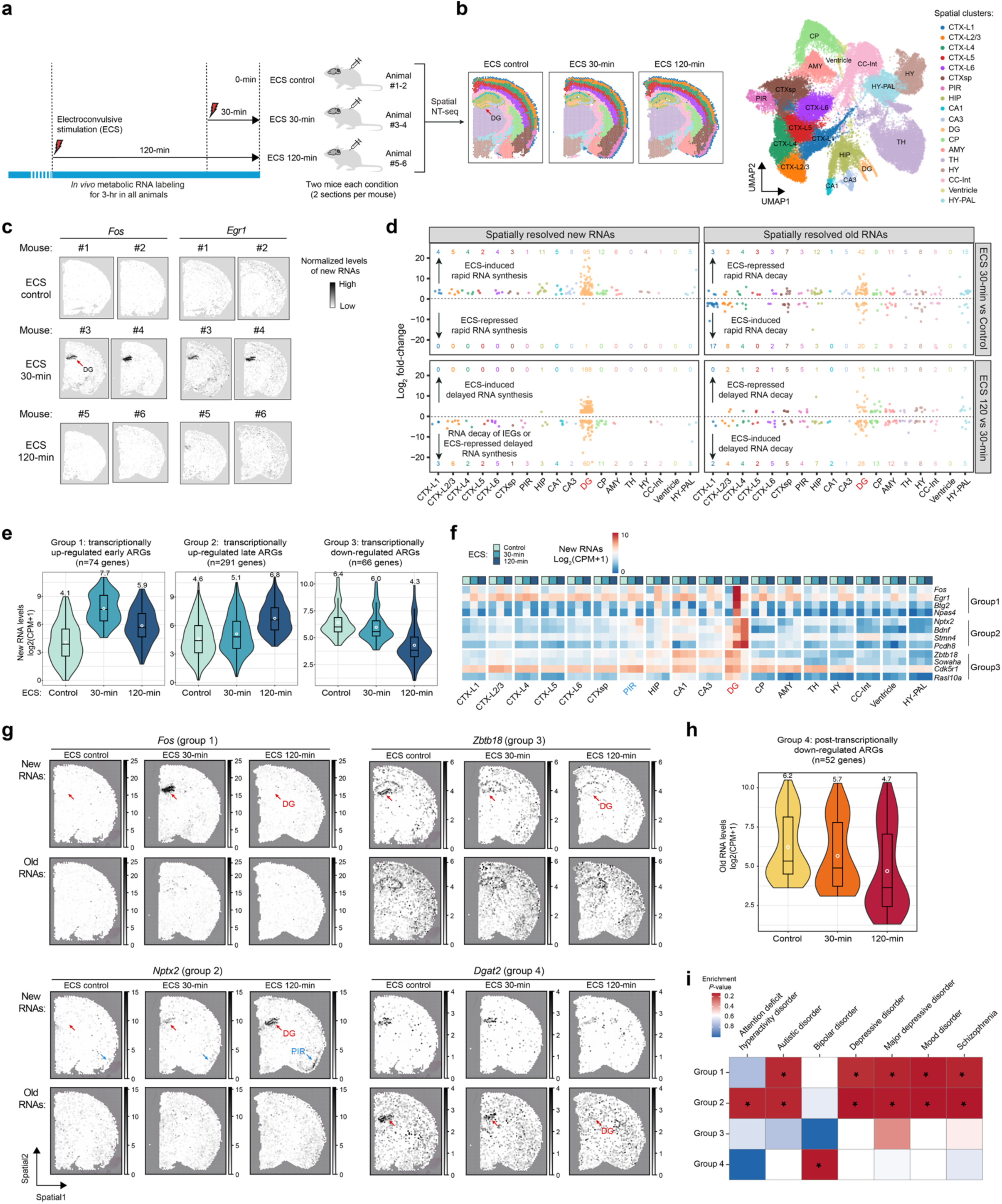
Spatiotemporally resolved profiling of electroconvulsive stimulation (ECS) induced *in vivo* RNA dynamics. **a.** Experimental design for spatiotemporally resolved analysis of RNA dynamics in mouse brains following acute ECS using spatial NT-seq. All mice were subjected to *in vivo* transgenesis-free metabolic RNA labeling (i.e. a total of 3-hour 4sU labeling). For each of three time points: non-ECS (control), 30min post-ECS (ECS-30min), and 120min post-ECS (ECS-120min), samples were collected from two adult mice per time point as biological replicates and two sections per mouse as technical replicates. **b.** Shown on the right are three representative sections with spatial annotations and UMAP visualization of all spatial spots (n= 44,344) from 16 brain sections (see **Supplementary Figure 14a**), colored by their annotated spatial regions. Region abbreviations: CTX, cortex; CTXsp, cortical subplate; PIR, piriform area; HIP, hippocampus; CA, cornu ammonis; DG, dentate gyrus; CP, caudoputamen; AMY, amygdalar nucleus; TH, thalamus; HY, hypothalamus; CC, corpus callosum; Int, internal capsule; PAL, pallidum. **c.** Panels displaying the normalized spatial expression of newly synthesized RNAs (in CP10K) for two early ARGs that encode TFs, *Fos* and *Egr1*, from two animals per time points. Red arrows indicate the DG region. **d.** Scatter plots representing the number and log-scaled fold change (FC) of differentially regulated new (left) or old (right) RNAs (adjusted P-value <0.001 and FC>4) in response to ECS across 18 brain regions for pair-wise comparisons: ECS-30min vs. control (top panels) and ECS-120min vs. ECS-30min (bottom panels). Distinct regulatory strategies through either transcriptional (via changes of new RNAs) or post-transcriptional (primarily via changes of old RNAs) mechanisms are indicated. Note that these mechanisms are not mutually exclusive, but can be synergistically or antagonistically deployed by cells to achieve desired dynamic changes at total RNA levels. Moreover, we cannot exclude the possibility that ECS-induced new RNAs can also be regulated by RNA decay mechanisms. **e.** Volin plots illustrating ECS-induced dynamic changes of three groups of transcriptionally regulated ARGs in DG. The plots showing normalized expression levels (log-scaled (CPM+1)) of new RNAs within the DG across three time points: control, ECS-30min, and ECS-120min. Open white circles indicate the mean value of new RNA levels for each group of ARGs at a given time point. **f.** Heat maps illustrating the ECS-induced dynamics of new RNAs for 12 select transcriptionally regulated ARGs (in DG) from three groups across 18 brain regions. **g.** Panels displaying the normalized spatial expression (in CP10K) of new (upper panel) and old (lower panel) RNAs for four representative genes from each ARG group using a representative spatial NT-seq dataset (section #2 from mouse #2/4/6). Notably, red and blue arrows in the *Nptx2* panel indicate the DG and PIR regions, respectively. **h.** Violin plots illustrating the ECS-induced dynamics of post-transcriptionally down-regulated ARGs. The plots showing normalized expression levels (log-scaled (CPM+1)) of old RNAs within the DG across three time points: control, ECS-30min, and ECS-120min. Open white circles indicate the mean value of old RNA levels for each group of ARGs at a given time point. **i.** Gene set enrichment analysis of four groups of ARGs within curated disease gene lists for various brain disorders. *, *P-*value < 0.05.

**Fig. 4.**
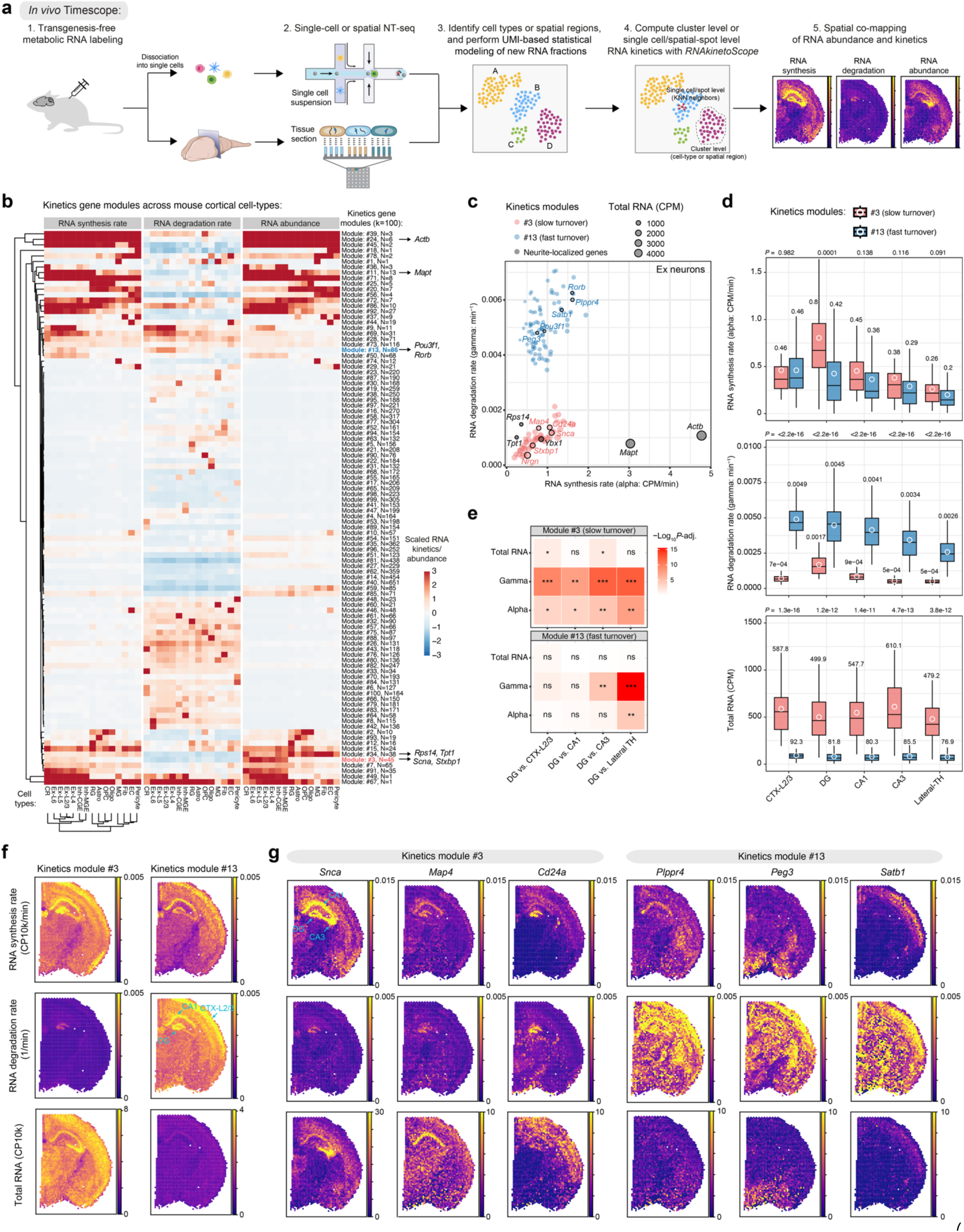
*In vivo* profiling of single-cell and spatial RNA kinetics landscapes in the mouse brain. **a.** Schematic illustrating “*in vivo* Timescope”, an integrated experimental and computational strategy for quantifying RNA kinetics parameters at across cellular or spatial contexts within complex tissues such as mouse brains. **b.** Clustered heat maps showing scaled RNA synthesis rate (alpha, left), degra^6^dation rate (gamma, middle), and total RNA abundance (right) for 100 kinetics gene modules across 15 cortical cell types. The number of genes (*n*) in each module is shown. Representative genes for specific kinetics gene modules (#3: *Scna*, *Stxbp1*; #13: *Pou3f1*, *Rorb*) and neurite-localized/enriched genes (*Actb*, *Mapt*, *Rps14*, and *Tpt1*) were indicated by arrows. **c.** Scatter plot showing RNA synthesis (x-axis: CPM/min) versus degradation (y-axis: min^-1^) rates in cortical Ex neurons for genes in kinetics modules #3 (n=45 genes in red: *Stxbp1*, *Nrgn*, *Map4*, *Snca*, and *Cd24a*) and #13 (n=86 genes in blue: *Pou3f1*, *Plppr4*, *Peg3*, *Satb1*, and *Rorb*). Genes are color-coded by kinetics modules, and their total RNA level is represented by dot size. Five representative neurite-localized/enriched genes (black: *Actb*, *Mapt*, *Rps14*, *Tpt1*, and *Ybx1*) are also highlighted. **d.** Box plots comparing spatial region–specific RNA synthesis rates (alpha, top), degradation rates (gamma, middle), and total RNA abundance (bottom) between two neuron-specific kinetics modules: module #3 (red, n = 45 genes; slower turnover) and module #13 (blue, n = 86 genes; faster turnover), across five representative brain regions. The mean is indicated by a white dot, with the exact value shown above each box. *P*-values from two-sided *t*-test comparing modules #3 and #13 are reported above each panel. **e.** Heatmaps showring statistical analysis of spatial region-specific total RNA abundance, RNA degradation rates (gamma), and RNA synthesis rates (alpha) between DG vesus representative non-DG brain regions of P7 mouse brains for genes in kinetics modules #3 and #13 genes. In the heatmap, significance is indicated as follows: ***, *P* < 1×10^-8^; *\*\*, P* < 1×10^-4^; *, *P* < 0.05; ns, non-significant. **f.** Spatial feature plots displaying aggregated RNA synthesis rate/alpha (top), degradation rate/gamma (middle), and total RNA abundance (bottom) for all genes in kinetics modules #3 (n=45 genes; slower turnover) and #13 (n=86 genes; faster turnover). **g.** Spatial feature plots highlighting RNA synthesis rate/alpha (top), degradation rate/gamma (middle), and total RNA abundance (bottom) for six representative genes: *Scna*, *Map4* and *Cd24a* from module #3, along with *Plppr4*, *Peg3* and *Satb1* from module #13.

To assess the accuracy of spatial NT-seq in analyzing *in vivo* ARG dynamics, we examined immediate early genes (IEGs) such as *Fos* and *Egr1*, which are transiently activated following ECS ^59–61^. Rapid activation of *Fos* and *Egr1* was specifically and reproducibly detected in metabolically labelled new RNAs, but not in old RNAs, within the DG region across biological replicates (**Fig. 3c** and **Extended Data Fig. 6a**). Labeling-based RNA velocity analysis ^21,62^ revealed early ARG activation within 30-min post-ECS (**Extended Data Fig. 6b**), demonstrating the high temporal resolution of spatial NT-seq. In contrast, splicing-based analysis failed to resolve the spatial dynamics of new (unspliced) and old (spliced) RNAs for IEGs (**Supplementary Fig. 8a**). Consistently, labeling-based new RNA analysis substantially outperformed splicing-based approaches in capturing IEG activation signatures ^63^ across brain regions (**Supplementary Fig. 8b–e**).

While standard snRNA-seq has shown that ECS can up-or down-regulate gene expression in hippocampal neurons ^59^, it cannot distinguish between transcriptional and post-transcriptional contributions to ARG dynamics. To address this gap, we jointly analyzed both newly synthesized and pre-existing transcriptomes within the same spatial spot across brain regions, enabling mechanistic dissection of transcriptional (e.g. new RNA induction) and post-transcriptional (e.g. old RNA decay) regulations. This analysis revealed that the DG was the most responsive to ECS, exhibiting significant changes in both new and old RNAs levels, independent of the filtering thresholds (fold change (FC)>4 in **Fig. 3d** or FC>2 in **Extended Data Fig. 5d**; adjusted *P*<0.001 for both). Other brain regions showed more modest responses, particularly in old RNAs, suggesting more prevalent post-transcriptional regulation (**Fig. 3d** and **Extended Data Fig. 5d**).

Next, to elucidate regulatory strategies underlying ARG dynamics within the DG region, we classified genes based on their ECS-induced temporal expression patterns of either new or old RNAs. Analysis of new transcriptomes identified two groups of transcriptionally activated ARGs in the DG following ECS. Group 1 comprises 74 early-response ARGs (adjusted *P*<0.001, FC>4; **Supplementary Table S1**), including IEGs such as *Fos*, *Egr1* and *Btg2* (**Fig. 3e-g** and **Supplementary Fig. 9a-b**), which peak within 30-min post-ECS. Group 2 includes 291 late-response ARGs, such as *Nptx2*, *Bdnf* and *Stmn4* (**Fig. 3e-g** and **Supplementary Fig. 9a-b**). Early ARGs in group 1, often encoding TFs that drive the transcription activation of ARGs in group 2, are rapidly degraded in part through AU-rich element (ARE) directed deadenylation and mRNA decay ^8,64,65^. Consistently, gene ontology enrichment analysis linked group 1 to transcriptional regulation and Group 2 to synaptic functions (**Supplementary Fig. 9c**). Notably, transcriptional activation of *Nptx2* (Group 2) was also detected in the piriform area (PIR in **Fig. 3f-g**), suggesting a broader role of this late ARG.

Spatial co-profiling of new and old transcriptomes enables a more detailed dissection of ARG down-regulation mechanisms. Analysis of new RNA dynamics identified 66 genes (group 3) with progressively reduced levels of new RNAs post-ECS, suggesting transcriptional repression (e.g. *Zbtb18* and *Bhlhe22* in **Fig. 3e-g** and **Supplementary Fig. 10a-b**). Moreover, analysis of old RNA dynamics, focusing on down-regulation of old RNAs to avoid biases from incomplete capture of new RNAs, identified 52 genes (group 4) likely regulated via ECS-induced mRNA decay (e.g. *Dgat2* and *Sphkap* in **Fig. 3g-h** and **Supplementary Fig. 10a-b**). Notably, Groups 3 and 4 are not mutually exclusive as some genes (e.g. *Rasl10a* in **Supplementary Fig. 10b**) might be co-regulated through both transcriptional repression and post-transcriptional degradation. Conversely, other genes (e.g. *Cracdl* in **Supplementary Fig. 10b**) exhibited opposing effects between transcriptional activation (i.e. increase in new RNA) and post-transcriptional degradation (i.e. decrease in old RNAs), which may serve to buffer expression levels. These ECS-induced transcriptional and post-transcriptional regulatory effects are specific, as control genes showed no major changes in either new or old RNA levels (**Supplementary Fig. 10c**).

To enhance spatial resolution of ARG dynamics analysis, we applied a tailored *iStar* pipeline ^66^, which integrates spot-level spatial NT-seq data (∼1-10 cells per spatial spot in standard Visium platform) with high-resolution histological images from the same tissue section to reconstruct near single-cell resolution (spatial pixel size: 8×8 μm^2^) expression profiles of new and old RNAs (**Extended Data Fig. 6c**). Predicted near single-cell resolution expression dynamics of representative ARGs (group 1-4) closely matched spot-level data for both new and old RNAs (**Extended Data Fig. 6d**), demonstrating the potential of this approach to enable whole-transcriptome, time-resolved spatial analysis at high-resolution from spatial NT-seq data.

Finally, we assessed the disease relevance of ECS-induced ARGs by comparing each group to curated disease gene sets ^67^. Transcriptionally activated ARGs (Group 1 and 2) were significantly enriched for genes associated with major depressive disorder, mood disorder, and schizophrenia (**Fig. 3i**). In contrast, only Group 4 showed significant enrichment for bipolar disorder (**Fig. 3i**), suggesting a distinct role for post-transcriptional regulation in this disease. Several ECS-induced ARGs are also linked to neurological diseases beyond the mood disorders. For example, *Nptx2* overexpression (group 2; encoding a synaptic protein) in the hippocampus alleviates stress-induced anxiety-like behaviors, while its hippocampus-specific deletion exacerbates them ^68^. Mis-regulation of NPTX2 has also been implicated in TDP43-induced neurodegeneration ^69^. Similarly, down-regulated ARGs such as *Zbtb18* (group 3; encoding a neuronal TF) and *Dgat2* (group 4; encoding an endoplasmic reticulum/mitochondrion-associated protein) are associated with neurodevelopmental disorders ^70,71^ and inherited neuropathies ^72^. Together, these findings demonstrate the utility of high spatiotemporal resolution profiling of ARG dynamics by spatial NT-seq in uncovering the molecular mechanisms underlying therapeutic effects of ECS.

### *In vivo* RNA kinetics landscape profiling across brain cell types

RNA abundance reflects a balance between RNA synthesis and degradation, with similar steady-state levels potentially arising through distinct kinetic mechanisms. Moreover, mRNA stability may directly influence RNA localization to distant compartments in highly polarized cell-types such as neurons ^11^. However, *in vivo* RNA kinetics across diverse cell-types and brain regions remains largely unexplored.

We applied an optimized binomial mixture model to correct for incomplete 4sU labeling and background mutations, enabling more accurate quantification of new RNA fractions at the level of individual transcripts (linked by unique molecular identifiers, UMIs) ^21^. This statistical approach enables direct estimations of steady-state RNA synthesis and degradation rates ^16,21,62^, providing a robust framework for *in vivo* RNA kinetics profiling from single-cell and spatial NT-seq datasets (**Fig. 4a**). Prior simulation analysis ^73^ established that our ‘single-pulse’ RNA labeling strategy allows quantitative analysis of RNA turnover kinetics ^16^. Using the P7 cortical single-cell dataset, we systematically benchmarked the performance of our transcriptome-wide RNA kinetics analysis across 15 cortical cell types, observing high concordance between biological replicates for cell-type specific RNA abundance, synthesis, and degradation rates (**Supplementary Fig. 11a-b**). Furthermore, *in vivo* RNA half-lives computed for these cell-types inversely correlated with RNA turnover rates (**Supplementary Fig. 11c**), consistent with previous studies in cultured cells ^3,16,21,38^.

Our *in vivo* RNA kinetics analysis reveals an average RNA half-life of 4.6-hr across diverse cortical cell types (**Supplementary Fig. 11d**), which is highly consistent with averaged previous *in vitro* estimates (∼4.7-hr) obtained using single-pulse or pulse-chase metabolic RNA labeling methods (e.g. mouse ESCs: 3.9-hr ^42^, mouse fibroblasts: 4.6-hr ^74^, and mouse primary cortical neurons: soma-enriched mRNAs – 3.7-hr and neurite-enriched mRNAs – 5.6-hr ^11^). Notably, *in vivo* brain RNA half-lives spanned more than two orders of magnitude across the transcriptome of various cortical cell-types (ranging from <1-hr to>100-hr; **Supplementary Fig. 11d**). Transcriptome-wide RNA half-life estimates were highly consistent between 2-hr and 4-hr labeling experiments (*R*=0.95, *P*<2.2×10^-16^ in **Supplementary Fig. 11e**), suggesting that our *in vivo* RNA kinetics measurements are robust to 4sU labeling duration. As expected, accurate estimation of RNA degradation rates from a single short labeling (<4-hr) remains challenging for transcripts with long half-lives ^73^. Simulation analysis further revealed that the confidence intervals of estimated, cell-type specific RNA half-lives depend on both gene expression levels and intrinsic RNA stability, and genes with low expression and longer half-lives exhibit greater uncertainty (**Supplementary Fig. 11f**).

Transcriptome-wide RNA kinetics analysis of spatial NT-seq datasets also demonstrated high reproducibility between replicates for RNA abundance, synthesis, and degradation rates across 22 brain regions (**Supplementary Fig. 12a-c**). Moreover, RNA half-life profiles derived from *in vivo* single-cell and spatial NT-seq data showed high concordance in matched cortical Ex neuron subtypes (**Supplementary Fig. 12d**).

### *In vivo* mapping of RNA turnover kinetics across brain cell-types in the cortex

To investigate how *in vivo* RNA kinetics regulation shapes expression of genes with different functions, we analyzed the relationship between RNA abundance (total RNA: counts per million reads [CPM]) and rates of synthesis (alpha: CPM/min) or degradation (gamma: min^-1^) in cortical Ex neuron. As expected, abundant genes (in dark red) exhibited high synthesis and low decay rates, while low-abundance genes (in dark blue) showed the opposite pattern (**Supplementary Fig. 13a**), supporting a model in which total RNA levels are regulated through coordinated control of RNA synthesis and decay *in vivo*.

RNA kinetics regulation is closely linked to gene functions. For instance, mRNAs encoding Ex-neuron specific transcriptional regulators had shorter half-lives (yellow in **Supplementary Fig. 13b**; n=115, mean half-life: 4.7-hr), whereas neurite-enriched mRNAs ^75^, typically involved in local translation and structural functions, were more stable (black in **Supplementary Fig. 13b**; n=101, mean half-life: 11.9-hr). Higher RNA stability likely facilitates long-range transport of these mRNAs from the nucleus to distal compartments (e.g. axons and dendrites) in highly polarized neurons ^11^. Curiously, while many cortical Ex neuron specific transcriptional regulators were encoded by unstable transcripts (n=66, mean half-life: 2.7-hr; e.g. *Rorb*: 1.8-hr, *Satb1*: 2.1-hr, *Pou3f1*: 2.4-hr), a subset of these mRNAs was unexpectedly stable (n=49, mean half-life: 7.5-hr). Many of these stable mRNAs encode Zinc-finger domain containing DNA binding proteins (adjusted *P*=4.4×10^-6^), typically involved in transcriptional repression (e.g. *Scrt1*: 10.8-hr, *L3mbtl3*: 6.7-hr, and *Ikzf2*: 4.2-hr). These results suggest that functionally distinct gene groups employ different RNA kinetics regulatory strategies to precisely regulate transcript abundance.

To further investigate how differential RNA kinetics regulation contributes to RNA abundance *in vivo*, we quantified RNA synthesis, degradation and abundance for 12,483 genes across all 15 cortical cell types. Using *k*-means clustering of scaled synthesis (alpha), degradation (gamma) and expression level (RNA abundance), we identified distinct RNA kinetics modules (**Fig. 4b** and **Supplemental Table S2**). As expected, cell-type specific gene expression patterns were often driven by regulated RNA synthesis (**Fig. 4b**). For example, the abundance of neurite-localized mRNAs, enriched in kinetics modules #11 and #34 (25 out of 51 genes in these two modules, *P*=4.34×10^-40^, Hypergeometric test), was primarily driven by differential RNA synthesis (e.g. *Actb*: 4.8, *Mapt*: 3.0, *Ybx1*: 0.85, *Rps14*: 0.36, and *Tpt1*: 0.25 CPM/min; **Fig. 4c**). Conversely, RNA degradation differentially modulated the RNA abundance of functionally distinct genes within the same cell-type. For example, two neuron-specific modules (#3 and #13) in cortical Ex neurons exhibited comparable synthesis rates (module #3: 0.75 CPM/min, #13: 0.87 CPM/min) but differed substantially in degradation rates (module #3: 9.5×10^-4^ min^-1^, #13: 50.8×10^-4^ min^-1^), resulting in markedly different RNA abundance levels (module #3: 789.3 CPM, #13: 174.1 CPM) (**Fig. 4c** and **Supplementary Fig. 13c**). Thus, module #13 showed substantially shorter RNA half-lives compared to module #3 (mean half-life: 15.2-hr for module #3, 2.34-hr for module #13).

To systematically classify how different RNA kinetics regulatory strategies modulate homeostatic gene expression, we calculated similarity correlations (*r*) between RNA synthesis (alpha; averaged across all genes within each module) and degradation rates (gamma) for 100 RNA kinetics gene modules (**Supplementary Fig. 13d**). Based on the correlations between RNA synthesis and degradation across 15 cortical cell types, these kinetics modules were grouped into three RNA regulatory strategies as previously described ^21,22^: “cooperative” (n=4; negative correlation between RNA synthesis and degradation rates), “neutral” (n=18; weak or no correlation), and “destabilizing” (n=78; positive correlation). These results support earlier observations that “destabilizing” and “neutral” represent the predominant regulatory strategies of RNA abundances across cell-types or states ^21,22^. Furthermore, gene ontology analysis of genes in the top “cooperative” and “destabilizing” modules showed that gene regulated by different regulatory strategies are associated with different cellular functions (**Supplementary Fig. 13e-f**). For example, the fast turnover module #13 followed a “destabilizing” regulatory strategy (*r* = 0.96), with simultaneous up-regulation of RNA synthesis and degradation rates across cortical cell-types. This is consistent with enrichment for fast-turnover, neuron-specific transcription factors in module #13 (e.g. *Pou3f1* and *Rorb*). In contrast, the slow turnover module #3 is associated with a “cooperative” regulatory strategy (*r* =-0.36), exhibiting a moderate negative correlation. Genes in the module #3 preferentially encode synapse-localized proteins (e.g. *Nrgn* and *Stxbp1*), suggesting that cooperative tuning of RNA synthesis and degradation helps maintain high expression levels of these genes in neurons. Overall, these results indicate that distinct regulatory strategies are linked to specific cellular functions, underscoring the contribution of RNA turnover regulation to functional specialization across gene modules.

### Spatial mapping of RNA turnover kinetics across brain regions

To investigate the spatially resolved RNA kinetics regulation in mouse brains, we computed transcriptome-wide RNA synthesis and decay rates across spatial domains in P7 mouse brains. Consistent with *in vivo* single-cell kinetics analyses, spatial profiles of RNA kinetics suggest that RNA abundance is coordinately regulated by synthesis and degradation (**Extended Data Fig. 7a**). Intriguingly, the DG exhibited globally elevated RNA synthesis and degradation, consistent with higher RNA turnover rates in DG compared to non-DG regions (**Extended Data Fig. 7a**). Comparing the DG to other brain regions (CTX-L2/3, CA1, CA3, and Lateral-TH in **Fig. 4d**), we found that genes in kinetics module #3 (slow turnover) but not those in module #13 (fast turnover) exhibited consistent and significant increase in RNA synthesis and degradation rates in the DG (**Fig. 4e** and **Extended Data Fig. 7b**). Similarly, both neurite-localized genes and Ex neuron enriched transcriptional regulators showed significant increase in both RNA synthesis and decay rates in the DG relative to non-DG regions (**Extended Data Fig. 7c-e**). Assuming that total RNA levels are comparable between DG and non-DG regions, these findings reveal a “kinetics scaling” mechanism ^5^ by which turnover of many genes (e.g. module #3 and neurite-localized genes) in the DG are elevated through coordinated regulation of synthesis and degradation. This mechanism may allow neurons in the DG region to remain in a functionally plastic state, enabling rapid induction of ARGs such as *Fos* and *Egr1* in response to external stimuli like ECS.

### *In vivo* analysis of RNA kinetics landscape at single-cell and spatial spot resolution

While clustering enables cell-type or spatial domain-specific analysis of RNA kinetics, single-cell and spatial transcriptomic profiles often exist on a continuum, suggesting that RNA kinetics may exhibit additional heterogeneity within the same cell-type or spatial region. However, the sparsity and noise of single-cell and spatial transcriptomics data pose major challenges for high resolution RNA kinetics analysis.

To address this, we developed *RNAkinetoScope*, a computational framework for measuring transcriptome-wide RNA synthesis and degradation rates at single-cell or spot resolution. By leveraging *k*-nearest neighbor (kNN) graph, *RNAkinetoScope* aggregates newly synthesized (new) and pre-existing (old) transcriptomes from neighboring single cells or spots, improving RNA kinetics inference while preserving intra-cluster heterogeneity. Using cluster-level RNA kinetics as a benchmark (**Supplementary Fig. 14a**), we optimized the local neighborhood size to balance resolution and sensitivity in RNA kinetics analysis (**Supplementary Fig. 14b-c**).

Using *RNAkinetoScope*, we first calculated RNA synthesis and degradation rates for two neuron-specific kinetics modules (#3 vs #13) at single-cell resolution (**Supplementary Fig. 14d**), yielding results highly concordant with those derived from pseudo-aggregated cluster-level data (**Supplementary Fig. 13c**). To uncover intra-cluster heterogeneity of RNA kinetics, we further analyzed gene-specific RNA kinetics at single-cell resolution for representative neuronal genes (e.g. *Stxbp1*, *Nrgn*, *Plppr4*, and *Rorb* in **Supplementary Fig. 14e-f**). Notably, *Stxbp1* and *Nrgn*, both involved in synaptic signaling, exhibited variable RNA synthesis rates across different excitatory neuronal subtypes, yet maintained high expression levels, likely via enhanced RNA stability in neurons. Conversely, *Rorb*, a TF specific to Ex-L4 neurons, showed both elevated synthesis and degradation to achieve its cell-type specific gene expression pattern.

We next investigated spatial RNA kinetics landscapes at single spatial spot resolution for neuronal kinetics modules (#3 and #13 in **Fig. 4f**). Consistent with pseudo-bulk analyses (**Fig. 4d**), module #3 exhibited lower degradation and correspondingly higher RNA abundance, while kinetics module #13 showed higher RNA degradation particularly in brain regions associated with higher RNA turnover such as CTX-L2/3, DG, and CA1 (middle panel in **Fig. 4f**). Supporting the “kinetics scaling” effect in the DG, coordinated up-regulation of RNA synthesis and degradation was prominent for module #3 (left in **Fig. 4f**), but less so for fast-turnover module #13 (right in **Fig. 4f**). Spatial kinetics analysis of representative genes at single-spot resolution revealed heterogeneity in gene-specific RNA synthesis and degradation both between and within spatial domains, with RNA abundance shaped not only by transcription but also brain region-specific degradation (**Fig. 4g** and **Extended Data Fig. 8a**). For instance, *Snca* and *Map4* (module #3 genes in **Fig. 4g**) were broadly synthesized at high levels across hippocampal and cortical regions but showed reduced RNA abundance specifically in DG, relative to CA1/CA3 or cortex, due to elevated degradation in the DG. Similarly, neurite-enriched genes exhibited higher RNA synthesis and decay rates specifically in DG, ensuring that genes with core neuronal functions are expressed at proper levels in this high-turnover brain region (**Extended Data Fig. 8b**).

Collectively, we have developed “*in vivo Timscope*”, an integrated experimental (*in vivo* RNA labeling coupled with single-cell or spatial NT-seq) and computational (*RNAkinetoScope*) framework for transcriptome-wide RNA kinetics analysis at cluster level and single cell/spatial-spot resolution. Applying this approach to the mouse brain generated the first cell-type-and spatial-region resolved map of *in vivo* RNA kinetics landscapes in mammalian brains, highlighting the intricate relationship between gene function and RNA turnover dynamics.

### Predictive modeling of RNA stability regulatory landscape in mouse brains

While transcription factor-DNA interactions are well-characterized, the RNA-encoded *cis*-elements and *trans*-acting factors that regulate RNA stability within cellular and spatial context *in vivo* remains poorly defined. To address this gap, we sought to leverage *in vivo* measurements of transcriptome-wide mRNA half-life to map the genetic features and molecular regulators influencing RNA stability across diverse *in vivo* cell-types and brain regions using recently developed machine learning approaches ^76,77^ (**Fig. 5a**).

**Fig. 5.**
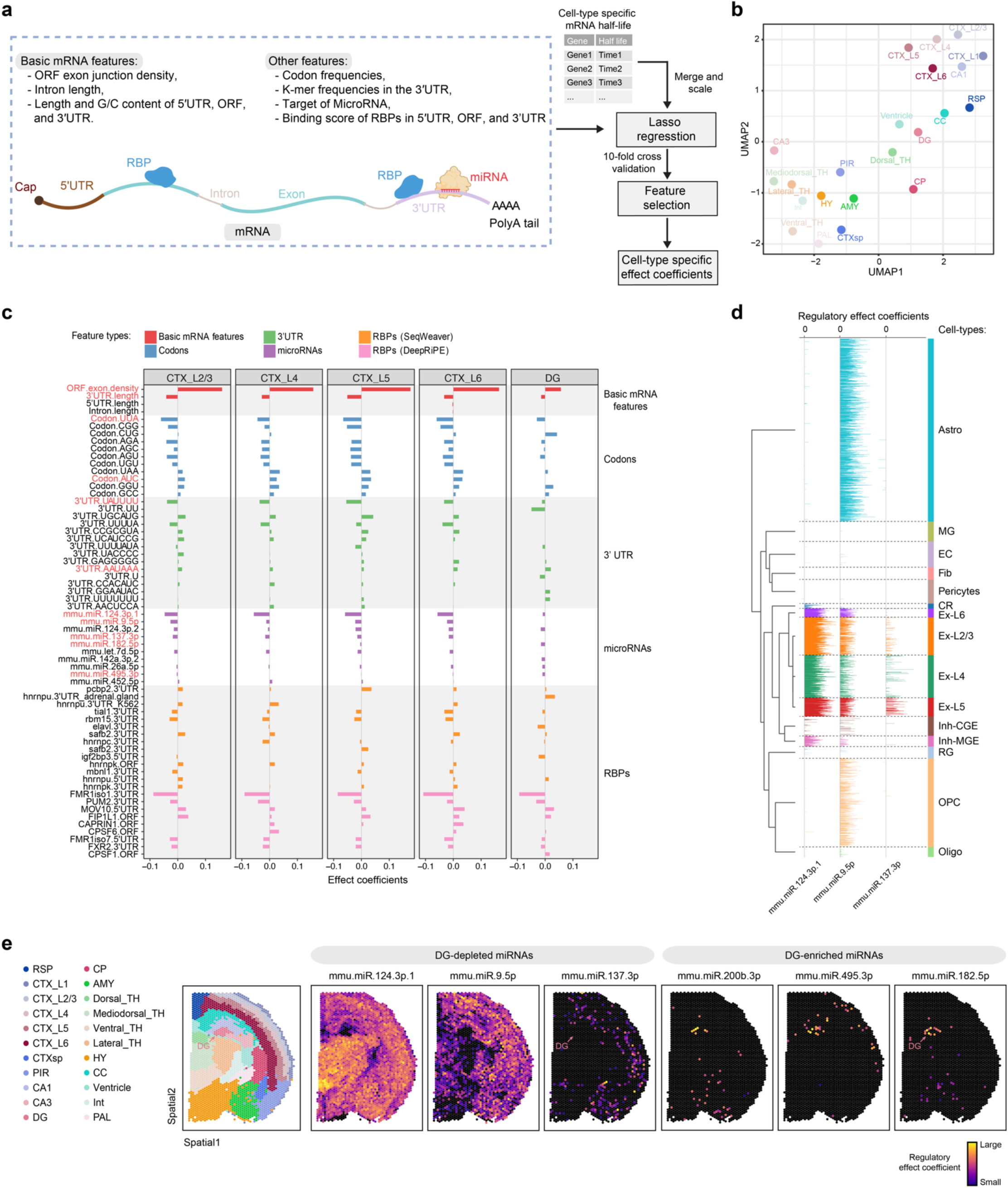
Machine learning based analysis of RNA half-life profiles to reveal *in vivo* mRNA stability regulation across cellular and spatial contexts in the mouse brain. **a.** Schematic diagram of the machine learning based computational pipeline to analyze post-transcriptional regulators of mRNA stability with the key features used in the model highlighted. ORF, open reading frame; RBP, RNA-binding protein; UTR, untranslated region. **b.** UMAP visualization of annotated spatial regions based on the mRNA half-lives of top 3000 highly variable genes. Brain region abbreviations: RSP, retrosplenial area; CTX, cortex; CTXsp, cortical subplate; PIR, piriform area; CA, cornu ammonis; DG, dentate gyrus; CP, caudoputamen; AMY, amygdalar nucleus; TH, thalamus; HY, hypothalamus; CC, corpus callosum; Int, internal capsule; PAL, pallidum. **c.** The top 5 features in each category associated with post-transcriptional regulation of RNA half-lives in five representative brain regions. The features are colored according to the categories. **d.** Track-plot visualization of the regulatory effect coefficients for three representative miRNAs in cortical cells derived from P7 mouse brains. In the track-plot, the magnitude and directionality of regulatory effect coefficients are represented by the height and direction (relative to “0”: left – positive, right - negative) of each vertical bar (corresponding to a single cell). Note that most of the effect coefficients of the three miRNAs are pointing to the right, indicating the expected negative impact of these miRNAs on mRNA stability. **e.** Spatial mapping of the regulatory effect coefficients for six representative miRNAs, including two pan-brain miRNAs (miRNA-9 and miRNA-124), a cortical-layer specific miRNA (miRNA-137) and three DG-specific miRNAs (miRNA-200b, miRNA-495, and miRNA-182). The spatial annotation of the same mouse brain section (treated-rep1 in Fig. 2b) are provided in the left panel.

First, we demonstrated that our *in vivo* RNA half-life data captured cell-type-and spatial region-specific signatures of RNA stability as these data alone were sufficient to segregate brain cell-types and spatial domains into groups corresponding to annotated cell-types and major anatomical regions (**Fig. 5b** and **Supplementary Fig. 15a**).

Next, we trained and benchmarked a Least Absolute Shrinkage and Selection Operator (LASSO) regression model ^76,77^ to identify molecular features regulating *in vivo* cell-type-specific RNA stability (**Fig. 5a** and **Supplementary Note 2**). The prediction results from trained models, evaluated by held-out data and 10-fold cross-validation (**Extended Data Fig. 9a** and **Supplementary Fig. 15b**), showed strong correlation between predicted and observed RNA half-lives across both cell-types and spatial regions (**Extended Data Fig. 9b** and **Supplementary Fig. 15c**). Systematic evaluations indicate that the trained model performs comparably across labeling strategies, sequencing methods, sample origin, labeling durations, and is reproducible across replicates (**Supplementary Fig. 15d-g**). Importantly, the regulatory effects of top-ranked features are concordant between two replicates across 22 spatial brain regions (**Supplementary Fig. 16a**).

Predictive modeling of RNA half-life in single-cell (**Supplementary Fig. 15h**) or spatial NT-seq datasets (**Supplementary Fig. 16b**) revealed that top-ranked features included both cell-type-agnostic (e.g. basic mRNA sequence properties, codon frequencies) and cell-type-specific features (e.g. miRNAs and RBPs). Consistent with *in vitro* studies ^76,78^, open reading frame (ORF) exon junction density positively correlated with RNA stability, whereas 3’UTR length was negatively associated, across both brain cell-types (**Supplementary Fig. 15i**) and spatial regions (**Fig. 5c**). Codon-specific effects on *in vivo* RNA stability closely aligned with known codon stability coefficients (CSCs) ^79,80^, with codons UUA and AGU linked to reduced stability, and codons AUC and GGU to increased stability (**Fig. 5c** and **Supplementary Fig. 15i**). Interestingly, while cortical layers (CTX_L2/3, L4, L5 and L6) showed similar codon regulatory profiles, DG exhibited distinct patterns (**Fig. 5c**), suggesting brain region-specific modulation of codon optimality mediated mRNA degradation (COMD) ^81^. For instance, the stabilizing effect of codon AUC was reduced, whereas that of CUG was increased in the DG relative to other regions (**Fig. 5c**). The model also uncovered known regulatory motifs enriched in 3’ UTR regions, including AUUUA (the core AU-rich element (ARE): destabilizing mRNA) and AAUAAA (the cleavage and polyadenylation motif: stabilizing mRNA) (**Fig. 5c** and **Supplementary Fig. 15i**). In contrast to basic mRNA features, regulatory effects of miRNA and RBP features exhibited greater cell-type specificity (**Fig. 5c** and **Supplementary Fig. 15i**).

Together, these finding demonstrate that while basic sequence determinants of RNA stability are broadly conserved, the regulatory impact of codon optimality, 3’ UTR motifs, and miRNAs/RBPs tend to vary across brain cell-type and regions. The strong concordance between our model predictions and prior knowledge highlights the accuracy and robustness of our approach for dissecting *in vivo* post-transcriptional regulation of RNA stability.

### *In vivo* mapping of miRNA-and RBP-mediated regulation of RNA stability in the mouse brain

To dissect post-transcriptional regulatory network regulating mRNA stability in the mouse brain, we modeled regulatory effects of both miRNAs and RBPs using *in vivo* single-cell and spatial RNA half-life data. Across cortical neuronal and non-neuronal cell-types, most miRNAs (92%) were predicted to destabilize mRNAs (blue dots in **Supplementary Fig. 17a)**, consistent with their canonical mRNA destabilizing role ^53^, while a minority (8%) appeared to exert a positive effect on mRNA stability (red dots in **Supplementary Fig. 17a**), potentially by interfering with RNA decay pathways ^52,82^. Spatial analysis across brain regions similarly revealed a predominantly destabilizing effect (87% of miRNAs; **Extended Data Fig. 9c**).

Further analysis revealed distinct clusters of miRNAs with cell-type-specific regulatory effects on mRNA stability. For instance, single-cell analysis confirmed that brain-enriched miRNAs ^83^ such as miR-124 and miR-137 showed strong destabilizing effects, predominantly within excitatory neuronal subtypes (“Cortical Ex neuron I” cluster; **Supplementary Fig. 17a**). Our approach also unveiled many miRNA candidates in non-neuronal cell-types such as cortical astrocytes and microglia (**Supplementary Fig. 17a**). Moreover, spatial analysis uncovers brain region-specific effects of miRNAs. For instance, miR-124.3p.1, miR-9.5p, and miR-137.3p showed widespread regulatory effects across cortical and subcortical brain regions (e.g. CTX, TH and HY), but exhibited reduced impact in the DG (**Fig. 5c** and **Extended Data Fig. 9c**). Conversely, some miRNAs (e.g. miR-182.5p, miR-495.3p, and miR-200b.3p) exhibited DG-specific effects (**Extended Data Fig. 9c**).

Leveraging high-resolution RNA half-life data generated by *RNAkinetoScope*, we further inferred miRNA regulatory effects at the single-cell resolution, confirming the mRNA destabilizing impact of key brain-enriched miRNAs and their cell-type specificity (e.g. miR-124.3p.1: Ex and Inh neurons; miR-9.5p: Ex neurons, OPC and Astro; miR-137.3p: specific Ex neuronal subtypes; **Fig. 5d** and **Supplementary Fig. 17b**). Notably, the high-resolution analysis also revealed substantial heterogeneity in miRNA regulatory effects within the same neuronal subtype (e.g. miR-124.3p.1 in Inh-CGE and miR-137.3p in Ex-L5 of **Supplementary Fig. 17b**). Extending the LASSO predictive modeling to high-resolution spatial RNA half-life data uncovered further spatial patterns and heterogeneity in miRNA-mediated regulatory effects, with broad effects for miR-124.3p.1 and spatially restricted effects in DG for miR-182.5p (**Fig. 5e**).

RBPs regulate multiple facets of mRNA metabolism ^84^, including stability ^85,86^, yet their roles in modulating *in vivo* RNA stability remain poorly understood. Utilizing LASSO modeling, we inferred the regulatory effects of diverse RBPs on mRNA stability across diverse brain cell-types and regions (**Supplementary Note 3**). Unlike miRNAs, which mainly destabilize mRNAs, RBPs displayed a balanced mix of stabilizing (45%) and destabilizing (55%) effects on mRNA stability (**Supplementary Fig. 18a**), consistent with the notion that RBPs can elicit positive and negative effects on target mRNA stability ^52,76,84^. Moreover, spatial analysis revealed a similar trend of dual functional roles for RBPs (**Extended Data Fig. 10a**). Predicted effects of well-characterized RBPs (8 out of 9 features) aligned with findings from *in vitro* cultured cells ^76^ (**Supplementary Note 3**). Furthermore, predicted *in vivo* regulatory effects of 5 out of 7 overlapping RBPs (destabilizing: DDX6, LARP4, and RBM15; stabilizing: CPSF6 and IGF2BP3) aligned with results derived from RBP knockdown experiments in HepG2 and K562 cells ^84^. Predictive modeling of RNA half-life data at single-cell/spatial spot resolution further revealed both broadly acting and cell type/region-specific RBP regulatory effects (**Supplementary Fig. 18b** and **Extended Data Fig. 10b**).

Collectively, predictive modeling of *in vivo* single-cell and spatial RNA half-life data provides a high-resolution view of both miRNA and RBP-mediated mRNA stability regulatory landscapes in the mouse brain. By predicting regulatory effects at single-cell or spot levels, beyond cluster-level, our method achieves greater resolution in detecting regulatory effects of genetic and molecular regulators of RNA stability and uncovers additional intra-cluster heterogeneity within individual cell-type or spatial domain.

## Conclusion

Despite growing recognition of its important role in gene regulation and human diseases ^7,87^, the transcriptome-wide landscape of *in vivo* RNA turnover and stability remains largely uncharted ^76^. Using “TU tagging” as a benchmark, we systematically optimized and validated a robust *in vivo* metabolic RNA labeling protocol to quantify cell-type-specific RNA turnover in both embryonic and postnatal mouse brains. By integrating this transgenesis-free labeling approach with a platform-agnostic *in situ* chemical conversion strategy, we developed spatial NT-Seq, the first method enabling spatially resolved co-mapping of newly synthesized and pre-existing RNAs in intact tissues. Although this study focuses on the mammalian brains, the versatility of spatial NT-seq makes it readily adaptable for investigating RNA turnover and stability regulation across diverse *in vivo* model organisms, *ex vivo* human tissue cultures, and *in vitro* human 3D organoid models.

Through simultaneous profiling of RNA abundance, synthesis and degradation in the mouse brain, spatial NT-Seq generated the first *in vivo* spatial map of RNA turnover kinetics, revealing pronounced regional heterogeneity in steady-state RNA turnover. Notably, the DG region exhibited coordinated up-regulation of RNA synthesis and decay without major changes in total RNA levels, consistent with a “kinetics scaling” mechanism in which dynamic control of RNA turnover maintains balanced global gene expression by linking RNA synthesis to decay pathways^5^. Following electroconvulsive stimulation, an effective intervention for severe, treatment-resistant depression, spatial NT-seq further identified the DG as the most ECS-responsive region, showing dynamic, activity-dependent regulation of early and late response genes at both transcriptional and post-transcriptional levels. These findings provide mechanistic insights into the molecular effects of ECS and suggest that enhanced RNA turnover may enable DG cells to rapidly adjust protein output from existing transcript pools ^4^, thereby supporting their cellular plasticity during adult neurogenesis ^88^ and in responses to environmental stimuli such as learning and ECS ^89^. Given the potential interplay between RNA and protein turnover in neurons and glia (**Supplementary Fig. 19 and note** 4), future studies integrating spatially resolved, transcriptome-and proteome-wide half-life measurements will be essential to elucidate how RNA stability influences protein turnover and abundances in the brain, either under homeostatic conditions and in response to diverse physiological, pathological, and therapeutic stimuli.

To dissect the regulatory basis of *in vivo* RNA stability, we applied machine learning to *in vivo* RNA half-life data in the brain, identifying sequence features and post-transcriptional regulators, including miRNAs and RBPs, that modulate RNA stability across spatial and cellular contexts. These findings provide a valuable resource for both *in silico* computational modeling and experimental investigation of RNA stability control, with implications for advancing mRNA therapeutics design ^90^.

In summary, our “*in vivo Timescope*” framework integrates transgenesis-free RNA labeling, spatial multimodal transcriptomics (i.e., co-detection of new, old, and total RNAs), and computational modeling to provide a spatially resolved view of RNA kinetic regulation in the brain. This approach enables new insights into the interplay between transcriptional and post-transcriptional control of gene expression, and provides a broadly applicable platform for studying RNA turnover, cellular state transitions, and dysregulation of RNA stability in development and disease ^7,8,10,91–93^. Future integration with next-generation high-resolution ST platforms ^94–98^ and multiple-pulse labeling using orthogonal nucleoside analogs (4sU and 6-thioguanosine) ^99^ will further illuminate the regulatory interplay between RNA synthesis and decay across complex tissue environments at unprecedented spatiotemporal resolution.

## METHODS

### Mouse models

The *CA>GFPstop>UPRT* (*CAG-LoxP-GFP-3xstop-LoxP-UPRT*; stock no. 021469; RRID:IMSR_JAX:021469) ^23^ and C57BL/6J (stock no. 000664; RRID:IMSR_JAX:000664) mice were obtained from the Jackson Laboratory. The *Nex/Neurod6-Cre* driver line ^32^ (RRID: MGI: 4429523) were generously provided by Klaus-Armin Nave (Max Planck Institute of Experimental Medicine, Göttingen, Germany) and Lazzerini Denchi (The Scripps Research Institute). The *Nex/Neurod6-Cre* mice, originally on a sv129 background, were backcrossed to C57BL/6 for at least 10 generations. All lines were maintained on the C57BL/6J background. Wild-type mice used for experiments were C57BL/6J.

For comparison of 4-thiouracil (4tU) and 4-thiouridine (4sU) labeling, conditional “Ex-UPRT” mice (*CA>GFPstop>UPRT/+*; *Nex-Cre*/+) were generated by crossing heterozygous *CA>GFPstop>UPRT* females with *Nex-Cre*/+ males and genotyping F2 offspring. Genotyping of the *UPRT* transgene was performed as previously described ^23^ using primers: 5’-AGTGACAACCCCTCTGGATG-3’, 5’-CATCGGATCTAGCAGCACA-3’. *Nex-Cre* insertion was genotyped using previously reported primers ^32^: 5’-GAGTCCTGGAATCAGTCTTTTTC-3’, 5’-CCGCATAACCAGTGAAACAG-3’.

All animal procedures were conducted in accordance with the ethical guidelines of the National Institutes of Health (NIH) and were approved by the Institutional Animal Care and Use Committee (IACUC) of the University of Pennsylvania. Mice were group-housed (3-5 per cage) in a controlled environment under a 12-hr light/dark cycle (light on at 7:00 am), 55% relative air humidity, and 22°C room temperature, with ad libitum access to food and water.

### *In vivo* metabolic RNA labeling in embryonic and postnatal animals

To compare TU-tagging (4tU in UPRT transgenic mice) with direct 4sU RNA labeling, littermate *CA>GFPstop>UPRT*/+; *Nex:Cre*/+ mice were injected intraperitoneally with either 4sU (0.8 mg per gram of body weight) or 4tU (0.4 mg/g) solution for 4-hr labeling. Notably, dosages were equimolar, accounting for the molecular weight difference between 4sU (260 g/mol) and 4tU (128 g/mol). Briefly, 4tU (Sigma, 440736) was dissolved in DMSO (250 mg/mL stock) and diluted in corn oil to 50 mg/mL immediately prior to injection, as previously described ^33^. For transgenesis-free *in vivo* metabolic RNA labeling using 4sU (Sigma, T4509), a 100 mg/mL stock solution was prepared in 0.9% sterile saline and stored in aliquots at-20 °C.

To optimize 4sU-based transgenesis-free *in vivo* metabolic RNA labeling, dosage was adjusted by body weight: 0.8 mg/g for early postnatal mice (P7-12) and 1.2 mg/g for pregnant females (E16.5 embryos) and adult mice (ECS experiments). Intraperitoneal injections of 4sU or saline (control) were administered accordingly. For dose-response experiments in P7 pups, 4sU was diluted to 50, 100, or 150 mg/mL to deliver 0.4, 0.8, and 1.2 mg/g at 8 μL/g injection volume. Labeling durations ranged from 2-4 hours. After labeling, mouse cortex or whole brains were collected for scNT-seq2 or spatial NT-seq. For scNT-seq2, tissues were dissociated to single cell suspension; for spatial NT-seq, whole brains were dissected, snap-frozen, embedded in optimal cutting temperature (OCT) compound, and stored at −80 °C following the Visium Tissue Preparation Guide (10xGenomics, CG000240, Rev D).

### Quantification of 4sU in mouse brain by liquid chromatography-tandem mass spectrometry (LC-MS/MS)

Wild-type P7 pups received intraperitoneal injection of 4sU at 0.8 mg/g body weight and were analyzed after 1-, 2-, or 4-hr. Cortical tissues were immediately dissected, weighed, and transferred to Eppendorf tubes for LC-MS/MS analysis in collaboration with Creative Proteomics. Briefly, an internal standard (IS) solution of uridine-d2 in water (2 μL/mg tissue) was added to each sample, followed by homogenization using an MM 400 mixer mill at 30 Hz for 3 minutes. Methanol (18 μL/mg tissue) was then added, and samples were homogenized for additional 2 minutes. After centrifugation at 21,000xg for 10-min, the supernatant was diluted 5-fold with water.

For LC-MS/MS analysis with multiple reaction monitoring (MRM) mode (LC-MRM/MS), 10 μL of each sample and calibration standard solution were injected into a C18 column (2.1×100 mm, 1.8 μm) using an Agilent 1290 UHPLC system coupled to an Agilent 6495C triple-quadrupole (QqQ) mass spectrometer with positive-ion detection. Briefly, gradient elution was performed using 0.1% formic acid in water and methanol as the binary solvents.

Quantification was based on a standard curve prepared from serial dilution of 4sU calibration solutions (ranging from 0.0001-10 nmol/mL) in a concentration-matched IS-containing solvent. The concentrations of 4sU in tissue samples (nmol/g tissue) were calculated using internal standard calibration by interpolating the calibration curve with the respective analyte-to-internal standard peak area ratios measured from the samples.

### Tissue dissection and sample processing for scNT-seq2

For embryonic cortex samples (E16.5), brains was dissected in cold DPBS under a microscope as previously described ^100^, keeping all tissue samples in cold buffers on ice throughout. Cortices from two embryos were pooled and resuspended in 0.5 mL DPBS with 0.01% BSA and 30 μM of transcription inhibitor Actinomycin D (ActD, Sigma, A9415). Tissue was dissociated by pipetting (∼20 times with a P1000 tip, followed by two times using a P200 nested in the P1000 tip) and filtering through a 40 μm cell strainer. Cells were pelleted (300xg, 5 min) and resuspended in 0.5 mL DPBS with 0.01% BSA (without ActD). Cell numbers were determined using the Countess II system, and cells were split into two 1.5 mL LoBind tubes (Eppendorf) for methanol fixation.

For postnatal cortex samples (P7-P12), mice were euthanized by cervical dislocation, and brains were dissected on an ice-cold platform ^101^. Cortical tissues were minced with a razor blade to small pieces and maintained in cold DPBS with 0.01% and 30 μM of ActD. Tissue dissociation was performed using the Worthington Papain Dissociation System with a 30-min enzymatic digestion, following the manufacturer’s protocol with following modifications. Briefly, transcription inhibitor (15 μM ActD) was added to both the digestion (with Papain/DNase) and the density gradient (with albumin-ovomucoid inhibitor) solutions. Cells were pelleted (100xg, 6-min), resuspended in 0.5 mL DPBS with 0.01% BSA (without ActD), counted using the Countess II system, and divided into two 1.5 mL LoBind tubes (Eppendorf) for methanol fixation.

Following tissue dissociation, cells was fixed as previously described ^21,102^. Single-cell suspensions were prepared as outline above and resuspended in 0.4 mL DPBS with 0.01% BSA. The suspension was split into two 1.5 mL LoBind tubes (Eppendorf), and 0.8 mL of methanol was added dropwise to achieve a final concentration of 80% methanol in DPBS. Cells were incubated on ice for 1-hr and then stored (in LoBind tubes) at-80°C for up to a month. For rehydration, cells were centrifuged at 1000xg for 5 min at 4°C, methanol-DPBS solution was carefully removed, and the cell pellet was resuspended in 1 mL DPBS with 0.01% BSA and 0.5% RNase inhibitor (Lucigen, 30281-2). After cell counting with the Countess II system, the cell suspension was diluted to 100 cells/μL and used immediately for the scNT-seq2 experiment.

### Optimization of second strand synthesis in scNT-seq2 using *in vitro* labeled K562 cells

Human K562 cells (ATCC, CCL-243) were cultured in RPMI media supplemented with 10% fetal bovine serum (FBS, Sigma, F6178). For *in vitro* metabolic RNA labeling, the culture media was replaced with new medium containing 100 μM 4sU. After 4-hr labeing, K562 cells were then collected by centrifugation, rinsed once with DPBS, and fixed in methanol as described above.

To enhance the efficiency of second strand synthesis in scNT-seq, we designed and tested a panel of random oligos (**Supplementary Note 1**), guided by previous studies ^21,103^. In the random primer comparison experiment (**Supplementary Fig. 1b-c**), each oligo was added to a 200 μl Klenow-based reaction mixture (Klenow buffer A: 1x Maxima RT buffer (ThermoFisher), 12% PEG-8000, 1 mM dNTPs (Clontech), 5 μM Template Switch Oligo (TSO)-GAATG (TSO-GAATG: /5SpC3/AAGCAGTGGTATCAACGCAGAGTGAATG), 10 μM TSO-random primer, and 1.25 U/μl Klenow exo-(Enzymatics)). Reactions were incubated with bead-bound samples at 37°C for 60 min with end-over-end rotation ^103^.

To further optimize reaction conditions, we compared two different DNA polymerases (**Supplementary Note 1**). First, we tested the Klenow exo-enzyme (Enzymatics) in two different reaction buffers (Klenow Buffer A: see above; 200 μL Buffer B: 1× Blue buffer (Enzymatics), 4% Ficoll PM-400, 1 mM dNTPs (Clontech), 10 μM Template Switch Oligo-random primer, and 1.25 U/μl Klenow exo-(Enzymatics)). For the Klenow-based system, reactions were incubated for 60 min at 37°C with rotation. Second, we tested *Bst* 3.0 DNA polymerase (Bst3, NEB, M0374), which offers strong strand displacement activity and high fidelity (**Supplementary Fig. 1d-f**). The Bst3 reaction mixture included: 1× Isothermal Amplification Buffer II (NEB), 6 mM MgSO4, 4% Ficoll PM-400, 1.4 mM dNTPs (Clontech), 10 μM TSO-random primer, and 0.4 U/μl *Bst* 3.0). The Bst3-based reactions were incubated on ice for 2 min for primer annealing, followed by incubation at 60 °C for 15 min with rotation.

Based on library complexity, uniquely mapping rate, and nucleotide substitution rates, systematic evaluation identified the combination of TSO-N3G2N4B random primer and *Bst* 3.0 DNA polymerase as the most optimal condition for second strand synthesis reaction in scNT-seq2.

### *In vivo* scNT-seq2 library preparation and sequencing

The scNT-seq2 library was generated using the custom-built droplet microfluidics-based scRNA-seq platform ^21^, incorporating an extensively optimized second strand synthesis reaction (see above). The scNT-seq2 consists of the following steps (See **Fig. 1a**):

*Step1. Droplet-based co-encapsulation of cells and barcodes beads.* Single cell suspensions were diluted to 100 cells/μL in DPBS containing 0.01% BSA, and approximately 1.5 mL of this suspension was loaded for each scNT-Seq2 run. Cells were then co-encapsulated with barcoded beads (ChemGenes) using an Aquapel-coated PDMS microfluidic device (μFluidix) connected to syringe pumps (KD Scientific) via polyethylene tubing (inner diameter: 0.38 mm, Scientific Commodities). Barcoded beads were resuspended at 120 beads/μL in lysis buffer composed of 200 mM Tris-HCl (pH 8.0), 20 mM EDTA, 6% Ficoll PM-400 (GE Healthcare), 0.2% Sarkosyl (Sigma-Aldrich), and 50 mM DTT (freshly prepared on the day of experiment). Flow rates for both cells and beads were set to 3,200 μL/hr, while QX200 droplet generation oil (Bio-rad) was run at 12,500 μL/hr.

*Step2. TimeLapse chemistry-based conversion of 4sU-labelded new RNAs on pooled beads.* Droplets were broken using Perfluoro-1-octanol (Sigma-Aldrich). After droplet breakage, beads were subjected to the on-bead TimeLapse reaction to convert 4sU to cytidine-analog ^38^. Briefly, 50,000-100,000 beads were washed once with 450 μL wash buffer (1 mM EDTA, 100 mM sodium acetate, pH 5.2), then resuspended in a reaction mixture containing TFEA (600 mM), EDTA (1 mM), and sodium acetate (100 mM, pH 5.2) in water. Sodium periodate (NaIO_4_) was added to a final concentration of 10 mM, and the reaction was incubated at 45°C for 1-hr with rotation. Beads were washed once with 1 mL TE buffer, followed by incubation in 0.5 mL of reducing buffer (10 mM DTT, 100 mM NaCl, 10 mM Tris pH 7.4, 1 mM EDTA) at 37°C for 30 min with rotation. Beads were then washed once with 0.3 mL 2x reverse transcription (RT) buffer.

*Step3. Reverse transcription and Bst3-based second-strand synthesis.* For RT reaction, up to 120,000 beads were resuspended in 200 μL of RT mix consisting of 1x Maxima RT buffer (ThermoFisher), 4% Ficoll PM-400, 1 mM dNTPs (Clontech), 1 U/μL RNase inhibitor, 2.5 μM TSO (AAGCAGTGGTATCAACGCAGAGTGAATrGrGrG) ^104^, and 10 U/μL Maxima H Minus

Reverse Transcriptase (ThermoFisher). The RT reaction was incubated at room temperature for 30 min, followed by incubation at 42°C for 120 min. After exonuclease I treatment, pooled beads were washed once with TE-SDS buffer and twice with TE-TW buffer. Beads were then resuspended in 500 μL of 0.1 M NaOH and incubated at room temperature for 5 min with rotation. The solution was neutralized with 500 μL of 0.2 M Tris-HCl (pH 7.5), followed by additional washes once with TE-TW buffer and once with 10 mM Tris-HCl (pH 8.0).

Second-strand cDNA synthesis was carried out using the optimized *Bst*3-based reaction.

Specifically, beads were resuspended in 200 μL of *Bst*3-based reaction mixture containing 1× Isothermal Amplification Buffer II (NEB), 6 mM MgSO4, 4% Ficoll PM-400, 1.4 mM dNTPs (Clontech), 10 μM TSO-N3G2N4B random primer (AAGCAGTGGTATCAACGCAGAGTGA (N1:25252525)(N1)(N1)GG(N1)(N1)(N1) (N1)(N2: 00333433); N1 represents a mixture of A, C, G and T at a 25:25:25:25 ratio, N2 represents a mixture of A, C, G and T at a 0:33:34:33 ratio), and 0.4 U/μL *Bst*3 DNA polymerase (NEB, M0374). Reactions were incubated on ice for 2 min for primer annealing, followed by incubation at 60 °C for 15 min with rotation. Reactions were stopped by washing the beads once with TE-SDS buffer and twice with TE-TW buffer. Downstream steps including cDNA amplification, sample tagmentation, and indexing PCR were performed as previously described ^21^.

*Step4. cDNA amplification.* To determine the optimal number of PCR cycles for cDNA amplification, an aliquot of 6,000 beads was initially amplified by PCR in a 50 μL reaction (25 μL of 2x KAPA HiFi hotstart readymix (KAPA biosystems), 0.4 μL of 100 μM TSO-PCR primer (AAGCAGTGGTATCAACGCAGAGT), 24.6 μL of nuclease-free water) using the following thermal cycling parameter: 95°C for 3 min; 4 cycles of 98°C for 20 sec, 65°C for 45 sec, 72°C for 3 min; 9 cycles of 98°C for 20 sec, 67°C for 45 sec, 72°C for 3 min; a final extension at 72°C for 5 min, hold at 4°C. After two rounds of purification with 0.6x SPRISelect beads (Beckman Coulter), amplified cDNA was eluted with 10 μL water. To assess amplification efficiency, 10% of amplified cDNA was analyzed via real-time quantitative PCR (qPCR) using Applied Biosystems QuantStudio 7 Flex with a 10 μL qPCR reaction (1 μL of purified cDNA, 0.2 μL of 25 μM TSO-PCR primer, 5 μL of 2x KAPA FAST qPCR readymix, and 3.8 μL of water). Based on this, the remaining beads from each sample were divided into multiple cDNA amplification reactions (∼6,000 beads per 50 μL reaction) and amplified for [4 + 9-12] cycles to enrich cDNA.

*Step5. Final library construction and sequencing.* After cDNA amplification, 1 ng of cDNA (pooled in equal amounts from all reactions) was tagmented using the Nextera XT DNA Sample preparation kit (Illumina, FC-131-1096). The tagmented samples was enriched with 12 PCR cycles using the Nextera XT i7 primers and the P5-TSO hybrid primer ^104^. Final libraries were assessed using a Bioanalyzer (Agilent) and sequenced on a NextSeq 500 using the 75-cycle High Output v2.5 Kit (Illumina). Specifically, libraries were loaded at 2.0 pM with 0.3 μM Custom Read1 Primer (GCCTGTCCGCGGAAGCAGTGGTATCAACGCAGAGTAC) added to position 7 of the reagent cartridge. Sequencing was performed in the following configuration: 20 bp (Read1), 8 bp (Index1), and 60 bp (Read2).

### Optimization of *in situ* chemical reaction using human K562 cells

Human K562 cells were metabolically labeled with 4sU and subsequent fixed as described above. We first optimized *in situ* SLAM reaction based on previously reported protocols ^42,105^, benchmarking its performance against the established on-bead TimeLapse chemistry. Specifically, chemical conversion was tested directly in the fixation solution (80% methanol and 20% DPBS) using varying concentrations of iodoacetamide (IAA, ranging from 10-40 mM), applied either overnight at room temperature or at 50°C. For the 50°C condition, the reaction was pre-incubated on ice for 15 min before incubation at 50°C with rotation. Following the reaction, cells were rinsed with 1 mL cold fixation solution (80% methanol and 20% DPBS) and incubated for 5 min at room temperature in 1 mL quenching buffer (0.1% BSA, 1 U/uL RNase-Inhibitor, 100 mM DTT in DPBS). Library preparation proceeded using the scNT-seq2 protocol (omitting the on-bead conversion step). For samples treated at 50°C, the Bst3-based second strand synthesis reaction awas incorporated following reverse transcription and exonuclease I treatment.

### *In situ* chemical reaction and spatial NT-seq library preparation

To develop spatial NT-seq on a widely adopted spatial transcriptomics platform, we integrated the optimized *in situ* chemical conversion reaction with the Visium platform from 10x Genomics (CG000239 Rev D). Two major modifications were made to the standard Visium workflow: 1) incorporation of the optimized *in situ* SLAM reaction; and 2) replacement of the original “Reverse Transcription Master Mix” and “Second Strand Synthesis” reagents with the “Maxima H Minus Reverse Transcriptase mix” and optimized *Bst*-based second strand synthesis reaction mix, respectively, as used in scNT-seq2. Importantly, these methodological innovations are platform-agnostic and can be adapted to other sequencing-based spatial transcriptomics platforms. The spatial NT-seq workflow consists of the following steps (see **Fig. 2a**):

*Step1. Tissue sample preparation.* Following *in vivo* metabolic RNA labeling (see above), mice were euthanized and brains were dissected, embedded in optimal cutting temperature (OCT) compound (Tissue-Tek), and stored at −80 °C.

*Step2. Cryosection followed by H&E staining.* OCT-embedded frozen brain hemispheres were cryo-sectioned (10 μm thickness) at −20 °C using a cryostat (Leica, CM3050S). Methanol fixation, H&E staining, and tissue imaging were performed according to the Visium User Guide (10x Genomics, CG000160 Rev C) using the EVOS FL Auto 2 Cell Imaging System to generate tissue image files for downstream analysis.

*Step3. In situ chemical conversion.* The Visium Gene Expression Slide containing tissue sections were placed into the slide cassette and washed once with 100 μL methanol-based fixation buffer per well. The fixation buffer was then replaced with 80 μL of SLAM reaction mix, followed by incubation on ice for 15 min and then at 50 °C for 15 min in a thermal cycler. The reaction was quenched by adding 100 μL quenching buffer (DPBS with 0.1% BSA, 1 U/uL RNase-Inhibitor, and 100 mM DTT), followed by incubation at room temperature for 5 min.

*Step4. Permeabilization and Reverse Transcription.* Permeabilization and reverse transcription steps were performed immediately following the *in situ* SLAM reaction. Permeabilization time was first optimized with the Visium Tissue Optimization kit (12-18 min, 37°C). To enhance the T-to-C conversion efficiency, the original “Reverse Transcription Master Mix” was replaced with the scNT-seq2 RT mix: 1x Maxima RT buffer (ThermoFisher), 4% Ficoll PM-400, 1 mM dNTPs (Clontech), 1 U/μL RNase inhibitor, 2.5 μM Template Switch Oligo (TSO:

AAGCAGTGGTATCAACGCAGAGTGAATrGrGrG), and 10 U/ μL Maxima H Minus Reverse Transcriptase (ThermoFisher). The RT reaction was incubated at room temperature for 30 min, followed by incubation at 42°C for 120 min.

*Step5. Second strand synthesis.* The standard second strand synthesis reaction mix was replaced with a *Bst*-based reaction mixture: 1× Isothermal Amplification Buffer II (NEB), 6 mM MgSO_4_, 4% Ficoll PM-400, 1.4 mM dNTPs (Clontech), 10 μM random primer (TSO-N3G2N4B: AAGCAGTGGTATCAACGCAGAGTGA(N1:25252525)(N1)(N1)GG(N1)(N1)(N1)(N1)(N2: 00333433); N1 represents a mixture of A, T, C and G at a 25:25:25:25 ratio, N2 represents a mixture of A, T, C and G at a 0:33:34:33 ratio), and 0.4 U/μl *Bst* DNA polymerase (NEB, M0374). After adding reaction mix, the slide was incubated on ice for 2 min, followed by incubation at 60 °C for 30 min in a thermal cycler. The second strand cDNA molecules were eluted before cDNA amplification.

*Step6. cDNA amplification.* To determine the optimal number of PCR cycles for amplification, 1 μL of cDNA was used to perform real-time qPCR analysis (1 μL of cDNA, 0.2 μL of 25 μM TSO-PCR primer (AAGCAGTGGTATCAACGCAGAGT), 0.3 μL of cDNA primers (10x Genomics), 5 μL of 2x KAPA FAST qPCR readymix, and 3.5 μL of water) using the QuantStudio 7 Flex (Applied Biosystems). The remaining cDNA was amplified in a 100 μL reaction containing 35 μL of cDNA, 50 μL of 2x Amp Mix (10x Genomics), 3 μL of 25 μM TSO-PCR primer, 12 μL of cDNA primers (10X Genomics). Thermal cycling conditions were: 98°C for 3 min; 12-13 cycles of 98°C for 15 sec, 63°C for 20 sec, 72°C for 1 min; 72°C for 1 min, hold at 4°C.

*Step7. Library construction and sequencing.* Final library construction was performed following the standard Visium workflow, including fragmentation, end repair and A-tailing, adaptor ligation, and index PCR. Libraries were quality-checked using an Agilent Bioanalyzer and sequenced on an Illumina NextSeq 500 instrument using the 150-cycle High Output v2 or v2.5 Kit (Illumina). Libraries were loaded at 2.0 pM and sequencing was performed with the following configuration: 28 bp (Read1), 10 bp (Index1), 10 bp (Index2), and 110 bp (Read2).

### Electroconvulsive stimulation (ECS)

Prior to stimulation, both ECS-treated and control mice underwent a 3-hr *in vivo* metabolic RNA labeling via intraperitoneal injections of 4sU (1.2 mg/g body weight). For ECS treatment, 8-week-old C57BL/6J male mice were subjected to a 1.0 s electrical stimulus (total duration) consisting of 100-Hz, 18-mA, 0.3-ms pulses, delivered using an Ugo Basile ECT unit (model 57800) as previously described ^60^. Control mice were handled identically in parallel bud did not receive the electrical stimulation. Mice were euthanized either 30 min or 120 min following ECS. For each of three time points, including non-ECS (control), 30 min post-ECS (ECS-30min), and 120 min post-ECS (ECS-120min), biologically independent samples were collected from two adult mice.

Whole brains were dissected and embedded in OCT compound as described for the spatial NT-seq. OCT-embedded tissue blocks were stored at-80°C in a sealed container. For the non-ECS mice, four coronal brain sections were collected from each mouse: two for the *in situ* SLAM reaction and the other two as non-reaction controls. For ECS-treated mice, two coronal sections per mouse were harvested for spatial NT-seq experiments.

### scNT-seq2 data preprocessing

Pre-processing of scNT-seq2 datasets, including read alignment, quality filtering, and count matrix generation, was performed utilizing the ‘*Dynast*’ pipeline (version 1.0.1; https://github.com/aristoteleo/dynast-release), an inclusive and efficient toolkit for processing metabolic labeling-based scRNA-seq datasets. Modifications were made to accommodate the droplet-based scNT-seq2 workflow. The pipeline four main steps: ‘*dynast align*’, ‘*dynast consensus*’, ‘*dynast count*’, and ‘*dynast estimate*’.

First, raw sequencing reads were aligned to the mouse reference genome (GRCm39) using’dynast align’ with paired-end mode (i.e., “--strand forward-x dropseq”). High-quality cells were selected with the parameter “--soloCellFilter TopCells” from’*STARsolo*’ to generate a cell-barcode whitelist for downstream analysis. Second, ‘dynast consensus’ constructed a consensus sequence for each transcript by pooling reads with the same UMI index and selecting the most frequent variant at each position as previously described ^21^.’*dynast count*’ was then used to quantify unspliced, spliced, newly-synthesized, and total RNAs for each cell. Moreover, sites with background T-to-C substitutions were identified from control samples without chemical conversion and excluded for further analysis. Finally, ‘*dynast estimate*’ applied a UMI-based binomial mixture model to statistically infer the new RNA fractions, correcting for incomplete new RNA labeling. To improve computational efficiency, we used the alpha correction model (i.e., “--method alpha”), as previously described ^21^. The corrected count matrices of new and old RNAs were generated by ‘dynast estimate’.

### Spatial NT-seq data preprocessing

Pre-processing of Visium platform-based Spatial NT-seq data was first conducted using *’Space Ranger*’ (v2.0.0, 10x Genomics), including read alignment, quality filtering, and generation of total RNA count matrices for each spatially barcoded spot. Reads were aligned to the mouse reference genome (GRCm39), and both exonic and intronic reads were incorporated into count matrices. Histological H&E images were processed using *’Space Ranger*’ to identify regions covered by tissue sections, by aligning pre-recorded spot locations with fiducial border spots in the image.

Subsequent preprocessing steps of spatial NT-seq datasets was performed using the ‘*Dynast*’ pipeline (v1.0.1) as described for the scNT-seq2 dataset. Tailored for the 10x Genomics platform, raw sequencing reads were aligned to the reference genome GRCm39 using’dynast align’ in paired-end mode (i.e., “-x 10xv3”). Spatial spots were filtered using a barcode whitelist generated by *’Space Ranger*’ to retain high-quality spatial spots for downstream analyses. Estimation of wew and old RNAs were obtained as described in the scNT-seq2 preprocessing workflow.

### Cell-type clustering and marker gene identification for scNT-seq2

All downstream data preprocessing, clustering, and differential expression analyses were performed using *Scanpy* (v1.9.1). The total RNA count matrices were imported into *Scanpy* as AnnData objects. The analysis pipeline for scNT-seq2 datasets consists of following main steps:

*Quality control*: For postnatal (P7-12) cortex datasets, cells with fewer than 400 or more than 8000 genes, or with >10% mitochondrial gene content, were excluded. For the E16.5 dataset, cells with fewer than 400 or more than 6000 genes, or >10% mitochondrial gene content were removed. In all scNT-seq2 datasets, genes expressed in <5 cells were removed.

*Doublet detection*: Doublets were identified using the *Scrublet* package. For postnatal (P7-12) cortex datasets, doubles were removed using default parameters. For E16.5 datasets, cells with doublet score higher than 0.35 were excluded.

*Highly variable gene selection*: For postnatal (P7-12) cortex datasets, the top 3000 highly variable genes were selected based on Pearson residuals. For E16.5 datasets, highly variable genes were identified using the ‘sc.pp.highly_variable_genes’ function with parameters (min_mean=0.0125, max_mean=3, min_disp=0.5).

*Batch correction*: To correct for batch effects across samples, the batch balanced KNN (*bbknn*) algorithm implemented in *Scanpy* was applied.

*Clustering and Visualization*: First, principal component analysis (PCA) was performed by computing the top 50 (P7-12 cortical data) or 40 (E16.5) components. Next, a *k*-nearest neighbor (*kNN*) graph was constructed, and clustering was performed using the Leiden algorithm. Finally, clusters were visualized in two dimeonsions space using Uniform manifold approximation and projection (UMAP).

*Cell-type annotation*: Cell types were annotated based on known marker genes and visualized in the UMAP plot. Marker genes for each cluster were identified using the *Wilcoxon rank-sum* test and visualized using dot plots.

### Identification and annotation of spatial clusters for spatial NT-seq

For spatial NT-seq data analysis, total RNA count matrices generated by ‘*Space Ranger*’ were first imported into *Scanpy* (v1.9.1) as AnnData objects. All downstream data preprocessing, clustering, and differential expression analyses were performed using *Scanpy*. The preprocessing of Spatial NT-seq datasets consists of following main steps.

For P7 mouse brains, spatial spots with less than 200 UMIs, or >15% mitochondrial gene content were filtered out. Genes expressed in fewer than 10 spots were also removed. For P56 (ECS) samples, spatial spots with less than 100 UMIs, or >30% mitochondrial gene content were removed, and genes expressed in fewer than 3 spots were also removed. Highly variable genes were identified using default settings in *Scanpy*, with the top 3000 selected for P7 samples and the top 2000 selected for P56 ECS samples. Batch corrections across multiple samples were performed using the *bbknn* algorithm in Scanpy. Dimensionality reduction was carried out via PCA analysis, retaining the top 50 components for downstream clustering and visualization using UMAP plots.

Spatial clusters identified through this workflow were mapped back to their original anatomical locations using Visium spatial barcodes and annotated based on the Allen common coordinate framework (CCF). The anatomical identity of each spatial cluster was further validated by comparison with a published single-nucleus RNA-seq (snRNA-seq) whole mouse brain reference dataset using the *cell2Loction* algorithm (see below).

### Spatial mapping and validation of cell-types in spatial NT-seq data with *Cell2location*

To validate the annotation of spatial clusters identified in the spatial NT-seq datasets (**Supplementary Fig. 11**: P7; **Supplementary Fig. 14**: P56), we employed ‘*cell2location*’ (v0.1.3; https://github.com/BayraktarLab/cell2location/) to spatially map cell-types in the postnatal mouse brains based on gene expression profiles derived from a published snRNA-seq dataset of adult wild-type whole brains (P56, C57BL/6 mice) ^45^. Cell-type reference signatures from the snRNA-seq dataset were computed using regularized negative binomial regressions, with estimated expression levels calculated as the average value in each cluster. For integrated analysis, raw counts from the spatial NT-seq data were used as input, and genes were filtered to exclude mitochondrial genes and to retain only genes shared between the snRNA-seq reference and spatial NT-seq datasets. The *cell2location* spatial mapping model was trained to infer cell type proportions and spatial gene expression patterns for each section. The model was configured with the default settings except for two hyperparameters: “N_cells_per_location=5” and “detection_alpha=20” as specified in the ‘*cell2location.models.Cell2location*‘ function. This allowed for decomposition of the multicellular RNA count matrix in each spot into reference-derived cell-type signatures, enabling spatially resolved mapping of cell-types in mouse brain sections. For visualization, we used top 5% quantile of the posterior distribution as high confidence cell abundance score.

### Transcriptome-wide analysis of the impact of metabolic RNA labeling on global gene expression

To assess the global effects of 4sU-mediated metabolic labeling on gene expression, we used an previously established approach ^36^. First, we calculated the new-to-total RNA ratio (NTR) using pseudo-aggregated new and total RNA counts from each experiment. When both the labeled and control groups included only a single replicate, the NTR was computed directly as the ratio of new to total RNA for that sample. In cases with multiple replicates in the labeling group, the mean NTR was computed across replicates. To ensure robust downstream analysis, genes with low expression levels (<100 UMIs per sample) were excluded. Differentially expressed genes (DEGs) were identified using the *edgeR* (v3.40.2). Each sample was treated as a pseudo-bulk replicate, and normalization for library complexity was performed. For datasets with replicates, common dispersion was estimated to quantify overall variability across genes, followed by tagwise dispersion to account for gene-specific variability. For datasets lacking replicates, a fixed biological coefficient of variation (bcv) of 0.2 was used to estimate variability. The exact test was performed to identify DEGs using the square of this value (bcv²), and the resulting statistics were merged with corresponding NTR values. Genes were ranked by NTRs (from low to high) and visualized using hexbin plots of NTR rank versus log_2_-scaled fold change (log_2_FC) using ggplot2 (v3.4.1). For additional visualization and interpretation, we also applied empirical thresholds (-3 to +3, depicted by red dash lines) to highlight genes potentially affected by 4sU labeling. These empirical thresholds were informed by the degree of expression variability observed in short *in vitro* 4sU labeling experiments, such as those used in E16.5 cortical primary cells (200μM of 4sU for 2-hr).

### Transcriptome-wide analysis of RNA kinetics at the cluster level

To quantify RNA kinetics parameters at the cluster level (specific cell-type or brain region), an optimized binomial mixture model was first used to statistically correct for incomplete 4sU labeling at the level of individual new transcript (linked by unique molecular identifiers, UMIs) and background T-to-C substitution mutations in unlabeled old RNAs. This approach enables more accurate estimate of new RNA fractions, as previously described ^16,21,38,106^. Specifically, ‘*dynast estimate*’ was used to statistically infer new RNA fractions in “T-to-C” background corrected data. To be computationally efficient, we estimated unlabeled and labeled counts using the alpha correction model, as previously described ^21^.

To generate cluster-level RNA kinetics parameters, corrected new and old transcripts were aggregated by gene across each cell-type or spatial region and normalized to counts per million (CPM) within each cluster. Using these normalized pseudo-bulk values for new and total RNAs, we calculated RNA half-life for each gene at the cluster-level ^107^. The RNA synthesis (alpha) and degradation (gamma) rates were then computed using the ordinary differential equations (ODEs), as previously described ^21,62^. To profile and visualize the transcriptome-wide landscape of RNA abundance and its kinetics parameters (RNA synthesis and degradation rates) across cortical cell-types (scNT-seq2) and spatial brain regions (spatial NT-seq), we established the following analysis pipeline: First, pseudo-bulk RNA half-life was computed for each cell type or spatial domain. Missing values were imputed for genes with more than half of the values across all clusters detected (i.e., not NA) using the median value. Following imputation, RNA synthesis and degradation rates were computed based on the cluster-level RNA half-life. Next, we integrated data from all cortical cell types, including RNA synthesis and degradation rates alongside total RNA abundance, while excluding genes with missing or infinity values. Finally, heatmaps of hierarchical clustering results were generated using the ‘*pheatmap’* (v1.0.12), with column-wise (within cell-type) scaling and the ‘kmeans_k’ parameter to group genes with similar kinetics profiles into modules (**Fig. 3b**).

For unsupervised clustering and dimensionality reduction of commonly detected RNA half-life values from each cell types or spatial domains, we established the following analysis pipeline. First, pseudo-bulk half-life data were filtered to remove genes with missing values. The top 3,000 highly variable genes based on RNA half-life were selected. Second, dimensionality reduction was performed using the top 10 principal components, following by UMAP projection. Finally, cell-types or spatial domains were then clustered based on RNA half-life similarity and visualized in UMAP plots (**Fig. 5b** and **Supplementary Fig. 25a**).

### *RNAKinetoScope*: transcriptome-wide RNA kinetics profiling at the single cell or spatial-spot level

To infer RNA kinetics parameters at the single-cell (for scNT-seq2) or single spatial-spot (for spatial NT-seq) level, we developed *RNAKinetoScope*, a computational pipeline for computing transcriptome-wide RNA synthesis and degradation rates from metabolic RNA labeling datasets. The *RNAKinetoScope* pipeline consists of the following key steps:

*Neighborhood-based smoothing of transcriptome profiles:* Data sparsity and technical noise inherent in single-cell and spatial transcriptomics could negatively affect the accuracy and sensitivity of RNA kinetics analysis at single cell or spatial-spot level. To mitigate this, we implemented a neighborhood-based smoothing approach. For each cell (scNT-seq2) or spatial spot (spatial NT-seq), neighboring cells/spots within the same cluster (cell-types or spatial region) were selected based on transcriptome similarity using Euclidean distance in PCA space (top 30 PCs) via the k-nearest neighbors (*kNN*) algorithm (i.e. ‘*neighbors.NearestNeighbors*’ function from scikit-learn, version 1.1.1) ^108^. Using cluster-level analysis as a ground truth reference, we systematically evaluated the effect of the number of neighbors on the detection sensitivity and chose the optimal neighbor number (n=13 for scNT-seq2; n=10 for spatial NT-seq) for downstream analysis. For each detectable gene, the CP10k-scaled mean of new and total RNAs from each neighborhood was assigned to the corresponding single cell or spatial-spot.

*Estimation of transcriptome-wide RNA half-life at the single cell or spatial-spot level:* Steady-state RNA half-life was calculated for each gene in each single cell/spot:

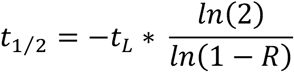

𝑤ℎ𝑒𝑟𝑒 𝑅 𝑖𝑠 𝑡ℎ𝑒 𝑟𝑎𝑡𝑖𝑜 𝑜𝑓 𝑛𝑒𝑤 𝑡𝑜 𝑡𝑜𝑡𝑎𝑙 𝑅𝑁𝐴, 𝑎𝑛𝑑 𝑡*_L_* 𝑖𝑠 𝑡ℎ𝑒 𝑙𝑎𝑏𝑒𝑙𝑖𝑛𝑔 𝑑𝑢𝑟𝑎𝑡𝑖𝑜𝑛.

*Cluster-constrained imputation of gene-specific RNA half-life for individual cell/spot:* To increase the gene coverage of RNA kinetics landscape in individual cell or spatial-spot, gene-specific RNA half-life was further imputed by leveraging information from neighboring cells within the same cell type or spatial region. Specifically, if a gene is detected in >10% of cells/spots within a specific cell or spatial cluster, RNA half-life imputation was performed using a *kNN* approach (i.e. ‘*KNNImputer*’ function from scikit-learn, version 1.1.1) to fill in missing values. Specifically, missing half-life values were imputed by considering the group mean derived from 5 nearest neighbors with available estimates.

*Estimation of RNA synthesis and degradation rates at the single cell/spot level:* Using the RNA half-life and Cell/Spot matrix as the input, we calculate RNA synthesis rate (α, CP10K/min) and RNA degradation rate (γ, 1/min) with following ODE:

The degradation rate constant (γ, units/h) can be calculated from RNA half-life (*t*_1/2_) using

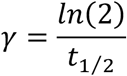

Then, we assumed the gene-specific RNA biogenesis rate (α, molecules/min) is a constant for all cells from each cell state, which can then be calculated using

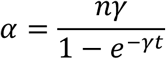

where *n* is the average labeled RNA abundance for each gene in each single cell/spot, γ is the degradation rate constant in each single cell/spot, and *t* (min) is the metabolic labeling time.

### Bootstrap estimation of RNA half-life uncertainty

To quantify uncertainty in half-life estimates, we applied a parametric bootstrap procedure to simulate repeated experiments. For each gene, labeled (L) and unlabeled (U) read counts were observed, with total counts n = L + U and fraction labeled f = L/n. We modeled the number of labeled reads as a binomial random variable, L* ∼ Binomial(n, f), and defined U* = n - L*.

Using this model, we generated 2,000 bootstrap replicates of (L*, U*), each representing a plausible experimental outcome under the same sampling conditions. For each replicate, half-life was recalculated using the same formula as for the observed data. The distribution of bootstrap half-lives was then used to obtain the point estimate (from the observed counts) and a 95% confidence interval (2.5th–97.5th percentile). This simulation-based approach captures the stochastic sampling of labeled and unlabeled reads and provides an approximate measure of uncertainty in half-life estimates.

### Predictive modeling of the post-transcriptional regulome of RNA stability

To model the regulatory effects of sequence-dependent features and post-transcriptional regulators (e.g., miRNAs and RBPs) on RNA stability, we implemented a multi-step machine learning pipeline, as previously described with modifications ^76^. This modified pipeline enabled predictive modeling of RNA half-life at both the cluster (cell-type or spatial domain) level and the single-cell/spatial spot level.

To systematically investigate how various genomic feature sets relate to transcriptome-wide RNA half-life values across cell types, we employed a LASSO regression model using the *glmnet* (v4.1-4) package in *R*. LASSO regression applies an L1 regularization penalty, which is known for selecting a minimal set of features that maximally explain the observed data. All input features considered in the model were z-score normalized and concatenated together. Gene-level feature sets were curated as previously reported ^76^. This analysis established the following feature set as the optimal configuration for collectively explaining *in vivo* RNA half-life: basic mRNA features (“B”, n=8), codon frequencies (“C”, n=61), 3’ UTR *k*-mer motifs (“3”, n=21,844), predicted repression scores of miRNA (“M”, n=315), and predicted RBP binding scores by SeqWeaver (“S”, n=780) or DeepRiPE (“D”, n=177).

For cluster-level modeling analysis, we used RNA half-life values derived from pseudo-bulk samples aggregated across all cells or spatial-spots for each cell type or spatial domain. LASSO regression models were trained independently for each cluster using matched gene-wide feature and RNA half-life data. Genes with infinite or missing half-life values were excluded, and RNA half-life values between 5th and 95th percentile range were retained to mitigate the influence of outliers. RNA half-life values in each cell-type or spatial domain were z-score normalized.

The regularization parameter (denoted as λ) in LASSO regression model controls the strength of the penalty applied to the coefficients. To identify the optimal λ value, 10-fold cross-validation (CV) was performed with different λ value. Model performance was evaluated by calculating Pearson correlation coefficients between predicted and observed RNA half-life values across folds. We also used paired *t*-test to determine significant improvements in performance attributable to specific feature subsets. Following identifying the best feature sets, the LASSO regression model was retrained on the full dataset, and non-zero coefficients were extracted to identify the most informative features. These features were organized into a data frame with corresponding effect sizes, associated cell types/spatial regions, and feature categories. The top-ranked coefficients within each feature class were visualized to highlight major contributions to RNA stability across cellular and spatial contexts.

To extend this approach to single cell/spot resolution, we applied the same modeling pipeline, using neighborhood-level RNA half-life values estimated from the *RNAKinetoScope* as input. Feature sets and preprocessing steps were identical to those used for cluster-level analysis. This enabled predictive modeling of RNA stability at the level of individual cells or spatial-spots and identification of local regulatory influences on RNA stability.

### Differential gene expression analysis for ECS

To identify differentially expressed genes (DEGs) across ECS time points, we performed pairwise comparisons for each spatial cluster using the Wilcoxon rank-sum test implemented in *Scanpy* (v1.9.1), based on both newly synthesized (“new”) and pre-existing (“old”) RNAs. Distinct thresholds were applied: for **Fig. 3d**, DEGs were defined as those with an adjusted *P*-value < 0.001, fold change > 4; for **Supplementary Fig. 15d**, the fold change cutoff was relaxed to >2. To capture temporal gene expression dynamics across the ECS time course, we applied unsupervised clustering analysis of new and old RNAs (CPM) using the *TCseq* R package (v1.22.6). Gene expression patterns of new and old RNAs were grouped into six clusters (*k*=6) by soft clustering (fuzzy c-means). Our analysis focused on gene clusters exhibiting transient upregulation or downregulation at the level of either new or old transcripts.

### Benchmarking RNA labeling-versus splicing-based analysis of spatial IEG dynamics

To quantitatively benchmark RNA labeling-versus splicing-based detection of neuronal activity-induced immediate early gene (IEG) signatures across spatial spots in spatial NT-seq datasets (activated by ECS at 30-min), we adapted “*in silico trapping*”, a computational approach designed to identify cells likely activated by a stimulus from single-cell RNA-seq snapshot data^63^. Per-spot IEG scores were calculated from a curated set of genes (*Arc, Bdnf, Btg2, Fos, Fosl2, Homer1, Npas4*). For each spatial spot, counts from total, unspliced, and new RNA matrices were normalized to counts per 10,000 (CP10k), and the IEG score was defined as the mean normalized expression of the eight-gene panel. Spatial spots with the top 5% of total RNA IEG scores were designated as “trapped” (i.e., IEG^high^ spots) and used as the reference for benchmarking. Predictor scores were similarly defined from unspliced and new RNA IEG data. The ability of each predictor to recover reference-trapped spatial spots was quantified using precision–recall (PR) curves (*precision_recall_curve*) and summarized by average precision (AP; *average_precision_score*), with a random baseline defined by the fraction of reference-trapped spots. To assess overlap and divergence across modalities, the 5% threshold was applied independently to total, unspliced, and new RNA IEG scores. Logical combinations were categorized as “total RNA only”, “unspliced RNA only”, “new RNA only”, “total & unspliced RNA (overlapping)”, and “total & new RNA (overlapping)”. These categories were visualized on UMAP embeddings and summarized in bar plots to evaluate how well new and unspliced RNA recapitulate the ECS-activated spots defined by total RNA.

### Modified iStar pipeline for high-resolution spatial NT-seq analysis

The *iStar* pipeline (https://github.com/daviddaiweizhang/istar) ^66^ was adapted for near single-cell resolution (pixel size: 8 × 8 μm^2^) analysis of Visium-based spatial NT-seq data (with a spatial spot diameter of 55 μm and a center-to-center distance of 100 μm), using the raw histology image and cell-by-gene count matrix as primary inputs. Additional input files included a position information file, a pixel size file, and a radius file, all of which were generated from *CellRanger* outputs. To enable comparisons of gene expression levels across different time points and between new and old transcripts, we omitted the original min-max scaling for normalization step used in the standard iSTAR pipeline. Instead, we utilized a CP10K-normalized gene expression matrix as input. Moreover, we modified the iSTAR visualization code to optimize the color scale for improved clarity and interpretability.

## Data visualization

All plots were generated using *ggplot2* (v3.4.1), *cowplot* (v1.1.1) and pheatmap (v1.0.12) packages in R (v3.5.1). In boxplots, the boxes display the median (center line) and mean (center dot).

## Data availability

All sequencing data associated with this study will be available on the NCBI Gene Expression Omnibus (GEO) database upon publication. The experimental protocols generated in this study will be shared via protocol.io upon publication.

## Code availability

The final version of the analysis source code will be available on GitHub repository upon publication.

## ACKNOWLEDGEMENTS

We are grateful to all members of the Wu lab for helpful discussion. This work is in part supported by a National Human Genome Research Institute (NHGRI) grant U01HG012047, a National Institute of Neurological Disorders and Stroke (NINDS) grant U19NS135528, and a National Eye Institute (NEI) grant U01EY034681.

## AUTHOR CONTRIBUTIONS

Q.Q., H.Z., and H.W. conceived the study. Q.Q. conducted most of the experiments. H.Z. performed most of the computational analysis and developed *RNAkinetoScope*. Q.Q., Z.X., and E.V.H developed and optimized *in vivo* metabolic RNA labeling and spatial NT-seq experimental conditions. Q.Q., D.L., and W.G. developed and optimized scNT-seq2 and *in situ* chemical conversion experimental conditions. Z.X., and Z.Z. generated excitatory neuron specific UPRT transgenic mice. W.G. and Y.L. performed additional data analysis. Q.Q., Z.X., and J.L. performed surgical dissection and single-cell dissociation of postnatal tissues. E.F. and E.K. performed surgical dissection and single-cell dissociation of embryonic mouse brains. Y.S. and H.S. performed ECS experiments. Q.Q., H.Z., Z.X., and H.W. interpreted the data and wrote the manuscript with contributions from all authors.

---

**Extended Data Fig. 1.**
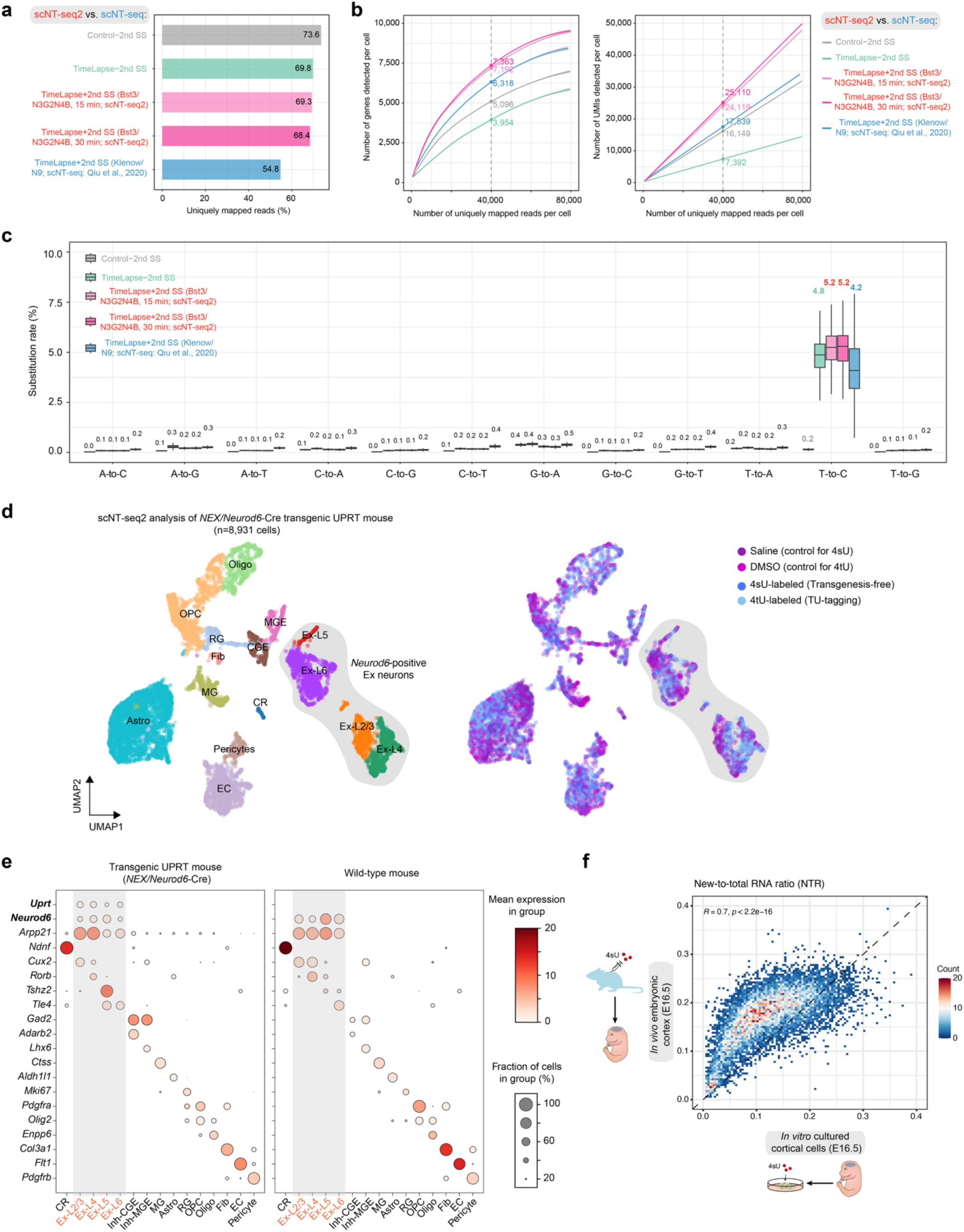
Benchmarking of *in vivo* transgenesis-free metabolic RNA labeling against TU-tagging and *in vitro* RNA labeling using scNT-seq2. **a.** Bar graph showing the fraction of uniquely mapped reads from the experiments either using optimized 2^nd^ SS (scNT-seq2) or the original Klenow-based 2^nd^ SS reaction (scNT-seq). **b.** Fitted line plots illustrating library complexity by comparing genes (left) or UMIs/transcripts (right) detected per cell as a function of aligned reads per cell across experiments in **(a)**. **c.** Box plot comparing nucleotide substitution rates in 4sU-labeled K562 cells from experiments in **(a)**. **d.** UMAP visualization of cortical cells (n=8,931 cells) from four *UPRT* transgenic mice colored by annotated cell types (left) or experimental conditions (right). DMSO injection (marked as DMSO) serves as the control for the 4tU-labeled mouse, while saline injection (marked as saline) is the control for the 4sU-labeled mouse. **e.** Dot plot representing RNA expression levels for 20 marker genes across distinct cortical cell types in conditional *UPRT* transgenic mice (*NEX/Neurod6-Cre; CAG-LoxP-GFP-3xstop-LoxP-UPRT)* at postnatal day 12 (P12) versus wild-type mice. *Uprt*, the transgenic gene coding for uracil phosphoribosyltransferase, which is specifically expressed in *Neurod6*-expressing excitatory neurons. Dot size indicates the percentage of cells expressing the gene in that specific cell type, while color intensity reflects mean gene expression level for that specific cell type. CR, Cajal–Retzius cells; Ex-L2/3/4/5/6, layer specific excitatory neurons; CGE, caudal ganglionic eminence (CGE)-derived interneurons; MGE, medial ganglionic eminence (MGE)-derived interneurons; MG, microglia; Astro, astrocytes; RG, radial glial cells; OPC, oligodendrocyte precursor cells; Oligo, oligodendrocytes; Fib, meningeal fibroblast; EC, endothelial cells. **f.** Scatterplots showing new-to-total RNA ratio (NTR, uncorrected) between *in vivo* labeled E16.5 cortical tissues (y-axis) and *in vitro* labeled cultured E16.5 cortical cells (x-axis).

**Extended Data Fig. 2.**
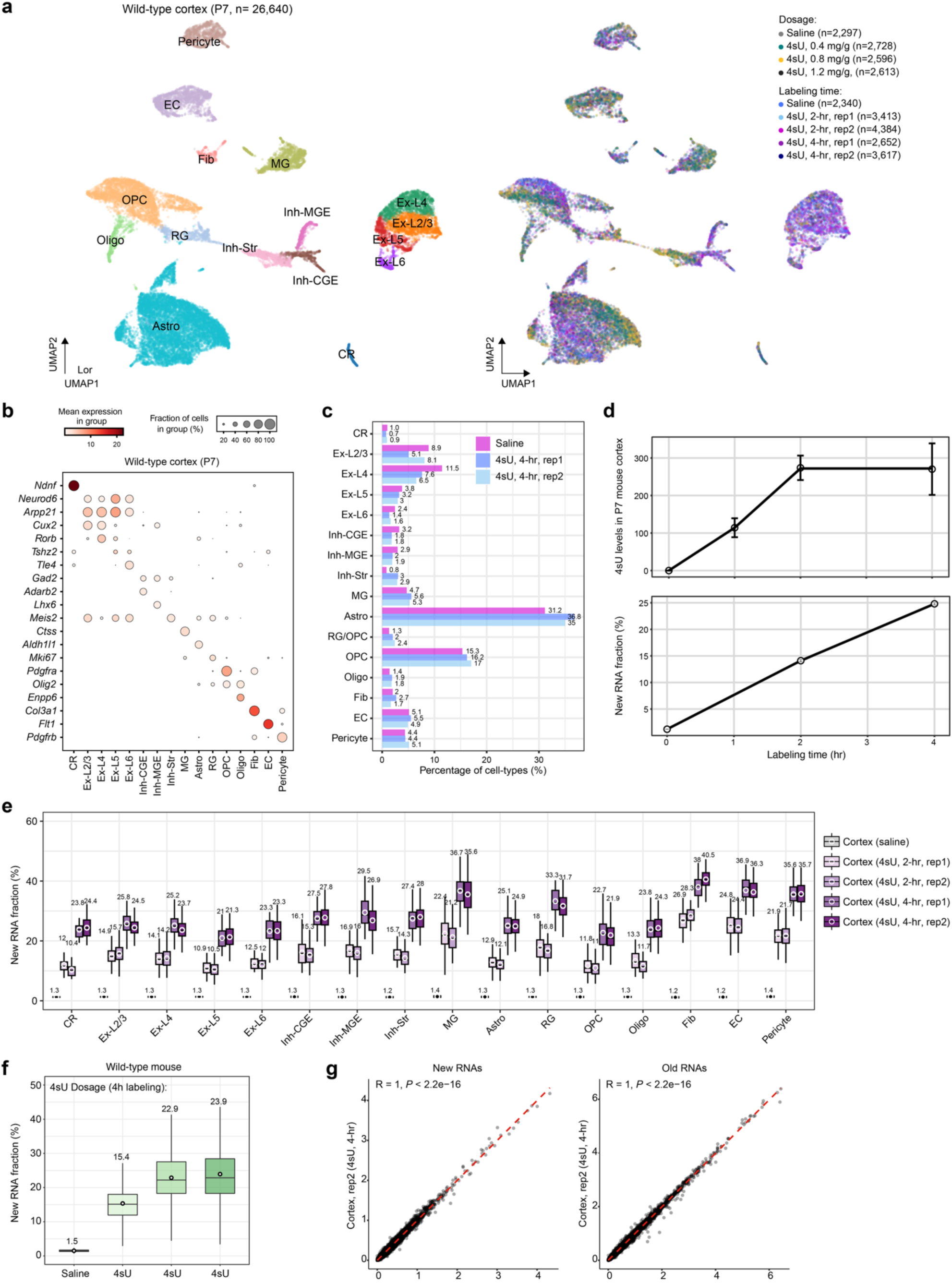
Optimization and validation of *in vivo* transgenesis-free metabolic RNA labeling using scNT-seq2 analysis of wild-type mouse cortical samples. **a.** UMAP visualization of 26,640 high-quality cortical cells from nine P7 wild-type mice, colored by annotated cell types (left) or by experimental conditions (right). Cell numbers for each sample are indicated. Cell type abbreviations: CR, Cajal–Retzius cells; Ex-L2/3/4/5/6, layer specific excitatory neurons; Inh-CGE, caudal ganglionic eminence (CGE)-derived inhibitory neurons; Inh-MGE, medial ganglionic eminence (MGE)-derived inhibitory neurons; Inh-Str, striatum-derived inhibitory neurons; MG, microglia; Astro, astrocytes; RG, radial glial cells; OPC, oligodendrocyte precursor cells; Oligo, oligodendrocytes; Fib, meningeal fibroblast; EC, endothelial cells. **b.** Dot plot illustrating RNA expression levels for 20 marker genes across 16 cortical cell types in P7 cortical tissue. Dot size indicates the percentage of cells expressing a given gene, while color intensity reflects mean gene expression level within that cell type. **c.** Bar plot showing the relative composition of cell types in three samples, including one saline control and two biological replicates of 4sU-labeled (4-hr) mice. **d.** Line plots showing the time course measurments of the 4sU concentration (upper panel: n=2, in units of nmol/mL, see Methods), and 4sU-labeled new RNA fractions (lower panel: n=2) in mouse cortical tissues. The error bars represent standard errors. **e.** Box plot representing the 4sU-labeled new RNA fractions per cell in all major cell types in five samples, including one saline control and two biological replicates of each 4sU-labeling time (2hr or 4hr). **f.** Box plot showing the 4sU-labeled new RNA fractions per cell for samples in 4sU dosage evaluation experiments. Cell numbers for the samples: Saline control (for dosage experiment), 2,297; 4sU-0.4mg/g, 2,728; 4sU-0.8mg/g, 2,596; 4sU-1.2mg/g, 2,613. **g.** Scatterplots indicating the Pearson’s correlation across two biological replicates of P7 cortical samples (4hr-rep1 and 4hr-rep2). Expression data for new and old transcripts (n = 16,020 genes) are scaled to CP10K.

**Extended Data Fig. 3.**
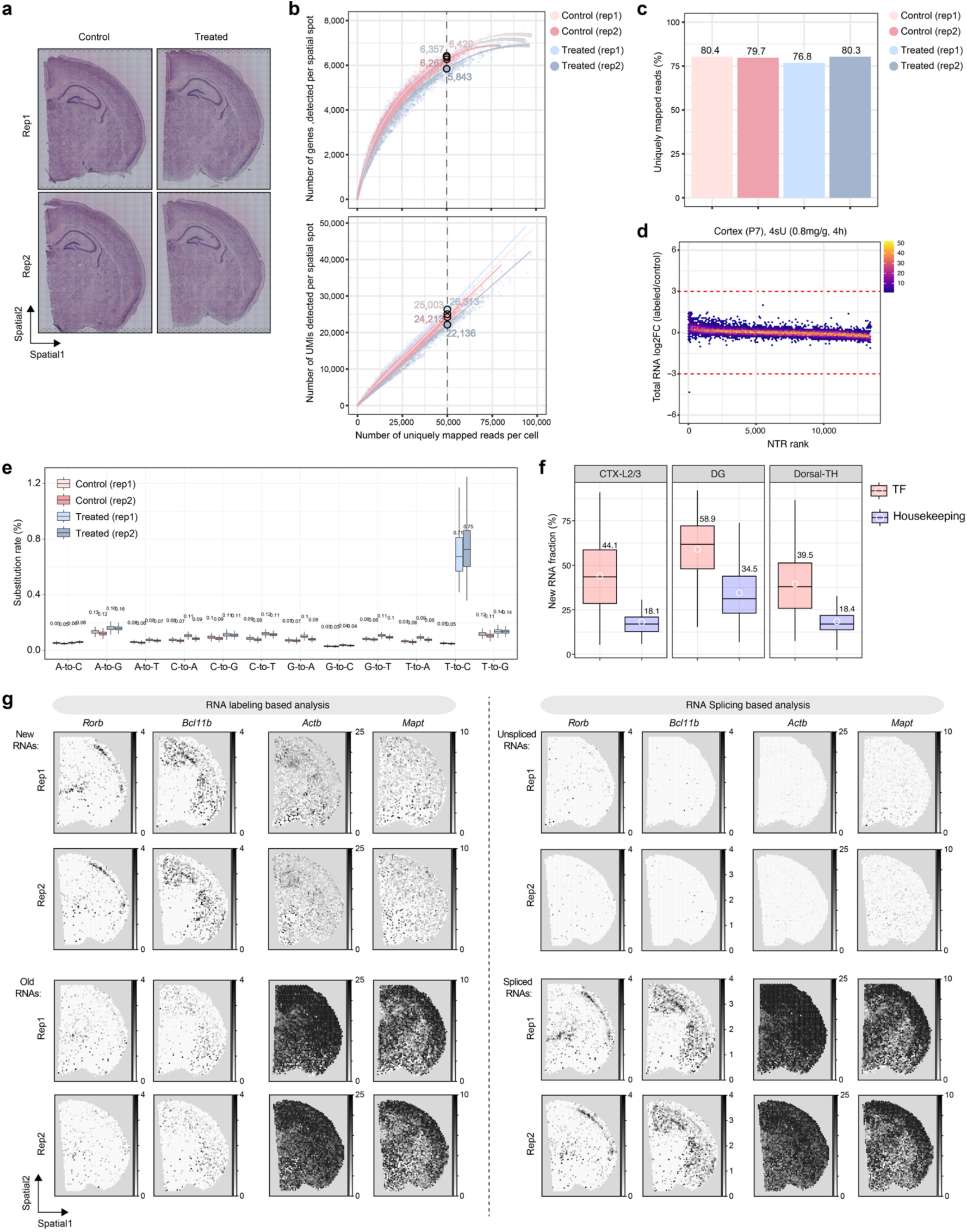
Development and benchmarking of spatial NT-seq using postnatal mouse brains. a. H&E staining images of four coronal mouse brain sections corresponding to those in Fig. 2b. Control, sections without chemical conversion. Treated, sections subjected to *in situ* SLAM chemical conversion. b. Scatter plots illustrating library complexity by comparing genes (left) or UMIs/transcripts (right) detected per spatial spot as a function of aligned reads per spot between control and treated conditions with four tissue sections in Fig. 2b. Estimated numbers of genes or UMIs detected per cell at matching sequencing depth (50,000 reads per spot) for different experiments are shown. The shaded regions depict 95% confidence intervals. c. Bar graph comparing the percentage of uniquely mapped reads between control and treated conditions with four tissue sections. d. Scatter plots comparing the ranks of the new-to-total RNA ratios (NTR) of each gene against the log_2_ fold change of labeled samples versus corresponding control samples in spatial NT-seq experiments. NTR low genes (half-life high) are associated with lower ranks, whereas NTR high genes (half-life low) are associated with higher ranks. The red dash lines depict the limits for identifying potentially affected genes by 4sU labeling. The color density represent gene counts. e. Box plots comparing the nucleotide substitution rates between control and treated conditions with four tissue sections. f. Box plot displaying corrected new RNA fractions derived from spatial NT-seq analysis of P7 brains for TF and housekeeping gene groups in three specific brain regions. Number of genes: TF – CTX-2/3 (n=228), DG (n=300), and Dorsal-TH (n=229); Widely expressed housekeeping genes (n=64) that are derived a previously curated list ^1^. g. Left: normalized spatial expression levels of new (in CP10K; top two rows) and old RNAs (in CP10K; bottom two rows) of four representative genes, including two fast-turnover TF genes (high NTR: *Rorb* and *Bcl11b*) and two slow-turnover housekeeping genes (low NTR: *Actb* and *Mapt*). Right: normalized spatial expression levels of unspliced (in CP10K; top two rows) and spliced RNAs (in CP10K; bottom two rows) of the same four representative genes.

**Extended Data Fig. 4.**
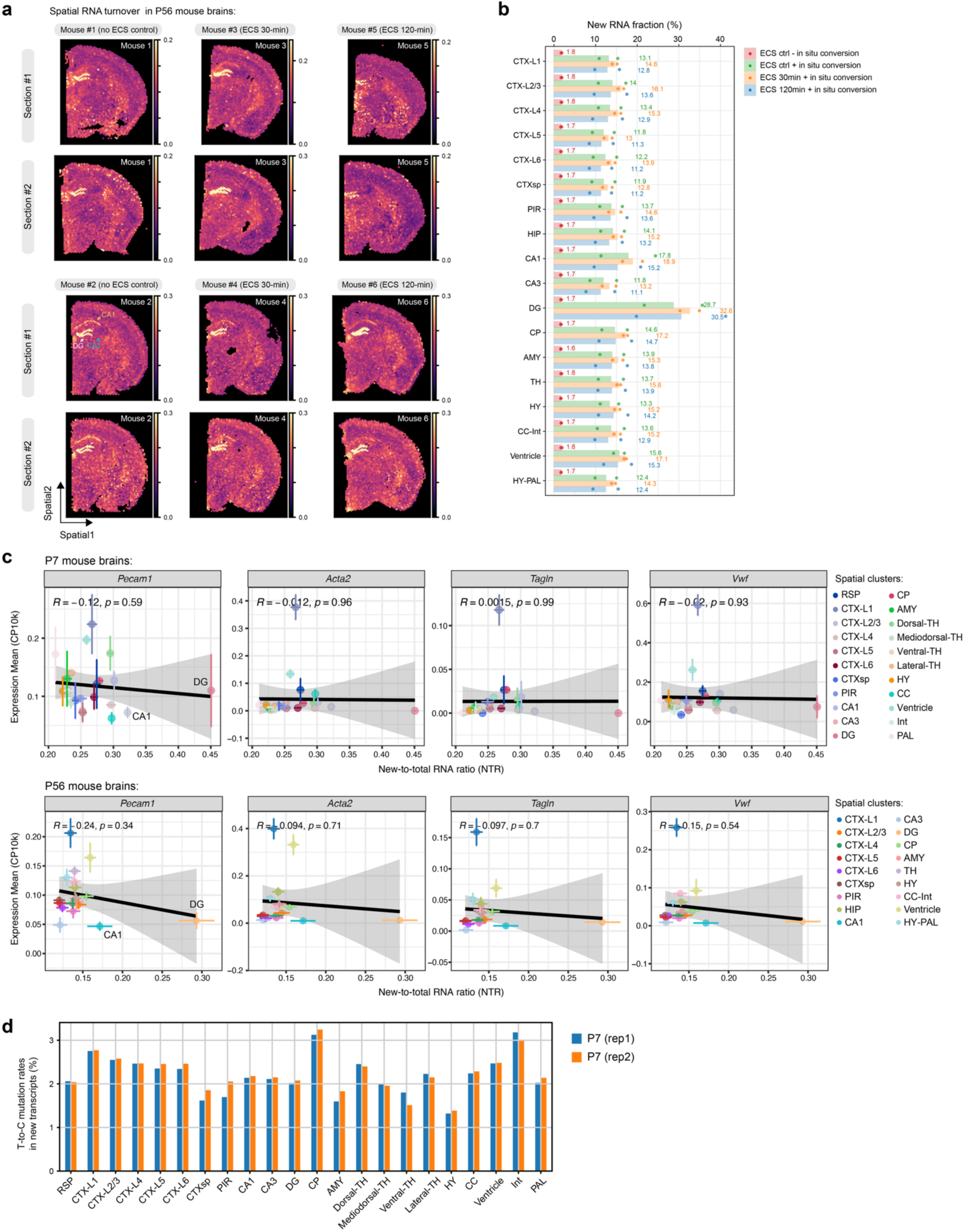
Spatial mapping of global RNA turnover in adult mouse brains. **a.** Spatial mapping of global RNA turnover (NTR) in 12 coronal mouse brain sections, derived from six metabolically labeled wild-type adult mice (P56), across three experimental conditions: No ECS control, ECS-30min, and ECS-120min. ECS, electroconvulsive stimulation. **b.** Bar plot illustrating the fraction of 4sU-labeled newly synthesized transcripts across 18 brain regions averaged over two animals. Each dot represents the mean new RNA fraction of two replicate sections from each animal. **c.** Scatter plots showing the correlation between spatial domain specific RNA turnover (NTRs in x-axis) and normalized total RNA levels (CP10k) of four blood vessel related genes ^2^ (*Pecam1*: capillary; *Acta2*: artery and vein; *Tagln*: artery-specific; *Vwf*: vein-specific) within P7 (upper panels, spatial cluster annotations are from Fig. 2b) and P56 (lower panelsm, spatial cluster annotations are from Fig. 3a) coronal mouse brain sections. Both the vertical (normalized gene expression) and honrizontal (spatial domain specific NTRs) error bars represent standard errors (for P7, n=2 sections; for P56, n=12 sections). The Pearson’s correlation coefficient R and *P*-value are indicated. **d.** Barplot showing T-to-C mutation rates in new transcripts, which is a parameter of the binomial mixture model and primarily reflects 4sU incorporation efficiency ^3^, across all annotated brain regions in two P7 spatial NT-seq replicates. The 4sU incorporation efficiency is relatively comparable across most brain regions (except CP, HY, and Int), with DG and CA1 not exhibiting the highest rate.

**Extended Data Fig. 5.**
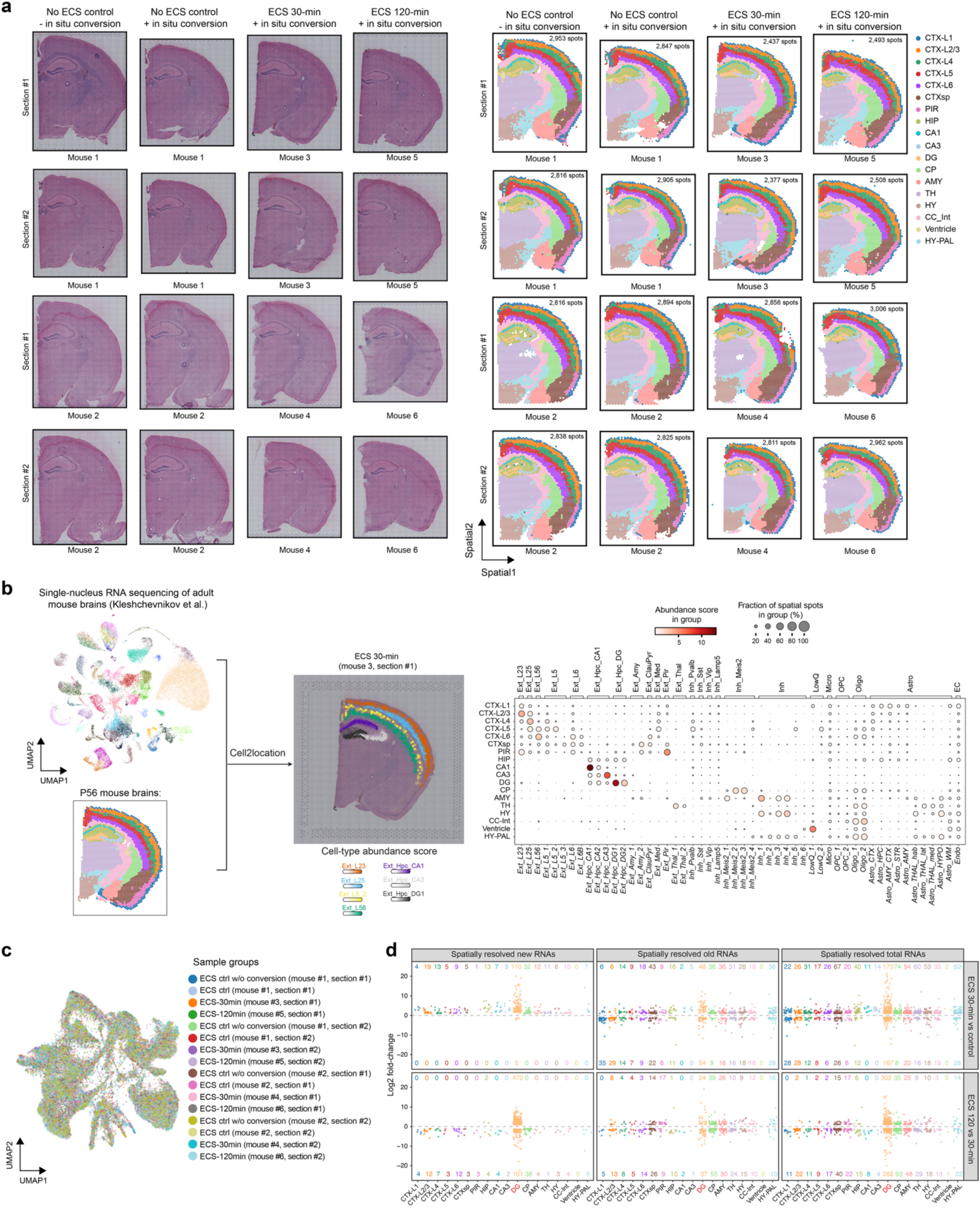
Validation and annotation of spatial NT-seq datasets across ECS time points and biological replicates. **a.** Panels depicting H&E staining images (left) and corresponding spatial annotations (right) of 16 coronal mouse brain sections, derived from six metabolically labeled wild-type adult mice, across three time points: No ECS control, ECS-30min, and ECS-120min. Region abbreviations: CTX, cortex; CTXsp, cortical subplate; PIR, piriform area; HIP, hippocampus; CA, cornu ammonis; DG, dentate gyrus; CP, caudoputamen; AMY, amygdalar nucleus; TH, thalamus; HY, hypothalamus; CC, corpus callosum; Int, internal capsule; PAL, pallidum. Number of spatial spots for each sample are indicated. The low-quality spots are removed from downstream analysis. **b.** Shown on the left is schematics of Cell2location based integration of single-nucleus RNA-seq data ^4^ with spatial NT-seq datasets to generate estimated cell abundance score (indicated by color intensity) for seven excitatory neuronal subtypes within cortical and hippocampal regions on a representative spatial NT-seq section (mouse 3, section #1, ECS 30-min). Dot plots (right) displaying the cell type composition within individual spatial regions, as determined by Cell2location. Dot size represents the fraction of spatial spots in an annotated brain region associated with a specific cell-type, while the color indicates the cell abundance score of that cell type within the regions. **c.** UMAP visualization (as in Fig. 3a) of spatial spots across the 16 brain sections. Colors correspond to sample names. **d.** Scatter plots representing the number and log-scaled FC of differentially regulated new (left), old (middle), and total (right) RNAs (adjusted P-value <0.001 and FC>2) in response to ECS across 18 brain regions for pair-wise comparisons: ECS-30min vs. control (top panels) and ECS-120min vs. ECS-30min (bottom panels). This plot is similar to the one in Fig. 3d but used less stringent cutoff (FC > 2 instead of FC >4).

**Extended Data Fig. 6.**
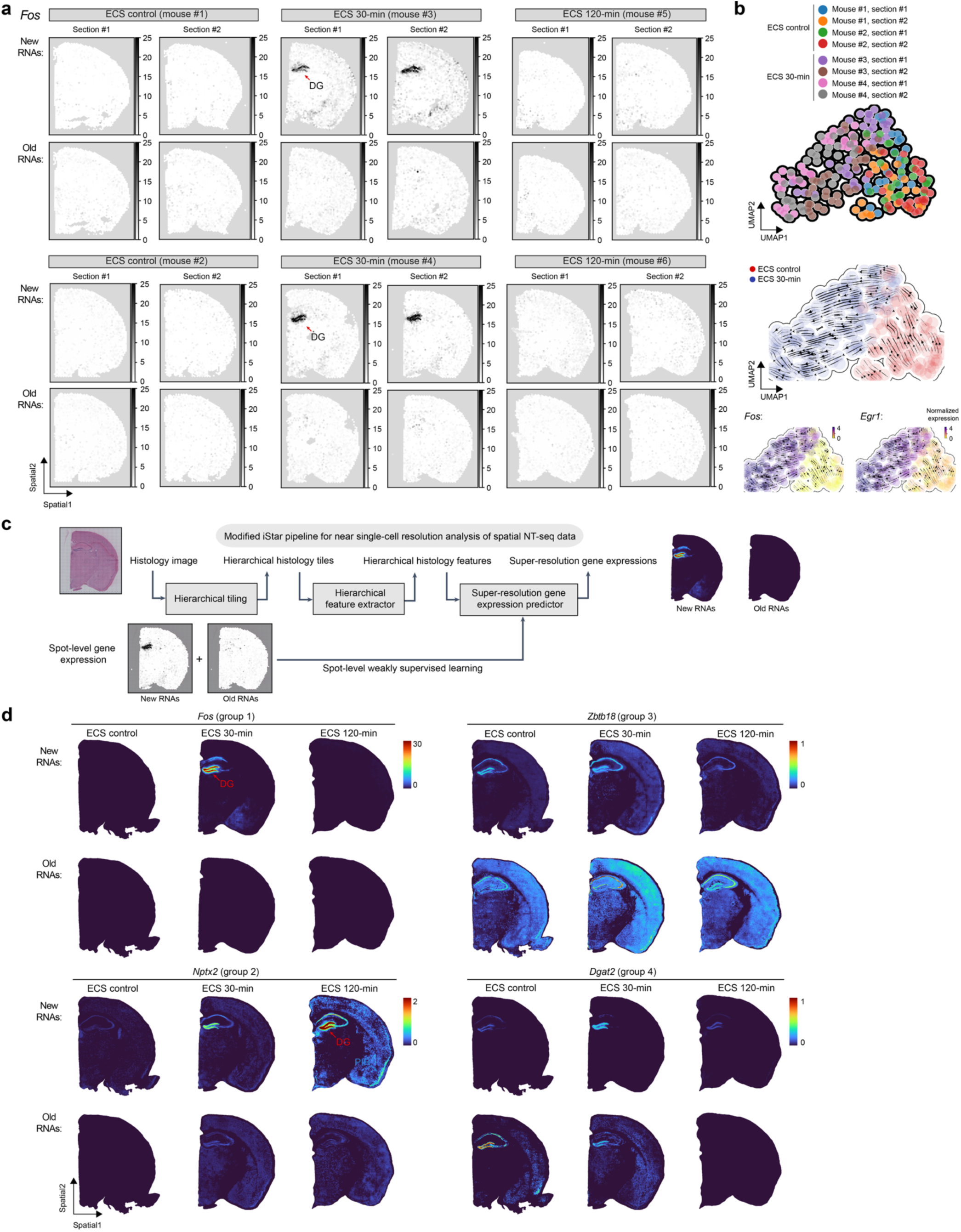
High temporal and spatial resolution profiling of ECS-induced ARG dynamics using spatial NT-seq and computational modeling. **a.** Panels displaying the normalized spatial expression (in CP10K) of new and old RNAs for one representative early ARG gene, *Fos*, across three time points (ECS control, 30-, and 120-min) from two sections (#1-2) of two biologically independent cohort of mice (top: mouse #1/3/5; bottom: mouse #2/4/6). **b.** UMAP visualization (top) of metabolic labeling-based RNA velocity analysis of early ARG activation in response to acute ECS (n = 227 spatial spots in DG). Shown in the bottom are normalized expression (total RNA levels, in CP10K) of two early ARGs, *Fos* and *Egr1*. UMAP visualization of early ARG activation in response to acute ECS (n = 227 spatial spots in DG from two biologically independent cohort of mice) that are colored by experimental samples. **c.** Model summary of modified iStar (inferring Super-resolution Tissue Architecture) workflow ^5^ for near single-cell resolution (pixel size: 8 μm) analysis of newly-synthesized and pre-existing RNA expression derived from spot-level spatial NT-seq data (spot diameter: 55 μm). **d.** Panels displaying the normalized near single-cell resolution (pixel size: 8 μm) spatial expression (in CP10K for input spot-level new and old RNA expression data) of new (upper panel) and old (lower panel) RNAs for four representative genes (same as in Fig. 3g) from each ARG group using a representative spatial NT-seq dataset (section #2 from mouse #2/4/6).

**Extended Data Fig. 7.**
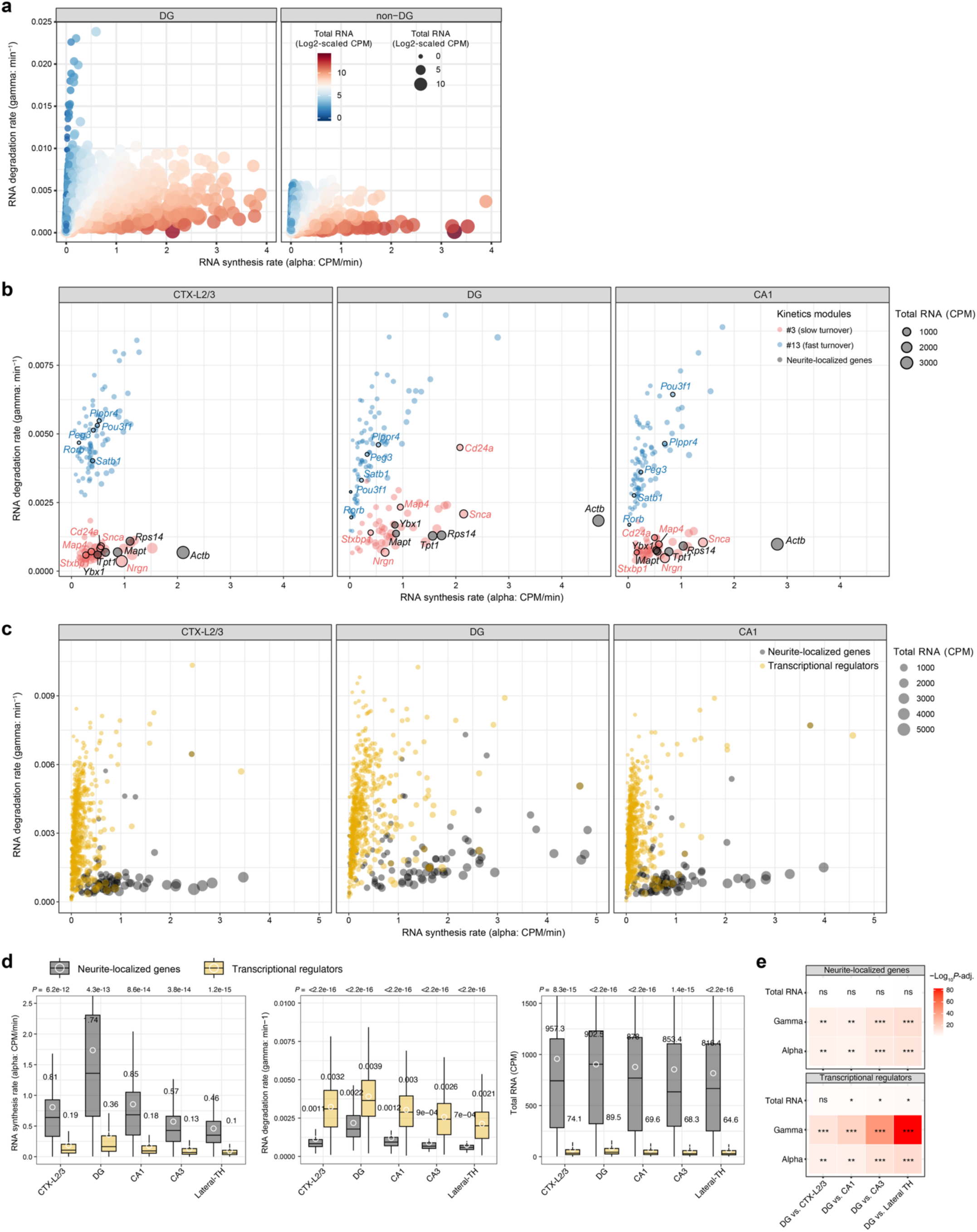
*In vivo* profiling of spatial RNA kinetics landscapes in the mouse brain. **a.** Scatter plot showing transcriptome-wide RNA synthesis (x-axis: CPM/min) versus degradation (y-axis: min^-1^) rates in DG (left) versus non-DG regions (right) of P7 mouse brains. Total RNA levels are represented by dot size and colors (log2-scaled). **b.** Scatter plot showing RNA synthesis (x-axis: CPM/min) versus degradation (y-axis: min-1) rates in three brain regions (P7) for genes in kinetics modules #3 (red: *Stxbp1*, *Nrgn*, *Map4*, *Snca*, and *Cd24a*) and #13 (blue: *Pou3f1*, *Plppr4*, *Peg3*, *Satb1*, and *Rorb*). Five representative neurite-localized/enriched genes (black: *Actb*, *Mapt*, *Rps14*, *Tpt1*, and *Ybx1*) are also highlighted. Total RNA levels are represented by dot size. **c.** Scatter plot showing RNA synthesis (x-axis: CPM/min) versus degradation (y-axis: min^-1^) rates for transcriptional regulators ^6^ (yellow; CPM>20) and neurite-locailized/enriched genes ^7^ (black) in three representative brain regions (P7). Total RNA levels are represented by dot size. **d.** Box plots comparing spatial region-specific RNA synthesis rates (alpha, left), degradation rates (gamma, middle) and total RNA abundance (right) for transcriptional regulators (yellow; CPM>20) and neurite-locailized/enriched genes (black) in five representative brain regions (P7). The mean for each is shown as a white dot, with its value indicated above the box. *P*-values from two-sided *t*-test comparing module #3 and #13 genes are reported above each panel. **e.** Heatmaps showing statistical analysis of spatial region-specific total RNA abundance, RNA degradation rates (gamma), and RNA synthesis rates (alpha) between DG vesus representative non-DG brain regions of P7 mouse brains for neurite-locailized genes and transcriptional regulators. In the heatmap, significance is indicated as follows: ***, *P* < 1×10^-8^; *\*\*, P* < 1×10^-4^; *, *P* < 0.05; ns, non-significant.

**Extended Data Fig. 8.**
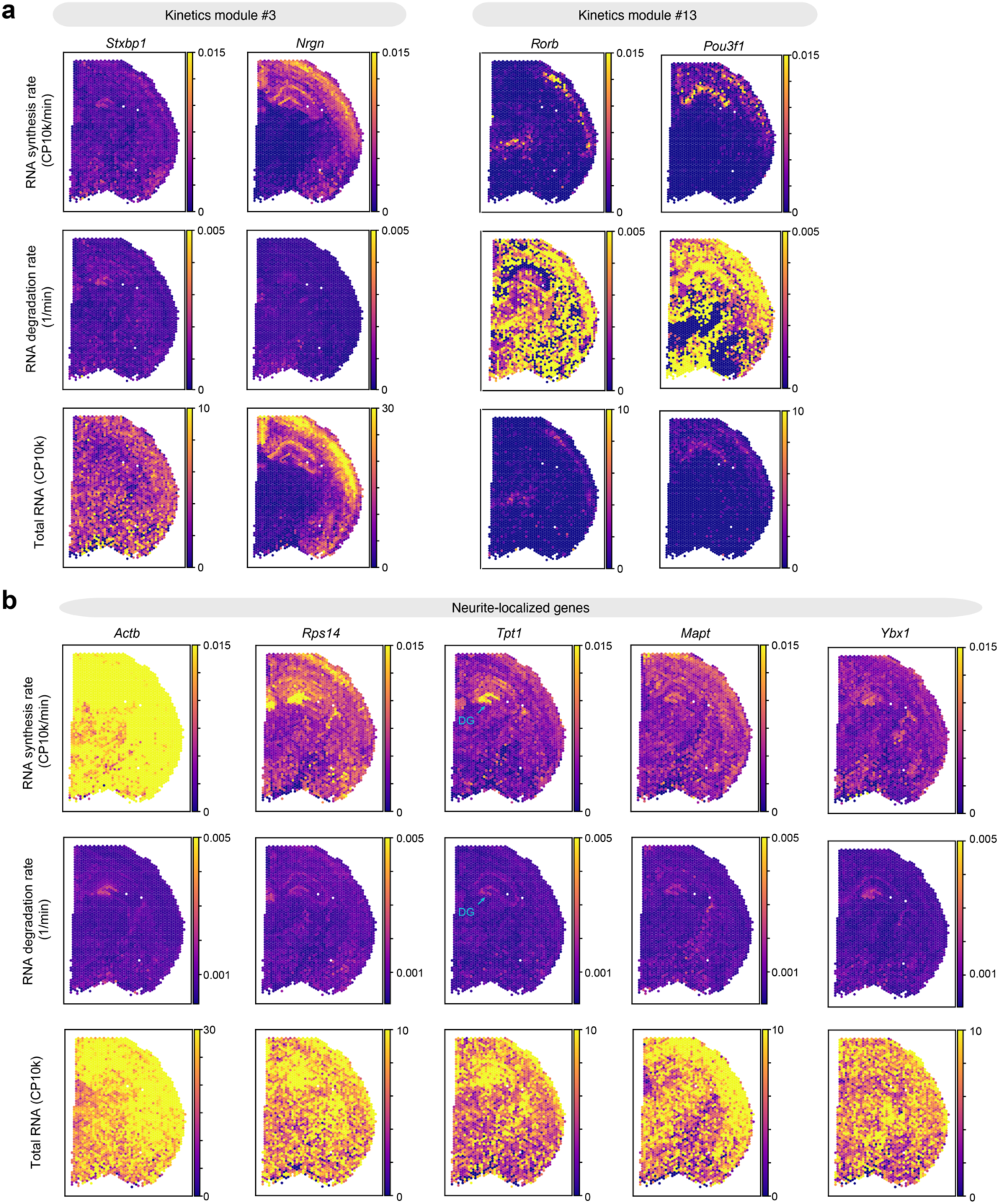
Spatially resolved analysis of RNA kinetics landscapes at single-spot resolution by *RNAKinetoScope*. **a.** Spatial feature plots highlighting RNA synthesis rate/alpha (top), degradation rate/gamma (middle), and total RNA abundance (bottom) for four representative genes: *Stxbp1* and *Nrgn* from module #3 (left panels), along with *Rorb* and *Pou3f1* from module #13 (right panels). **b.** Spatial feature plots highlighting RNA synthesis rate/alpha (top), degradation rate/gamma (middle), and total RNA abundance (bottom) for five neurite-localized/enriched genes (*Actb*, *Rps14*, *Tpt1*, *Mapt*, and *Ybx1*).

**Extended Data Fig. 9.**
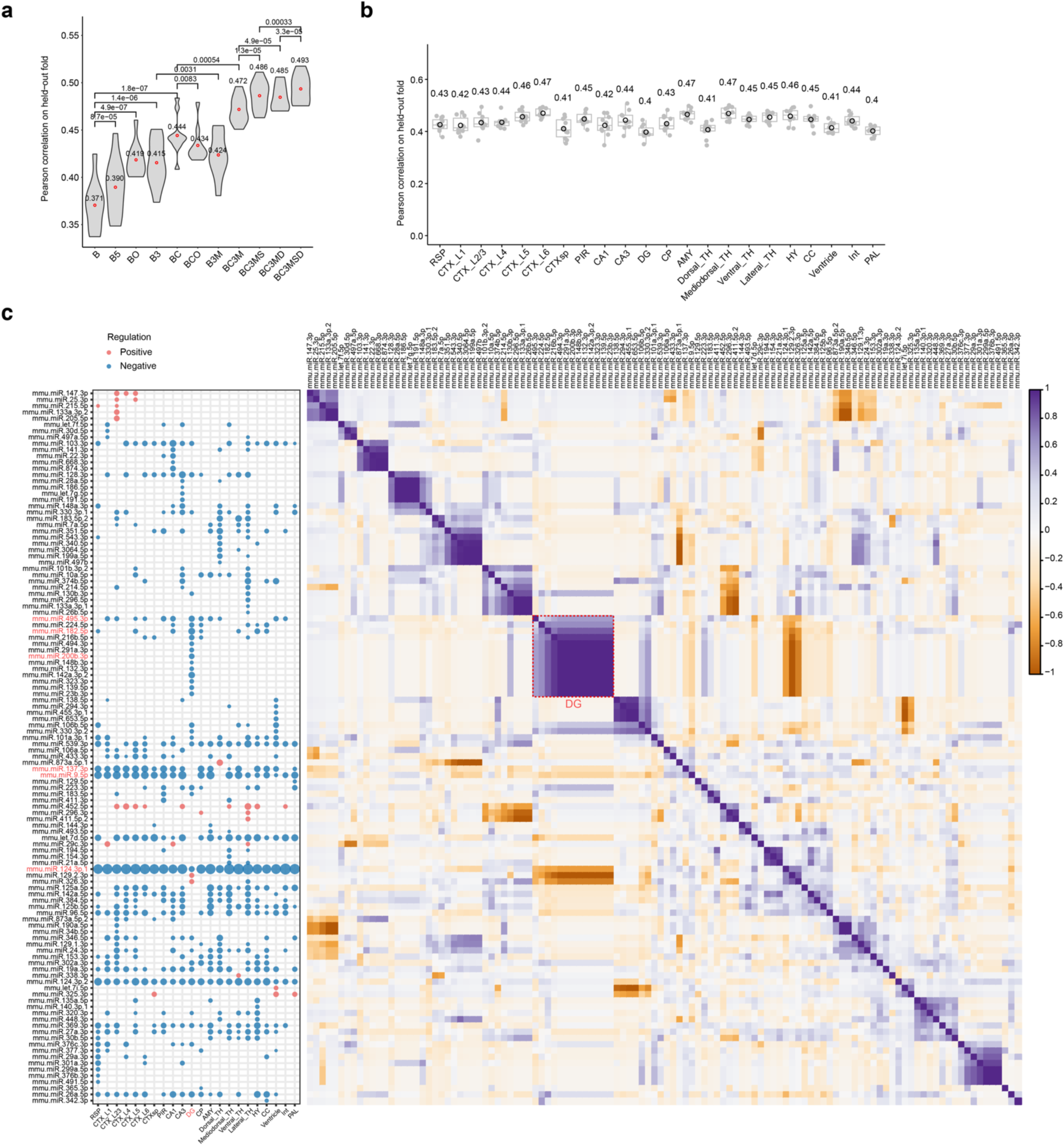
Evaluation of machine learning regression models for spatially resolved mRNA half-life prediction in mouse brains and LASSO modeling of regulatory effects of miRNAs in regulating in vivo mRNA stability across brain regions. **a.** Violin plots showing the performance of trained models in cortical regions (combined CTX domains) on each of 10 held-out folds of data. Number of the input genes: 15,754. The model evaluation was performed similarly as in **Supplementary** Fig. 15b. **b.** Box plots showing the performance of predictive modeling analysis across 22 brain regions on each of 10 held-out folds of data. Number of the input genes: RSP, 13,726; CTX_L1, 14,236; CTX_L2/3, 14,696; CTX_L4, 14,681; CTX_L5, 14,959; CTX_L6, 15,265; CTX_sp, 12,347; PIR, 14,100; CA1, 14,330; CA3, 13,831; DG, 13,329; CP, 14,175; AMY, 14,150; Dorsal_TH, 13,427; Mediodorsal_TH, 14,211; Ventral_TH, 13,675; Lateral_TH, 14,493; HY, 14,414; CC, 14,783; Ventricle, 14298; Int, 13,380; PAL, 12,320. **c.** The dot plot (left) and correlation matrix (right) of 114 miRNA features with non-zero coefficients across 22 spatial regions in P7 mouse brain sections. Dot plots showing their regulatory effect coefficients predicted by the optimized machine learning model (left). The magnitude and directionality of effect coefficients are indicated by the dot sizes and colors (red: positive regulators of mRNA stability; blue: negative regulators), respectively. Heat map showing the Pearson correlation matrix between these miRNA features (right). Brain region abbreviations: RSP, retrosplenial area; CTX, cortex; CTXsp, cortical subplate; PIR, piriform area; CA, cornu ammonis; DG, dentate gyrus; CP, caudoputamen; AMY, amygdalar nucleus; TH, thalamus; HY, hypothalamus; CC, corpus callosum; Int, internal capsule; PAL, pallidum.

**Extended Data Fig. 10.**
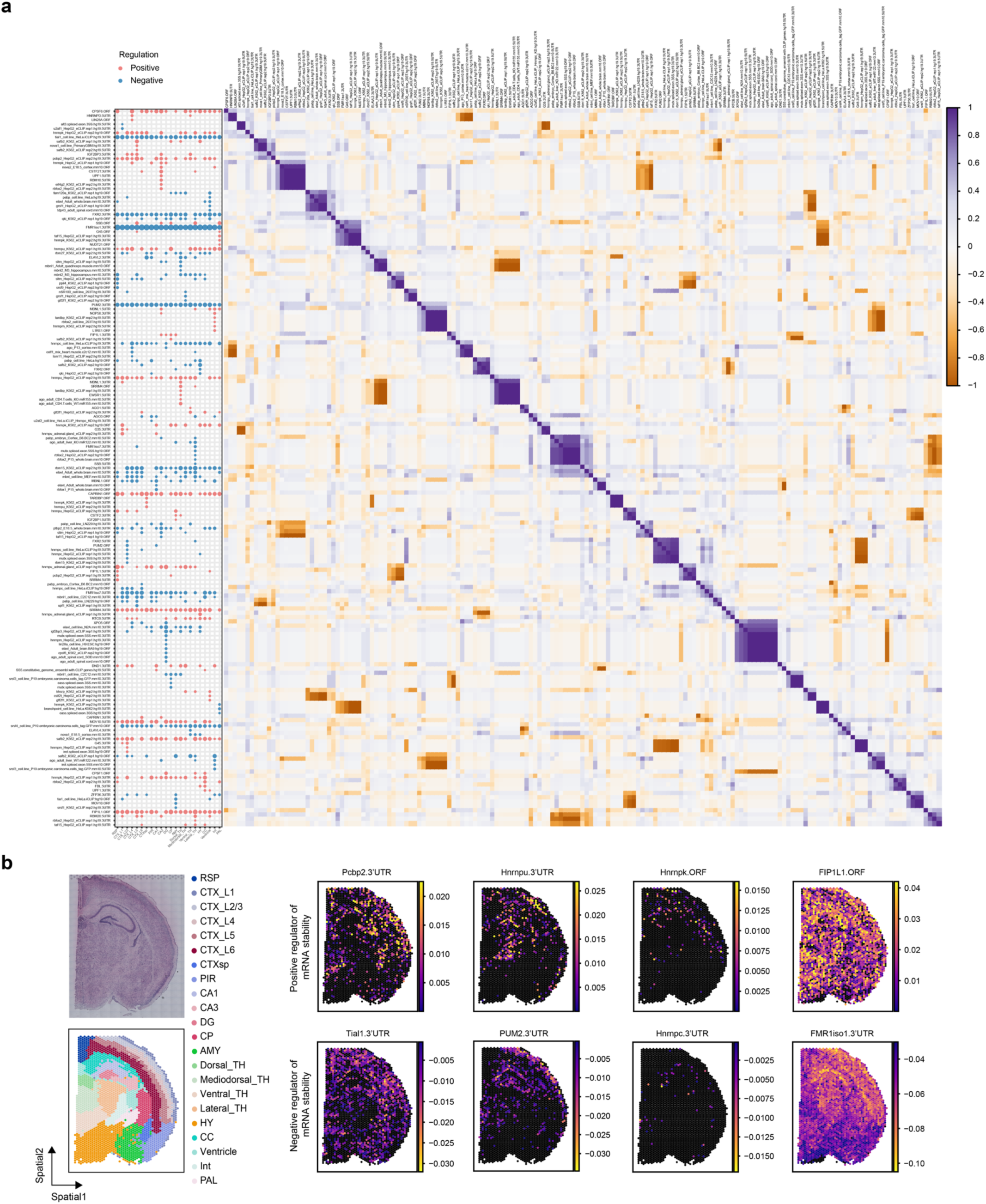
LASSO modeling of spatially resolved RNA half-life profiles reveals regulatory effects of RBPs in regulating *in vivo* mRNA stability across brain regions. **a.** The dot plot (left) and correlation matrix (right) of 167 RBP features with non-zero coefficients across 22 spatial regions in P7 mouse brain sections. Dot plots showing their regulatory effect coefficients predicted by the optimized machine learning model (left). The magnitude and directionality of effect coefficients are indicated by the dot sizes and colors (red: positive regulators of mRNA stability; blue: negative regulators), respectively. Heat map showing the Pearson correlation matrix between these RBP binding features (right). **b.** Spatial mapping of regulatory effect coefficients for eight exemplary RBP features. The top panels show four features with stabilizing effects (Pcbp2.3’UTR, Hnrnpu.3’UTR, Hnrnpk.ORF, and FIP1L1.ORF), while the bottom panels show four features with destabilizing effects (Tial1.3’UTR, Hnrnpc.3’UTR, PUM2.3’UTR, and FMR1-iso1.3’UTR). The H&E staining image (top) and spatial annotation (bottom) of the same mouse brain section (treated-rep1 in Fig. 2b) are provided in the left panels.

