## Supplemental Figures and Notes for "Spatial mapping of RNA turnover kinetics and regulatory landscapes of mRNA stability in the mammalian brain"

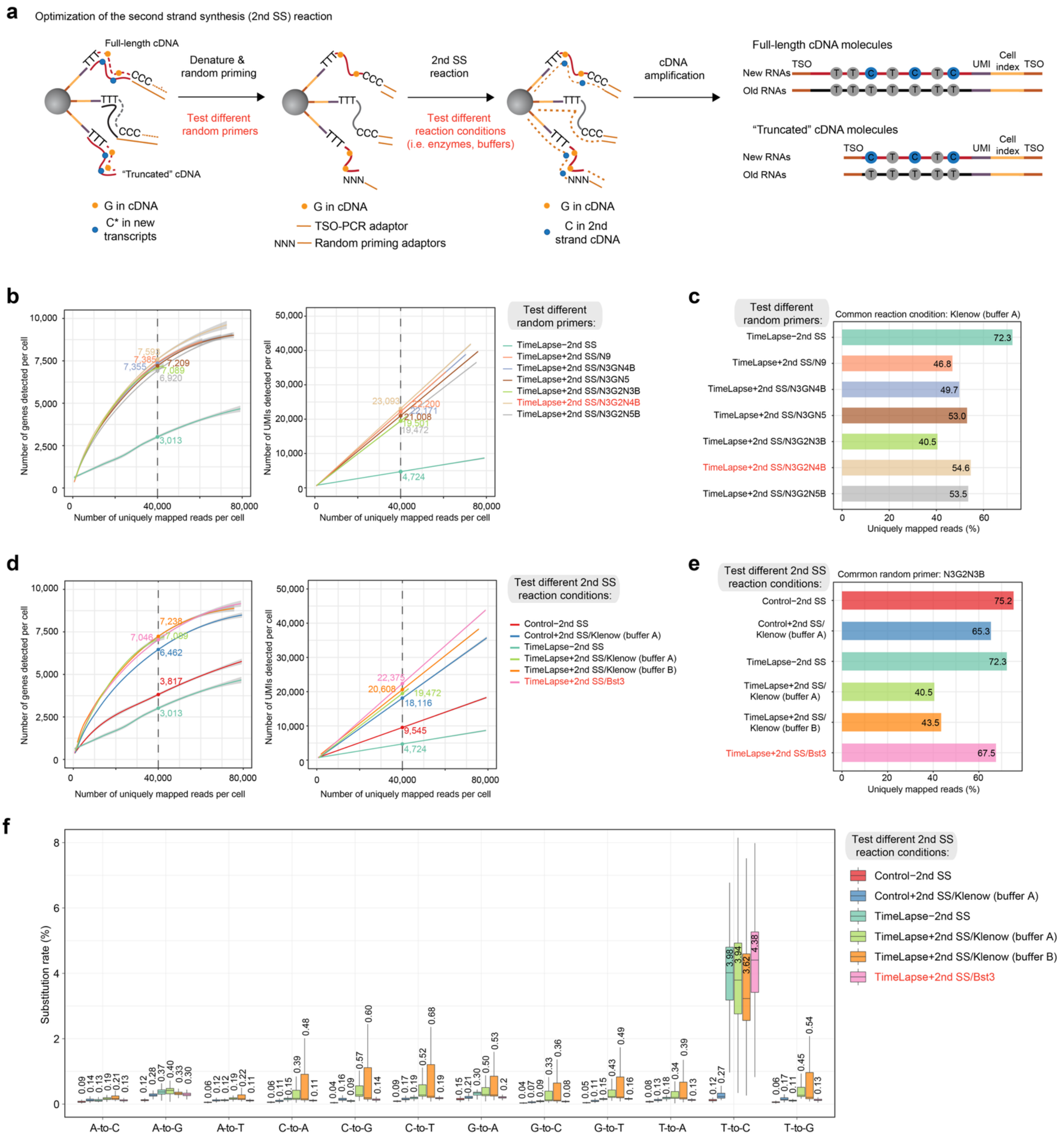


**Supplementary Fig. 1 | Development and benchmark of scNT-seq2 to enhance performance of droplet-based time-resolved single-cell RNA-seq.**

- **a.** Schematic representing the experimental strategy for optimizing the 2$^{nd}$ SS reaction in scNT-seq2.
- **b.** Fitted line plots illustrating library complexity by comparing genes (left) or UMIs/transcripts (right) detected per cell as a function of aligned reads per cell across experiments with various random primers. This set of experiments used a previously reported Klenow-based reaction mixture [1], with different random primer sequences (provided in **Supplementary Table 1**). All experiments were performed using the same batch of *in vitro* 4sU-labeled K562

cells (100 µM, 4 hours). Estimated numbers of genes or UMIs detected per cell at matching sequencing depth (40,000 reads per cell) for different experiments are shown. The shaded regions depict 95% confidence intervals.

**c.** Bar graph showing the fraction of uniquely mapped reads for the experiments in **(b)**.

**d.** Fitted line plots illustrating library complexity by comparing genes (left) or UMIs/transcripts (right) detected per cell as a function of aligned reads per cell across various enzymatic reaction conditions. Either reaction buffer A [1] or B [2] was used with Klenow Fragment (3'->5' exo-); *Bst* 3.0 DNA polymerase (denoted as Bst3) was identified as the more optimal condition in this study and was used in scNT-seq2. All experiments were performed using the same batch of *in vitro* 4sU-labeled K562 cells (100 µM, 4 hours). Estimated numbers of genes or UMIs detected per cell at matching sequencing depth (40,000 reads per cell) for different experiments are shown. The shaded regions depict 95% confidence intervals.

**e.** Bar graph showing the fraction of uniquely mapped reads from the experiments in **(d)**.

**f.** Box plot comparing nucleotide substitution rates in 4sU-labeled K562 cells from experiments in **(d)**. Cell numbers for each experimental conditions: control-2$^{nd}$ SS, 595; control+2$^{nd}$ SS/Klenow (buffer A), 602; TimeLapse-2$^{nd}$ SS, 364; TimeLapse+2$^{nd}$ SS/Klenow (buffer A), 600; TimeLapse+2$^{nd}$ SS/Klenow (buffer B), 603; TimeLapse+2$^{nd}$ SS/Bst3, 599.

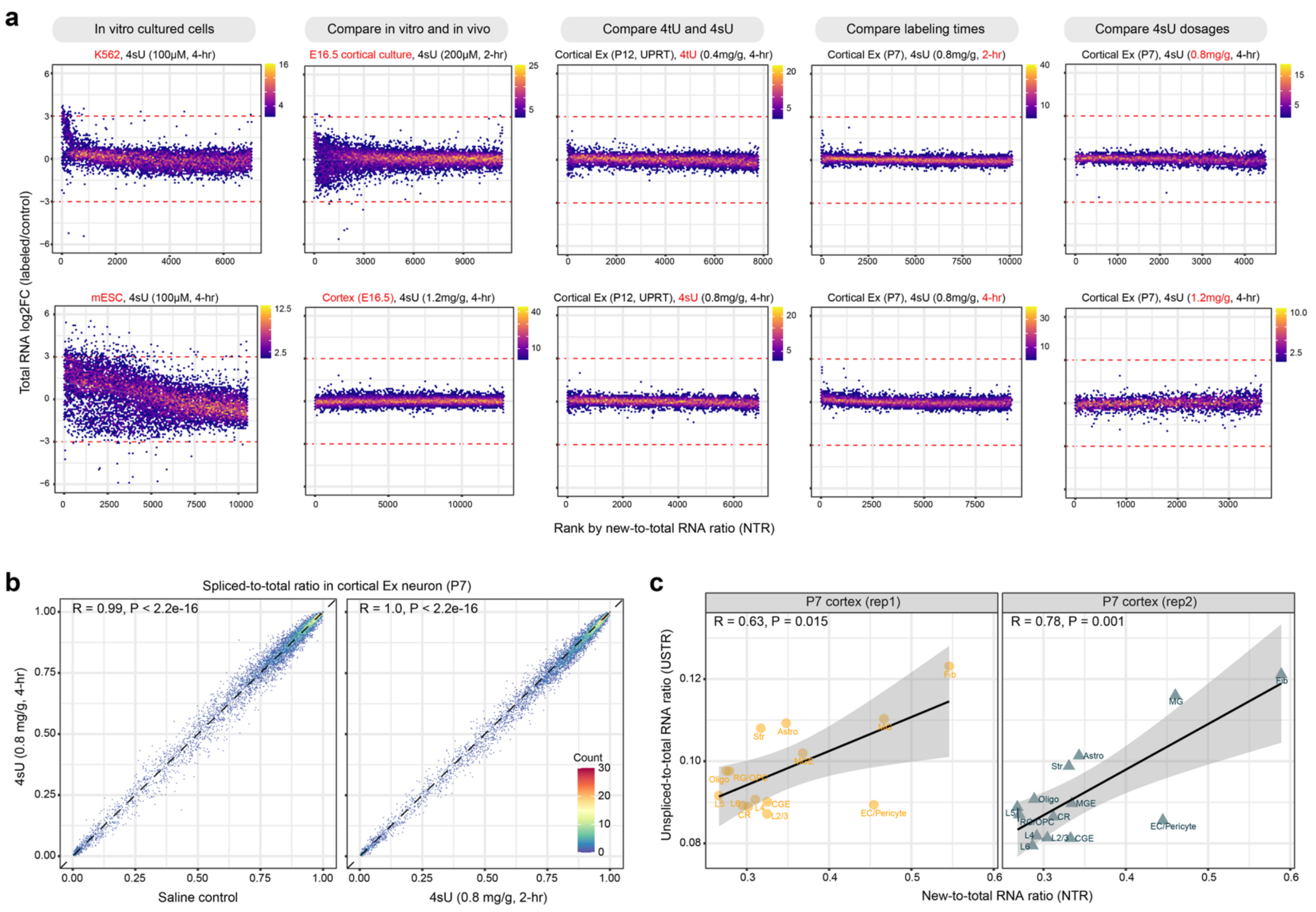


**Supplementary Fig. 2 | Evaluation the effect of *in vivo* transgenesis-free metabolic RNA labeling on global transcription and splicing in wild-type mouse cortical samples.**

- **a.** Scatter plots comparing the ranks of the new-to-total RNA ratios (NTRs) of each gene against the $\log_2$ fold change of labeled samples versus corresponding control samples. NTR low genes (half-life high) are associated with lower ranks, whereas NTR high genes (half-life low) are associated with higher ranks. From left to right: *in vitro* labeled cultured cells (K562 and mESC), comparison of matched *in vitro* and *in vivo* samples (E16.5 cortical cells), comparison of 4tU and 4sU labeling (P12 cortical Ex neurons), comparison of 4sU labeling with different labeling time (P7 cortical Ex neurons), and comparison of 4sU labeling with different dosages (P7 cortical Ex neurons). The sample type (cell-type or tissue type/age), labeling concentration and time are also indicated on the top of the panel. The red dash lines (based on *in vitro* cultured cells as a benchmark) depict the limits for identifying potentially affected genes by 4sU labeling. Setting of the limits are empirically guided by data variability from the relatively short *in vitro* 4sU labeling experiment such as E16.5 cortical primary cells (i.e. 200μM of 4sU for 2h). The color density represent gene counts.
- **b.** Scatter plots comparing global RNA splicing (measured by spliced-to-total RNA ratio) between 4-hr labeling (y-axix) and saline control (left panel, x-axis) or 2-hr labeling (right panel, x-axis) in P7 cortical Ex neurons. The color density represent gene counts.
- **c.** Scatter plots comparing NTRs (x-axis, from RNA labeling) and USTRs (y-axis, from RNA splicing) of commonly detected genes in cortical cell-types from two P7 mice (left: replicate 1; right: replicate 2).

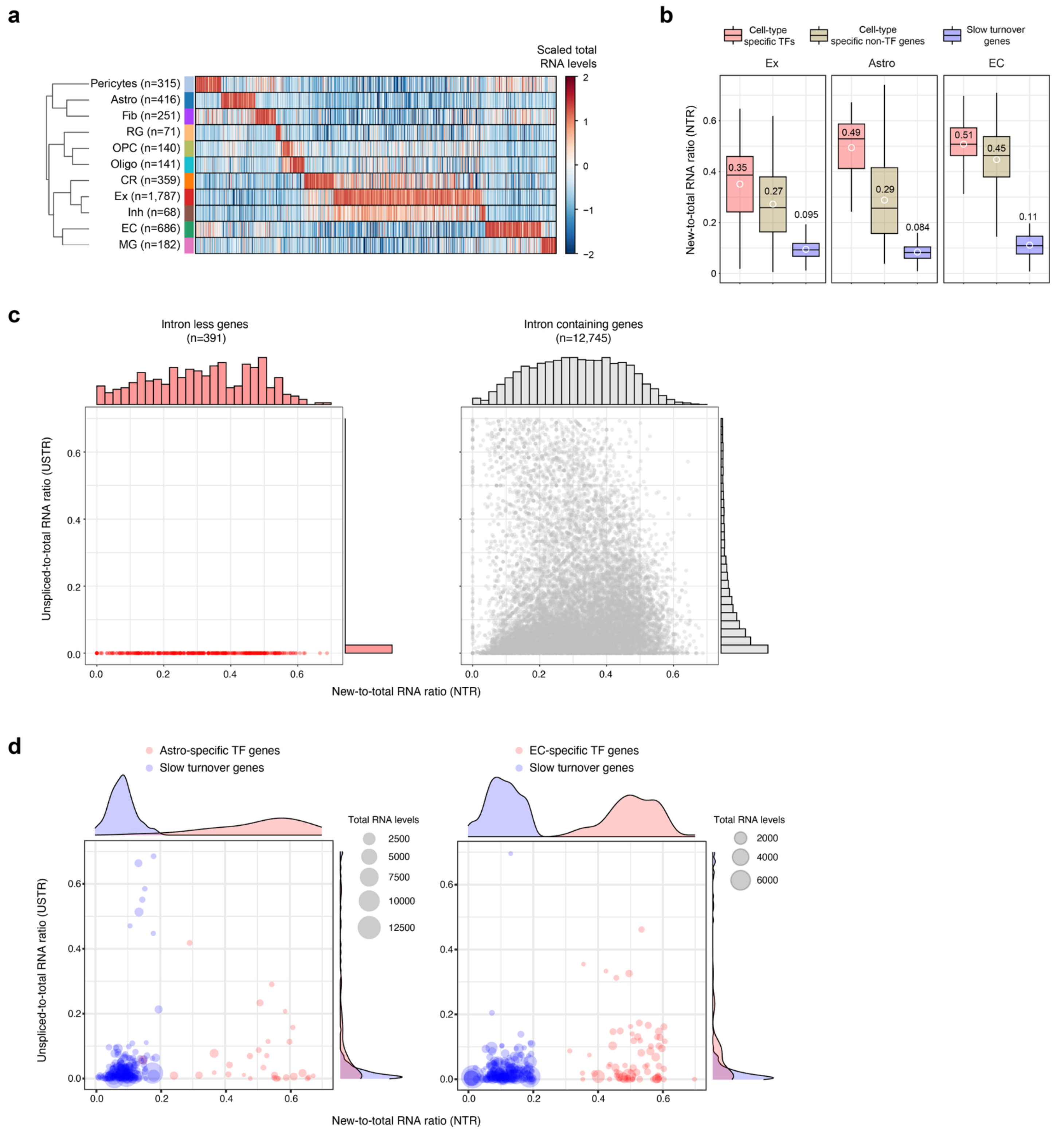


**Supplementary Fig. 3 | *In vivo* time-resolved scRNA-seq reveals RNA turnover dynamics in the mouse cortex.**

**a.** Heatmap displaying the scaled expression levels of 4,416 cell-type specific genes across major cortical cell-types in P7 cortex. The number of cell-type specific genes detected within each cell type is denoted in brackets.

**b.** Box plot displaying cell-type-specific NTRs for various gene groups. Number of genes: cell-type-specific TF – Ex (n=124), Astro (n=31), and EC (n=68); cell-type-specific non-TF genes – Ex (n=1,663), Astro (n=385), and EC (n=618); Broadly expressed, slow turnover house-keeping genes – Ex/Astro/EC (n=269).

**c.** Scatter plots showing NTRs (x-axis, from RNA labeling) and USTRs (y-axis, from RNA splicing) for gene groups with distinct genes structures (intron-less genes – red, without any intron; intron-containing genes – gray, with at least one intron) in cortical Ex neurons. Only expressed genes (transcript counts>5) are shown, and the number of genes for each group is indicated. The marginal histogram plots depict the distributions of NTRs (x-axis) and USTRs (y-axis) for each gene group.

**d.** Scatter plots showing new-to-total RNA ratio (NTR derived from RNA labeling) and unspliced-to-total RNA ratio (USTR from RNA splicing) for cell-type specific TFs (red) and broadly expressed slow turnover genes (blue) in cortical astrocytes (Astro: left panels) and endothelial cells (EC: right panels). The size of circles represent total RNA levels for each gene (CPM: counts per million reads). The marginal density plots depict the distributions of NTRs (x-axis) and UTRs (y-axis) for two gene groups.

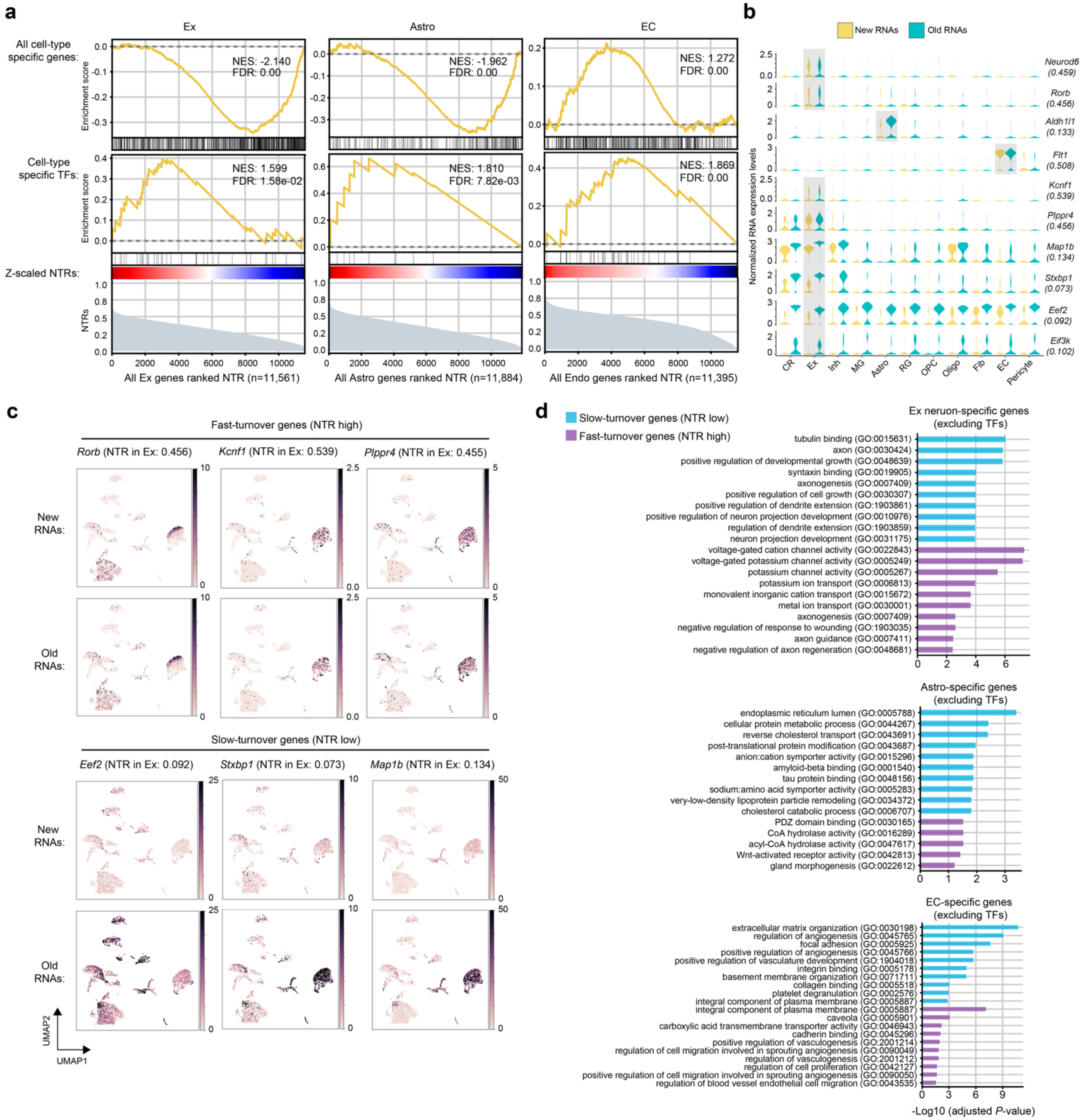

**Supplementary Fig. 4 | *In vivo* time-resolved scRNA-seq reveals cell-type-specific genes with distinct RNA turnover dynamics in postnatal mouse cortex.**

- **a.** Gene set enrichment analysis (GSEA) of cell-type specific genes (top panels) and cell-type specific TFs (middle panels) among three distinct cortical cell types. Genes were ranked based on their z-scaled labeled-to-total RNA ratio (NTR, bottom panels). Each panel specifies the corresponding false discovery rate (FDR) and normalized enrichment score (NES). Cell type abbreviations: Ex, excitatory neurons; Astro, astrocytes; EC, endothelial cells.

**b.** Violin plots illustrating the expression levels of both new and old RNAs (log2 transformation of CP10K) of 10 representative genes across major cortical cell-types. RNA turnover rates (i.e. NTRs) of each gene in specific cell-type (highlighted by gray shade) are shown below gene names.

**c.** Feature plots highlighting the new and old RNA levels (quantified as CP10K) of six exemplary genes. Among them, three exhibit high NTR (*Rorb*, *Kcnf1* and *Plppr4*), while the other three show low NTR (*Map1b, Stxbp1* and *Eef2*) in Ex neurons. RNA turnover rates (i.e. NTRs) of each gene in Ex neurons are shown next to gene names.

**d.** Enrichment analysis highlighting Gene Ontology (GO) terms among genes classified as slow turnover (bottom 30%) versus those with fast turnover (top 30%) across three cortical cell-types.

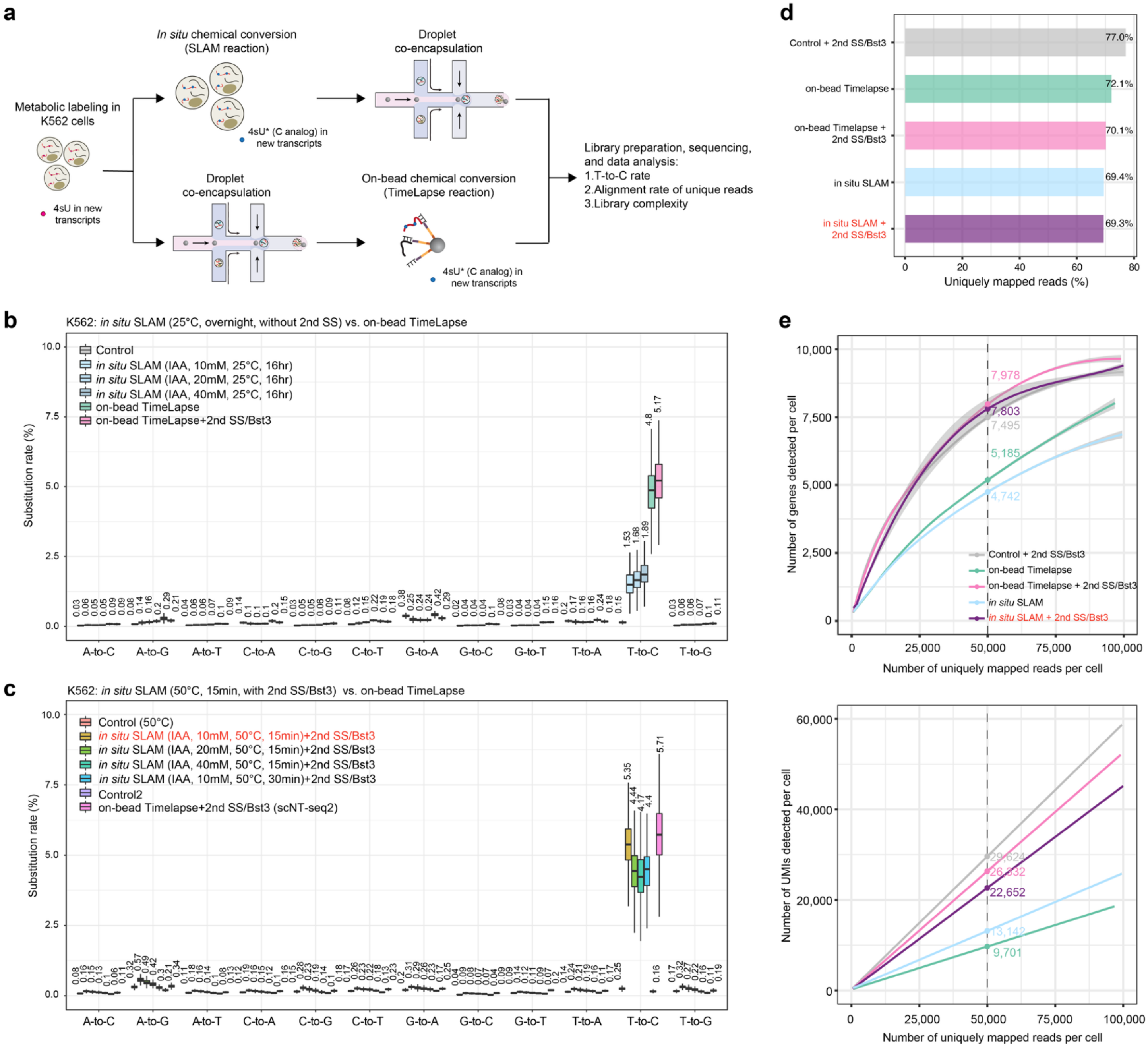


**Supplementary Fig. 5 | Optimizing and benchmarking *in situ* chemical conversion in cultured cells.**

- **a.** Overview of the experimental workflow for *in situ* chemical conversion optimization. The same batch of 4sU-labeled K562 cells was subjected to different chemical conversion conditions. The on-bead TimeLapse chemical conversion reaction (scNT-seq2) was employed using the same batch of input cells as a positive control for *in situ* SLAM chemical conversion.
- **b.** Box plot illustrating nucleotide substitution rates in 4sU-labeled K562 cells across experiments with varying IAA concentrations, incubated overnight (16hr) at room temperature (25°C). On-bead TimeLapse reaction used in scNT-seq2 was used as a positive control. Number of cells for each experimental condition: Control, 1,001; *in situ* SLAM (10mM IAA, 25°C/16hr), 920; *in situ* SLAM (20mM IAA, 25°C/16hr), 1,003 cells; *in situ* SLAM (40mM IAA, 25°C/16hr), 1,002; on-bead TimeLapse, 780; on-bead TimeLapse+2ndSS/Bst3, 1,002.
- **c.** Box plot showing nucleotide substitution rates in 4sU-labeled K562 cells across various experiment conditions with optimized second strand synthesis reactions. On-bead TimeLapse reaction used in scNT-seq2 was used as a positive control. Number of cells for each experimental condition: Control-50°C, 1,001; *in situ* SLAM (10mM IAA, 50°C/15min)+2ndSS/Bst3, 1,001; *in situ* SLAM (20mM IAA, 50°C/15min)+2ndSS/Bst3, 1,003; *in situ* SLAM

(40mM IAA, 50°C/15min)+2ndSS/Bst3, 1,002; *in situ* SLAM (10mM IAA, 50°C/30min)+2ndSS/Bst3, 1,001; Control-2, 1,002; on-beads TimeLapse+2ndSS/Bst3, 1,001. The final optimized condition is highlighted in red.

**d.** Bar graph depicting the percentage of uniquely mapped reads across the samples from the listed experimental conditions. The final optimized condition is highlighted in red.

**e.** Fitted line plots illustrating library complexity by comparing genes (upper panel) or UMIs/transcripts (lower panel) detected per cell as a function of aligned reads per cell across experiments with different conditions, including *in situ* SLAM and on-bead TimeLapse conversion reactions (with or without the 2nd SS/Bst3 reaction). Estimated numbers of genes or UMIs detected per cell at matching sequencing depth (50,000 reads per cell) for different experiments are shown. The shaded regions depict 95% confidence intervals. The optimized condition is highlighted in red.

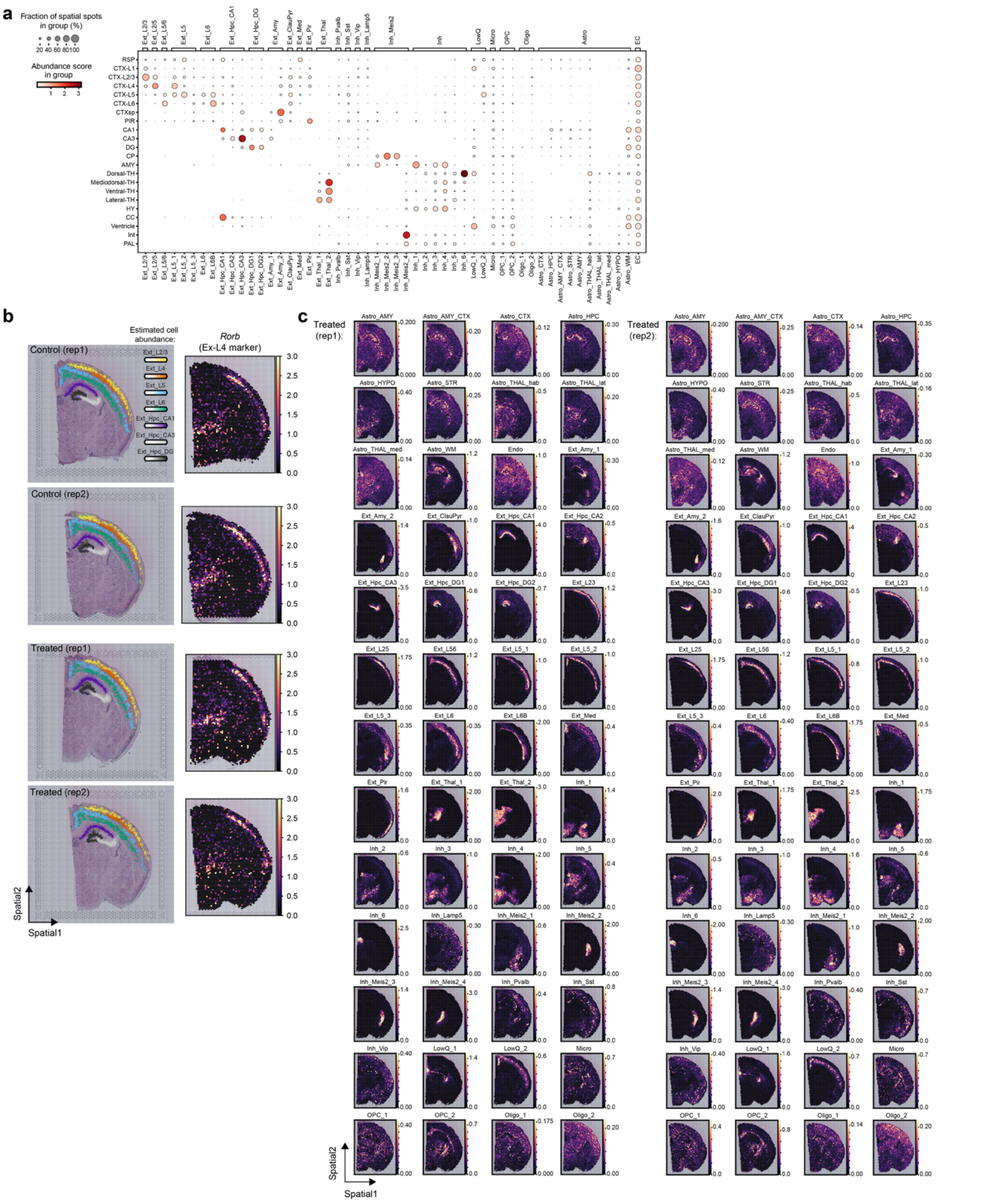

**Supplementary Fig. 6 | Validation and deconvolution of cell type composition in distinct spatial brain regions using Cell2location.**

**a.** Dot plots displaying the cell type composition within annotated spatial regions by integrated analysis of spatial clusters derived from spatial NT-seq (y-axis) and neuronal/non-neuronal cell-types (x-axis) determined by a published whole brain single-nucleus RNA-seq dataset using Cell2location [3]. Dot size represents the fraction of spatial spots in an annotated spatially resolved brain region associated with a specific cell type, while the color intensity indicates the cell abundance score of that cell type within the region. Brain region abbreviations: RSP, retrosplenial area; CTX, cortex; CTXsp, cortical subplate; PIR, piriform area; CA, cornu ammonis; DG, dentate gyrus; CP, caudoputamen; AMY, amygdalar nucleus; TH, thalamus; HY, hypothalamus; CC, corpus callosum; Int, internal capsule; PAL, pallidum.

**b.** Visualization of cell abundance score (indicated by color intensity) for seven representative excitatory (Ext) neuronal subtypes within cortical and hippocampal (Hpc) regions across two control and two treated tissue sections in **Fig. 2b**. The normalized spatial expression levels of *Rorb* gene (cortical layer 4 exitatory neuron marker) are shown in right panel.

**c.** Cell abundance scores of 52 annotated neuronal and non-neuronal cell-types across two *in situ* chemically “treated” mouse brain sections.

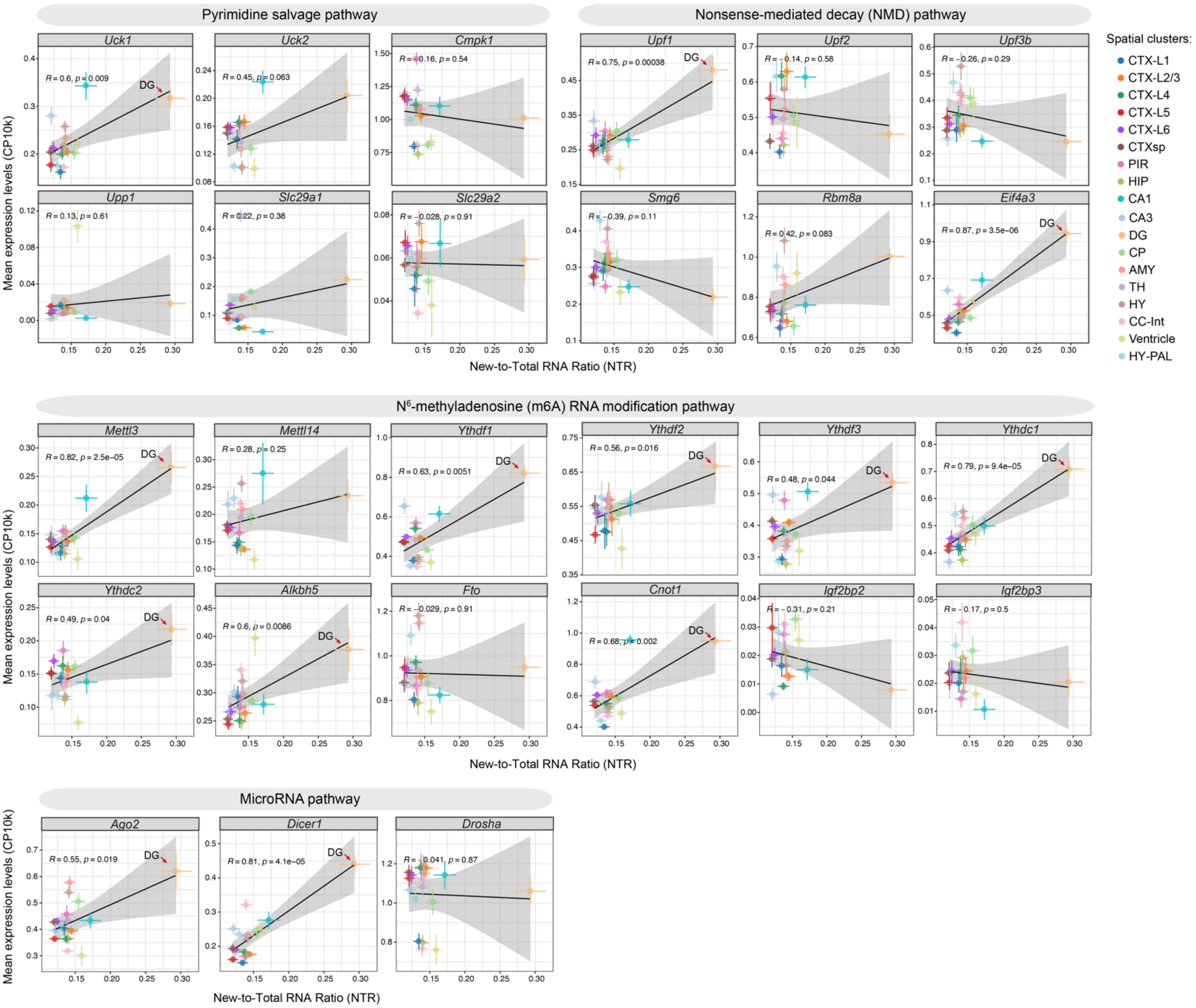


**Supplementary Fig. 7 | Relationship between spatial RNA turnover rates and expression levels of key genes involved in pyrimidine salvage and major RNA decay pathways in adult mouse brain regions.**

Scatter plots showing the correlation within P56 (spatial cluster annotations are from **Fig. 3a**) coronal mouse brain sections (12 sections from six P56 mice in **Supplementary Fig. 12a**) between spatial domain specific RNA turnover (NTRs in x-axis) and normalized total RNA expression levels (CP10k) of genes functionally related to the **pyrimidine salvage pathway** [4] (***Uck1/2***: uridine/cytidine kinase 1 and 2 (*Uck1/2*), two ribonucleoside kinases in the the pyrimidine salvage pathway that mediate the rate-limiting step of 4sU activation; ***Cmpk1***: cytidine/uridine monophosphate kinase 1, an essential enzyme for pyrimidine utilization in all cells as both *de novo* biosynthesis and intracellular/extracellular salvage requires Cmpk1 activity; ***Upp1***: uridine phosphorylase, an enzyme that degrade uridine; ***Slc29a1/2***: Na+-independent SLC29 family of equilibrative nucleoside transporters, the main uridine transporters that mediate 4sU uptake into mammalian cells), the **NMD RNA decay pathway** [5] (core NMD pathway components - ***Upf1***: Up-frameshift protein 1 (*Upf1*) is an ATP-dependent RNA helicase that plays a crucial role in various mRNA decay pathways, especially in the NMD pathway; ***Upf2***: Up-frameshift protein 2 (*Upf2*), an adaptor protein that interacts with eukaryotic release factors (eRFs) and Upf3b (see below); ***Upf3b***: Up-frameshift protein 3B (*Upf3b*), a RNA binding protein that mediate the interactions between Upf1/Upf2, exon junction complex (EJC) and eRFs; ***Smg6***: suppressor with morphological effect on genitalia-6 (*Smg6*), an endonuclease that cleaves NMD target mRNAs; EJC factors - ***Rbm8a***: RNA binding motif protein 8A; ***Eif4a3***: eukaryotic translation initiation factor 4A3), the **RNA m6A methylation pathway** [6,7] (***Mettl3/14***:

methyltransferase-like 3 or 14, the stable heterodimer core complex Mettl3-Mettl14 of a multi-subunite m6A writer enzyme complex; ***Ythdf1-3*** and ***Ythdc1/2***: YTH domain-containing direct m6A reader proteins, and as the major RNA decay-inducing reader protein, Ythdf2 can directly interact with ***Cnot1*** and recruits the CCR4-NOT deadenylase complex to m6A-modified RNAs, leading to the degradation of RNAs; ***Igf2bp2/3***: insulin-like growth factor 2 mRNA-binding proteins, two m6A reader proteins that stabilize RNAs; ***Alkbh5*** and ***Fto***: two m6A eraser enzymes by oxidatively demethylating m6A on RNA), and the **microRNA mediated RNA decay pathway** [8] (***Ago2/Dicer1/Drosha***: key proteins involved in microRNA biogenesis and processing). Both the vertical (normalized gene expression) and honrizontal (spatial domain specific NTRs) error bars represent standard errors (n=12 sections). The Pearson's correlation coefficient R and *P*-value are indicated. The genes showing significant correlation (*P*<0.05) are highlighted with red arrows.

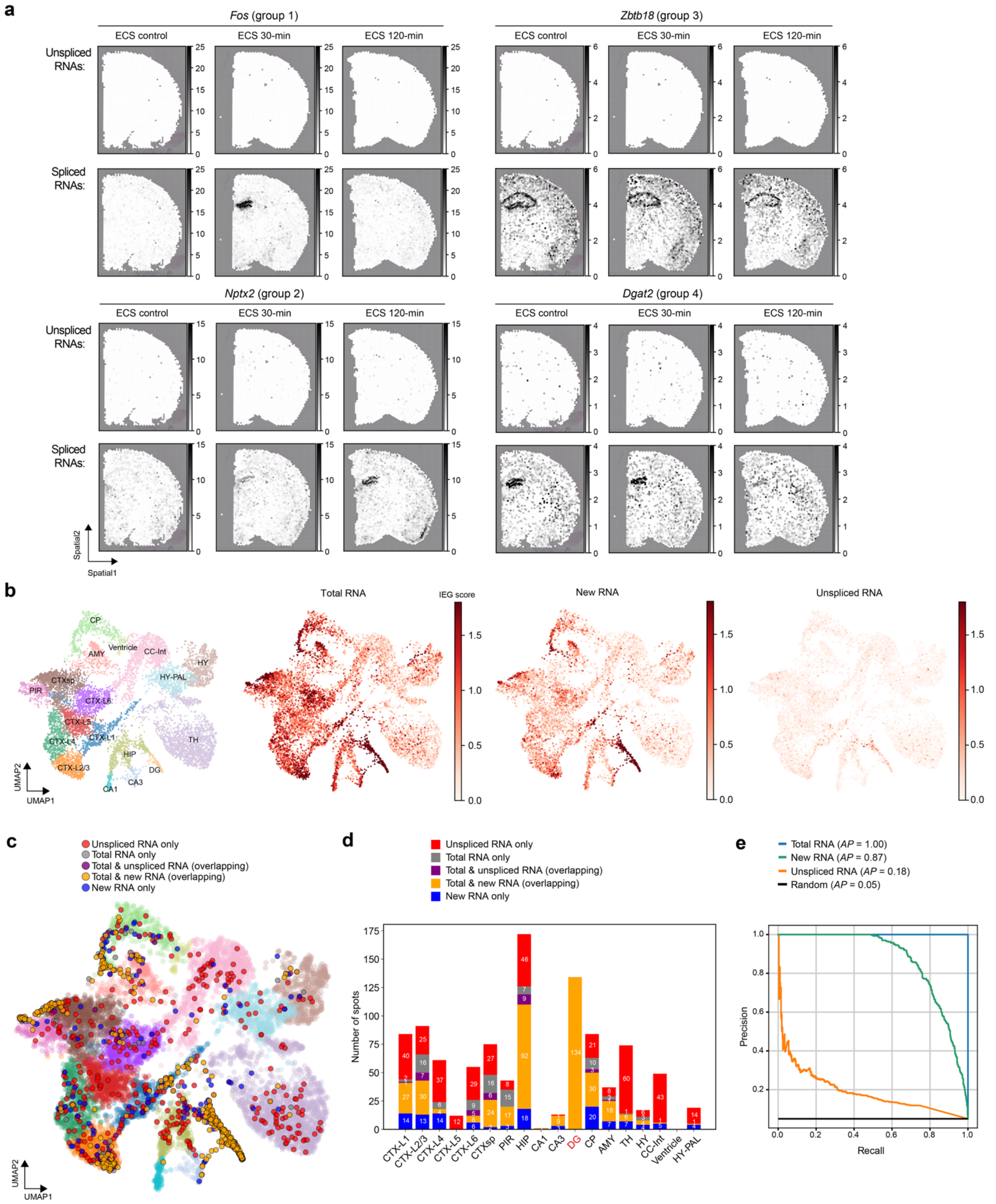


**Supplementary Fig. 8 | Benchmarking RNA labeling versus splicing based analyses of ARG dynamics in spatial NT-seq data.**

**a.** Panels displaying the normalized spatial expression (in CP10K) of unspliced (upper panel) and spliced (lower panel) RNAs for four representative genes (same as in **Fig. 3g**) from each ARG group using a representative spatial NT-seq dataset (section #2 from mouse #2/4/6).

**b.** Feature plot showing IEG scores across spatial spots in spatial NT-seq data (ECS 30 minutes). Spots are colored by annotated brain region and by IEG scores derived from total, new, and unspliced RNA. As previously described [9], IEG scores were calculated as the mean CP10k-normalized expression across the curated IEG panel: *Arc*, *Bdnf*, *Btg2*, *Fos*, *Fosl2*, *Homer1*, *and Npas4.*

**c.** UMAP visualization of trapped cells from spatial spots 30 minutes post-ECS. Top 5% of IEG scores from total, new, or unspliced RNA were classified as "trapped" ($IEG^{high}$). UMAP embeddings display exclusive and overlapping categories: total only (gray), unspliced only (red), new only (blue), total & unspliced (purple), total & new (orange), with all other spots shown in background (colored coded by brain regions).

**d.** Bar plot depicting the number of spatial spots per brain region classified as "trapped" ($IEG^{high}$) by each category.

**e.** Precision–recall benchmarking of labeling- (new RNA) and splicing-based (unspliced RNA) analysis in recovering spatial spots classified as "trapped" ($IEG^{high}$) when compared to total RNA IEG scores.

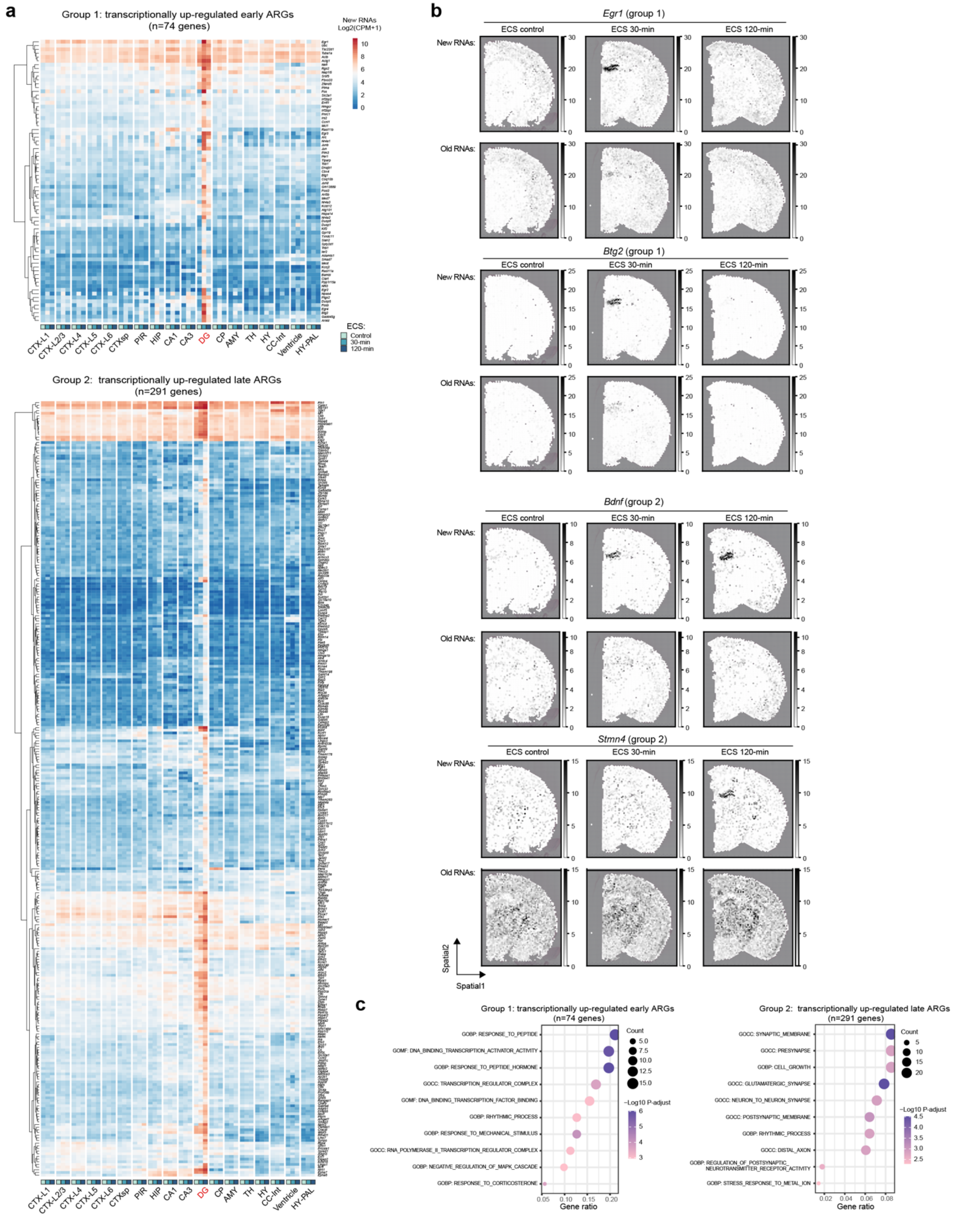


**Supplementary Fig. 9 | Spatial NT-seq reveals ECS-induced dynamics of transcriptionally up-regulated new RNAs.**

**a.** Heat maps illustrating the ECS-induced dynamics of new RNAs within two groups of transcriptionally up-regulated ARGs (group 1: early ARGs on the top; group 2: late ARGs on the bottom) across 18 brain regions.

**b.** Panels displaying the normalized spatial expression (in CP10K) of new and old RNAs for four representative genes across three time points (ECS control, 30-, and 120-min) using a representative spatial NT-seq dataset (section #2 from mouse #2/4/6). *Egr1* and *Btg2* are examples of transcriptionally up-regulated early ARGs (group 1), whereas *Bdnf* and *Stmn4* are examples of transcriptionally up-regulated late ARGs (group 2).

**c.** Gene Ontology enrichment analysis reveals significant biological processes associated with early ARGs (group1) and late ARGs (group2). Dot size corresponds to the number of genes in each category, while the color represents the log-scaled, adjusted *P*-values (Hypergeomtric test).

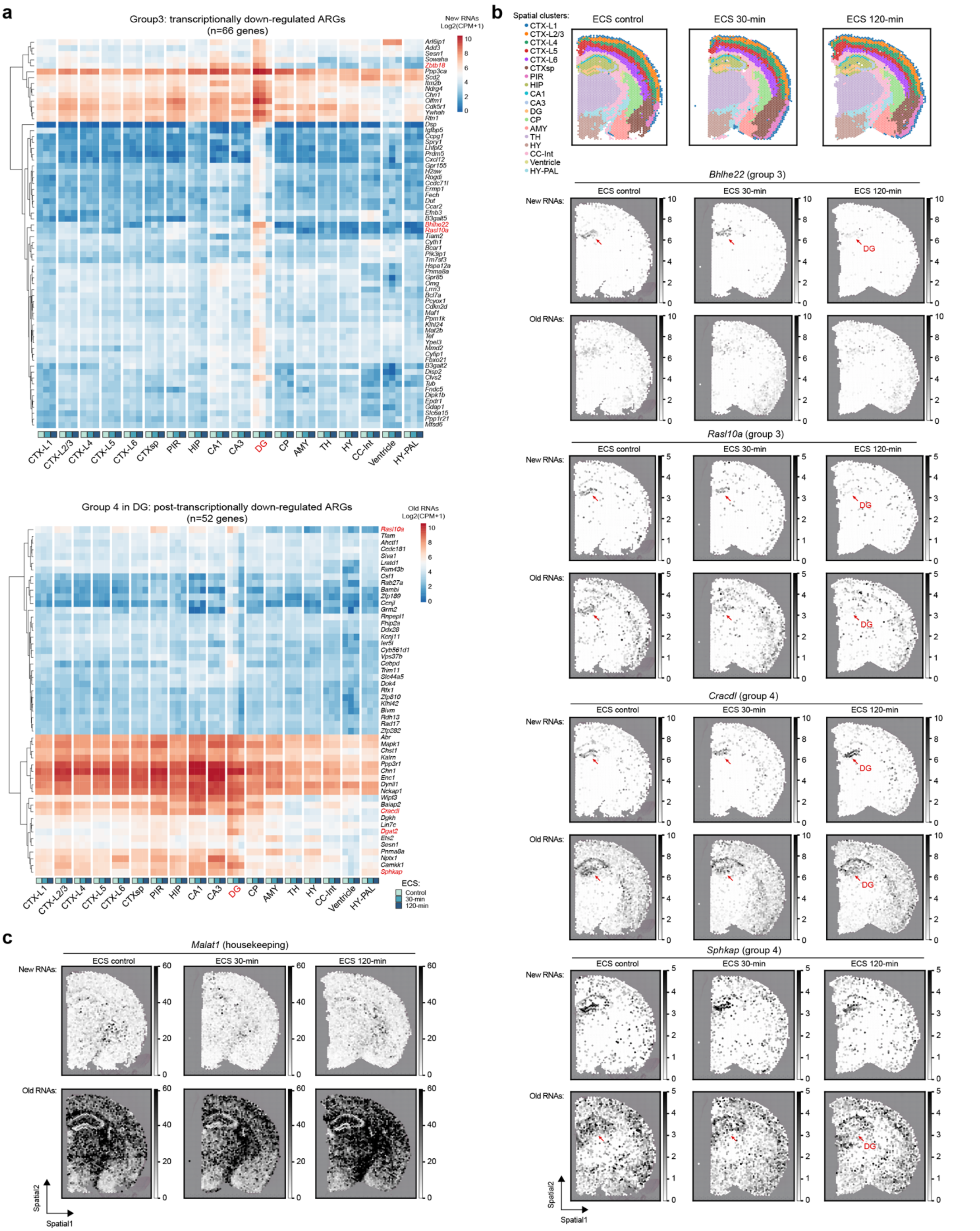


**Supplementary Fig. 10 | Spatial NT-seq reveals ECS-induced transcriptional or post-transcriptional down-regulation of ARGs.**

**a.** Heat maps illustrating the ECS-induced dynamics of new RNAs for transcriptionally down-regulated ARGs (group 3, top) or old RNAs for post-transcriptionally down-regulated ARGs (group 4, bottom) across 18 brain regions.

**b.** Panels displaying the normalized spatial expression (in CP10K) of new and old RNAs for four representative genes across three time points (ECS control, 30-, and 120-min) using a representative spatial NT-seq dataset (section #2 from mouse #2/4/6). *Bhlhe22* and *Rasl10a* are examples of transcriptionally down-regulated genes (group 3), whereas *Cracdl* and *Sphkap* are examples of post-transcriptionally down-regulated genes (group 4).

**c.** Panels displaying the normalized spatial expression (in CP10K) of new and old RNAs for *Malat1*, a widely expressed non-coding RNA in the brain that are not linked to ECS (as a control gene for ARGs).

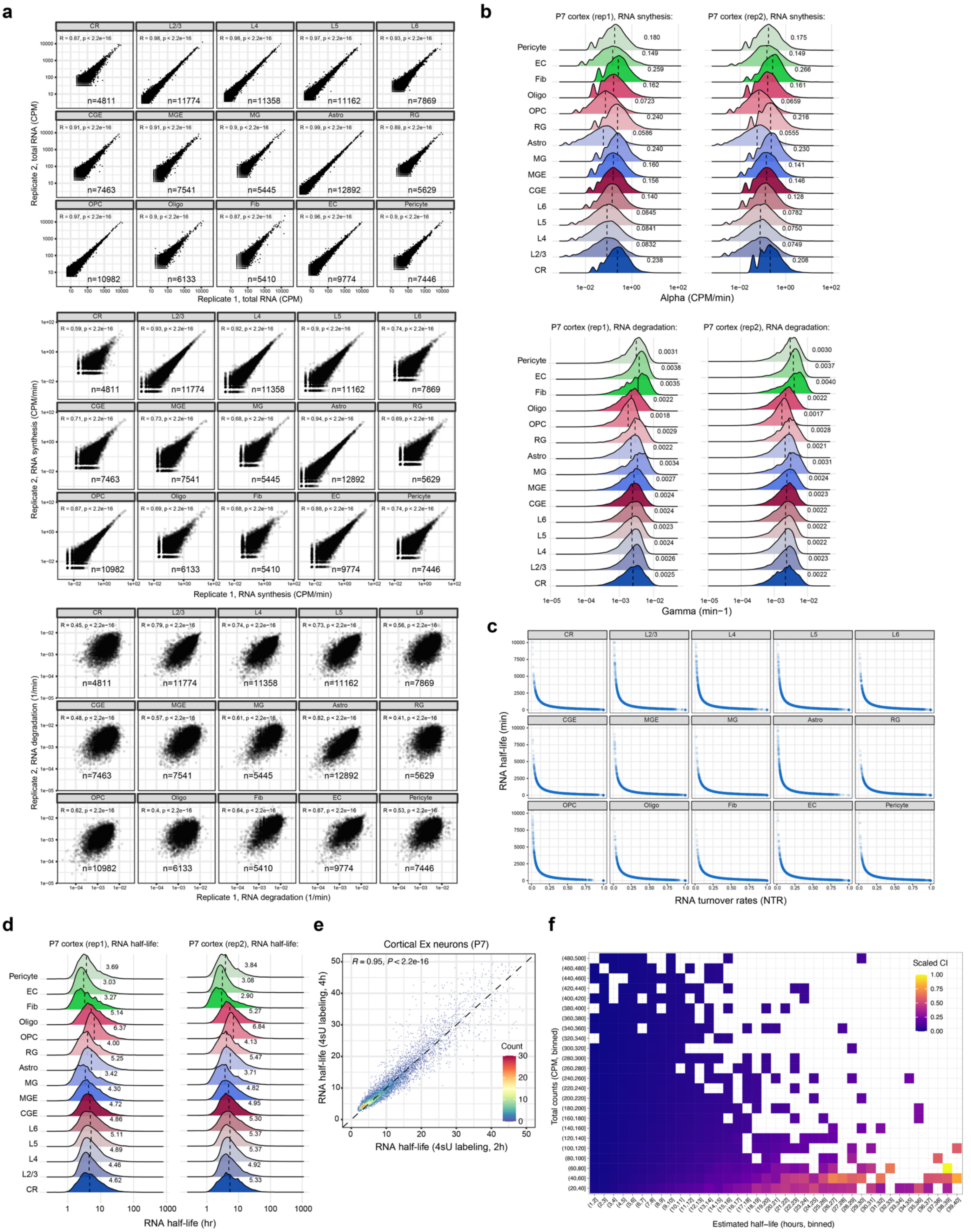

**Supplementary Fig. 11 | Validation of *in vivo* cell-type-specific RNA kinetics parameters analysis.**

**a.** Scatter plots showing Pearson's correlation for cell-type-specific RNA abundance (top), synthesis rate (alpha, middle), and degradation rate (gamma, bottom) across 15 cortical cell types between two independent biological replicates.

**b.** Density ridgeline plots depicting the distributions of RNA synthesis rate (alpha, top) and degradation rate (gamma, bottom) for 15 cortical cell types across two replicates. The mean is indicated by the dash vertical line and the values are shown on the side.

**c.** Scatter plots illustrating the inverse relationship between NTR and RNA half-lives across the 15 cortical cell types.

**d.** Density ridgeline plots showing the distributions of RNA half-lives for 15 cortical cell types between two replicates. The mean is indicated by the dash vertical line and the values are shown on the side. This analysis reveals an averaged *in vivo* RNA half-life of 4.6-hr across diverse cortical cell types (ranging from 3.1- to 6.6-hr), highly comparable to *in vitro* RNA half-life of ~4.7-hr averaged from various *in vitro* studies using single-pulse or pulse-chase metabolic RNA labeling (mouse NIH3T3: 9.9-hr [10] or 4.9-hr [11]; mouse embryonic fibroblasts: 2.7-hr, human K562: 1.6-hr [12]; mouse ESCs: 3.9-hr [13]; human B cells: 5.2-hr, mouse fibroblasts: 4.6-hr [14], and mouse primary cortical neurons: soma/cytoplasm-enriched mRNAs – 3.7-hr and neurite-enriched mRNAs – 5.6-hr [15]).

**e.** Scatter plots comparing transcriptome-wide RNA half-life (hours) between 4-hr labeling (y-axis) and 2-hr labeling (x-axis) in P7 cortical Ex neurons. Pearson's correlation coefficient ($R$=0.95) and statistical significance ($P$<$2.2 \times 10^{-16}$) are shown on the top. The color density represent gene counts.

**f.** Bootstrap resampling quantifies uncertainty in RNA half-life estimates across observed gene expression levels in cortical excitatory neurons (Ex). Half-lives (x-axis) and expression counts (CPM, y-axis) were binned, and median confidence interval (CI) widths were calculated per bin. CI widths are scaled across all bins, with color indicating relative uncertainty. This analysis revealed that genes with moderate to high expression and short to medium RNA half-life (<16 hrs) showed relatively small confidence intervals and thus have the more reliable estimates. By contrast, lowly expressed genes with longer half-life (>16 hrs) have wider confidence intervals and less reliable estimates.

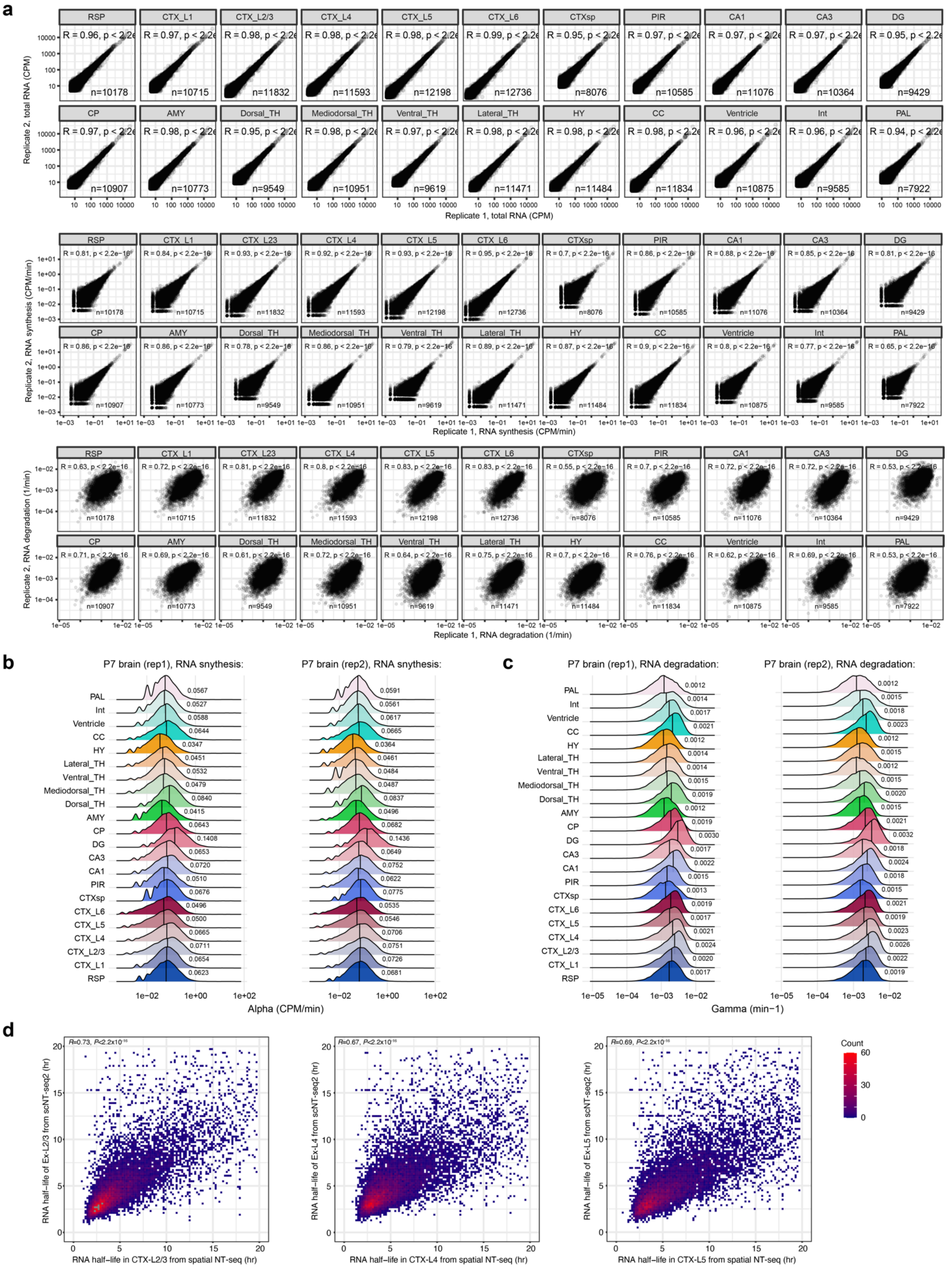


**Supplementary Fig. 12 | Benchmarking spatial RNA kinetics landscapes across specific brain regions.**

**a.** Scatter plots showing the Pearson's correlation between two replicates for total RNA abundance (top), RNA synthesis rates (middle), and degradation rates (bottom) across 22 brain regions from P7 mouse brain sections.
**b.** Density ridgeline plot illustrating the distribution of brain-region-specific RNA synthesis rates (CPM/min) between two replicates.
**c.** Density ridgeline plot illustrating the distribution of brain-region-specific RNA degradation rates (1/min) between two replicates.
**d.** Scatter plots showing the Pearson's correlation between single-cell (y-axis) and spatial (x-axis) NT-seq experiments for RNA half-life (hr) across three matched cortical cell-types/regions.

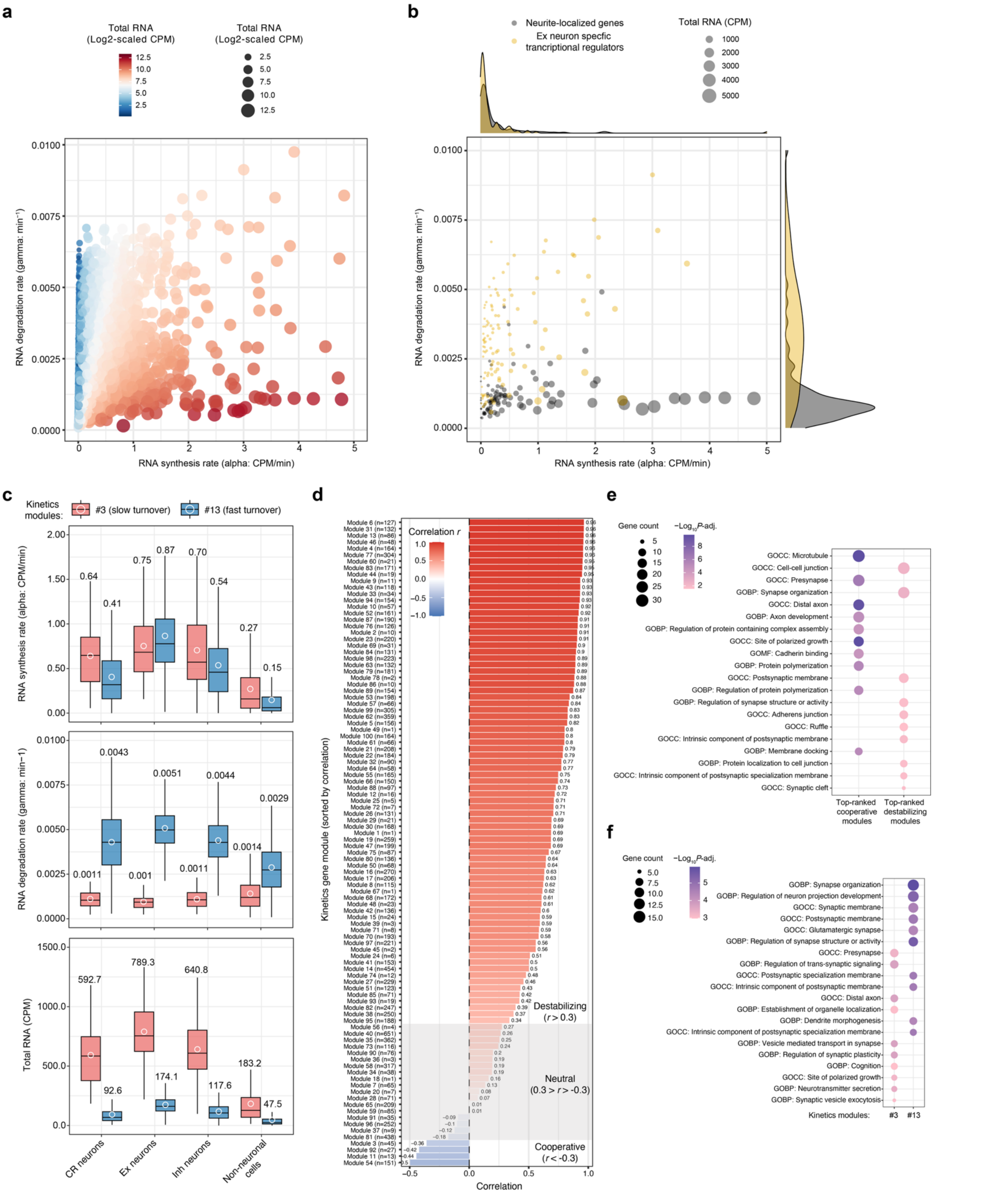


**Supplementary Fig. 13 | *In vivo* transcriptome-wide RNA kinetics landscapes across diverse cell types in the mouse brain.**

**a.** Scatter plot showing transcriptome-wide RNA synthesis (x-axis: CPM/min) versus degradation (y-axis: $min^{-1}$) rates in cortical Ex neurons of P7 mouse brains. Total RNA levels are represented by dot size and colors (log2-scaled).

**b.** Scatter plot showing RNA synthesis (x-axis: CPM/min) versus degradation (y-axis: $min^{-1}$) rates in cortical Ex neurons of P7 mouse brains for Ex neuron specific transcriptional regulators [16] (yellow, n=115 genes; e.g. *Rorb*: 1.8-hr, *Satb1*: 2.1-hr, *Pou3f1*: 2.4-hr) and neurite-locailized/enriched genes [17] (black, n=101 genes; e.g. *Mapt*: 14.8-hr, *Ybx1*: 12.2-hr, *Tpt1*: 11.5-hr, *Actb*: 10.8-hr, and *Rps14*: 7.8-hr). Total RNA levels are represented by dot sizes. Many neurite-locailized/enriched genes are required to encode proteins associated with core functions such as local translation, cytoskeleton, mitochondria and neurite formation [17]. The marginal density plots depict the distributions of RNA synthesis (x-axis) and degradation (y-axis) for two gene groups.

**c.** Box plots comparing cell-type-specific RNA synthesis rates (alpha, top), degradation rates (gamma, middle) and total RNA abundance (bottom) between two neuron-specific kinetics modules, #3 (red, n=45 genes; slower turnover) and #13 (blue, n=86 genes; faster turnover), across three major cortical neuronal cell types (CR, Ex and Inh) and non-neuronal cells (as a control). The mean is shown as a white dot, with its value indicated on the top.

**d.** Pearson correlation coefficients (*r*) between RNA synthesis rates (alpha, CPM/min) and degradation rates (gamma, $min^{-1}$) across 15 cortical cell types for 100 RNA kinetics gene modules (same modules as in Fig. 4b). Based on these correlations, we defined three distinct RNA regulatory strategies for 100 kinetics gene modules as previously described [2,18]: cooperative ($r < -0.3$; n=4), neutral ($-0.3 \leq r \leq 0.3$; n=18), and destabilizing ($r > 0.3$; n=78).

**e.** Gene ontology enrichment analysis for genes in top 4 “cooperative” or “destabilizing” kinetics gene modules (defined in Supplementary Fig. 13d). Dot size corresponds to the number of genes in each category, and the level of statistical significance for enrichment is color-scaled to represent $log_{10}$-scaled *P*-adjusted values (hypergeometric test).

**f.** Gene ontology enrichment analysis for genes in kinetics modules #3 and #13. Dot size corresponds to the number of genes in each category, and the level of statistical significance for enrichment is color-scaled to represent $log_{10}$-scaled *P*-adjusted values (hypergeomtric test).

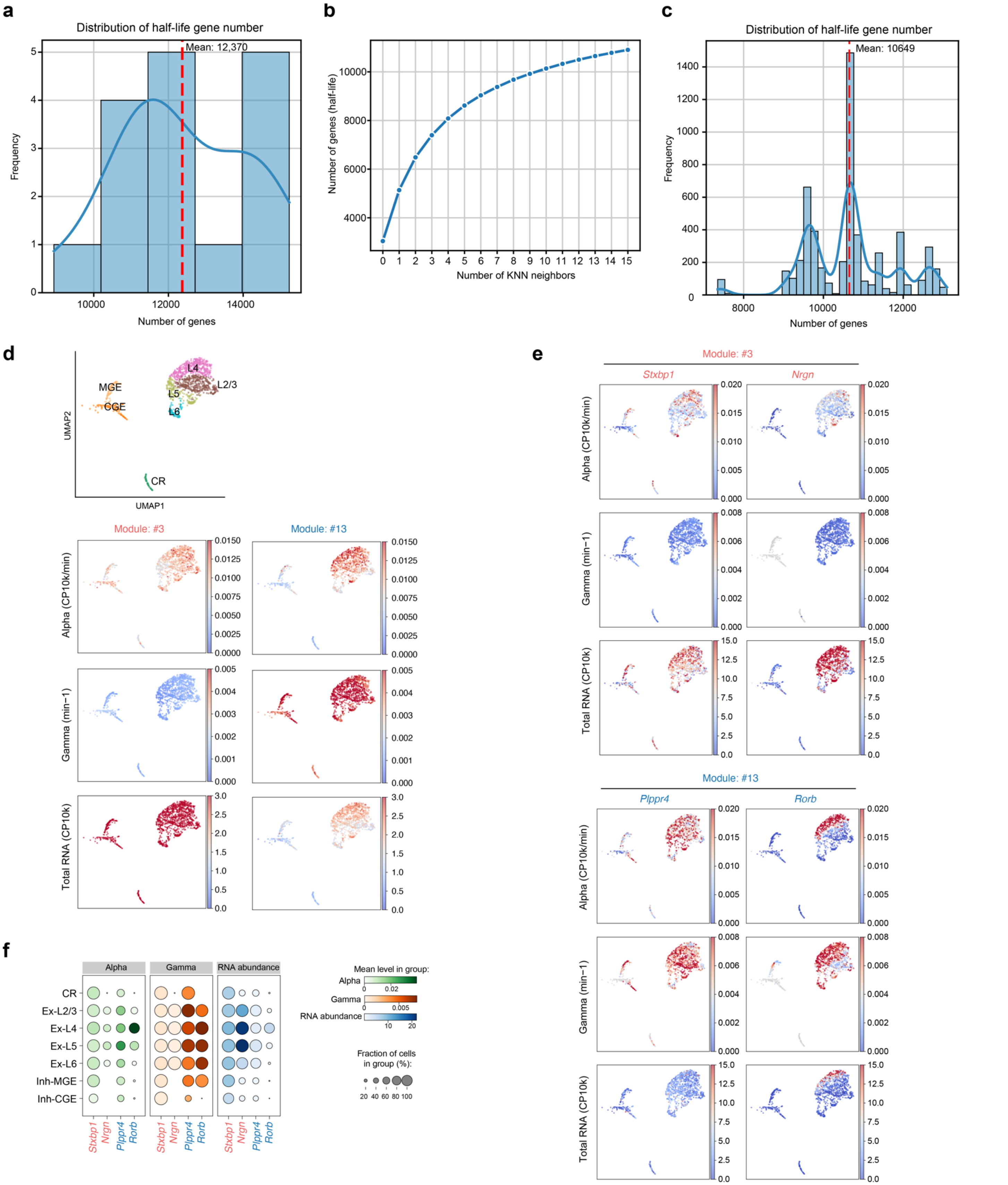


**Supplementary Fig. 14 | Development and benchmarking of *RNAKinetoScope* in analysis of single-cell RNA kinetics landscapes.**

**a.** Bar plots showing the distribution of gene numbers with calculated half-life (See Method for details) across 16 cortical cell types. The y-axis indicates the count of cell-types with distinct number of genes (x-axis).

**b.** Line plot illustrating the relationship between average number of genes with calculated half-life per cell (y-axis) with number of neighbors in *RNAKinetoScope* analysis (x-axis).

**c.** Bar plots showing the distribution of gene numbers with calculated half-life across individual cells via a *k*-nearest neighbor (KNN) graph approach (13 neighbors). The y-axis indicates the number of cells and the x-axis showing the specific gene numbers.

**d.** UMAP visualization showing seven cortical neuronal subtypes, colored by annotated cell-types, and single cell feature plots displaying aggregated RNA synthesis rate/alpha (top), degradation rate/gamma (middle), and total RNA abundance (bottom) for all genes in kinetics modules #3 (n=45 genes; slower turnover) and #13 (n=86 genes; faster turnover).

**e.** Single cell feature plots highlighting RNA synthesis rate/alpha (top), degradation rate/gamma (middle), and total RNA abundance (bottom) for four representative genes, including *Stxbp1* and *Nrgn* from module #3 (upper panels), along with *Plppr4* and *Rorb* from module #13 (lower panels).

**f.** Dot plots (bottom left) show the quantification of RNA synthesis/alpha (green), degradation/gamma (red) and abundance (blue) for four representative genes, including *Stxbp1* and *Nrgn* from module #3, along with *Plppr4* and *Rorb* from module #13 facross cortical neuronal subtypes.

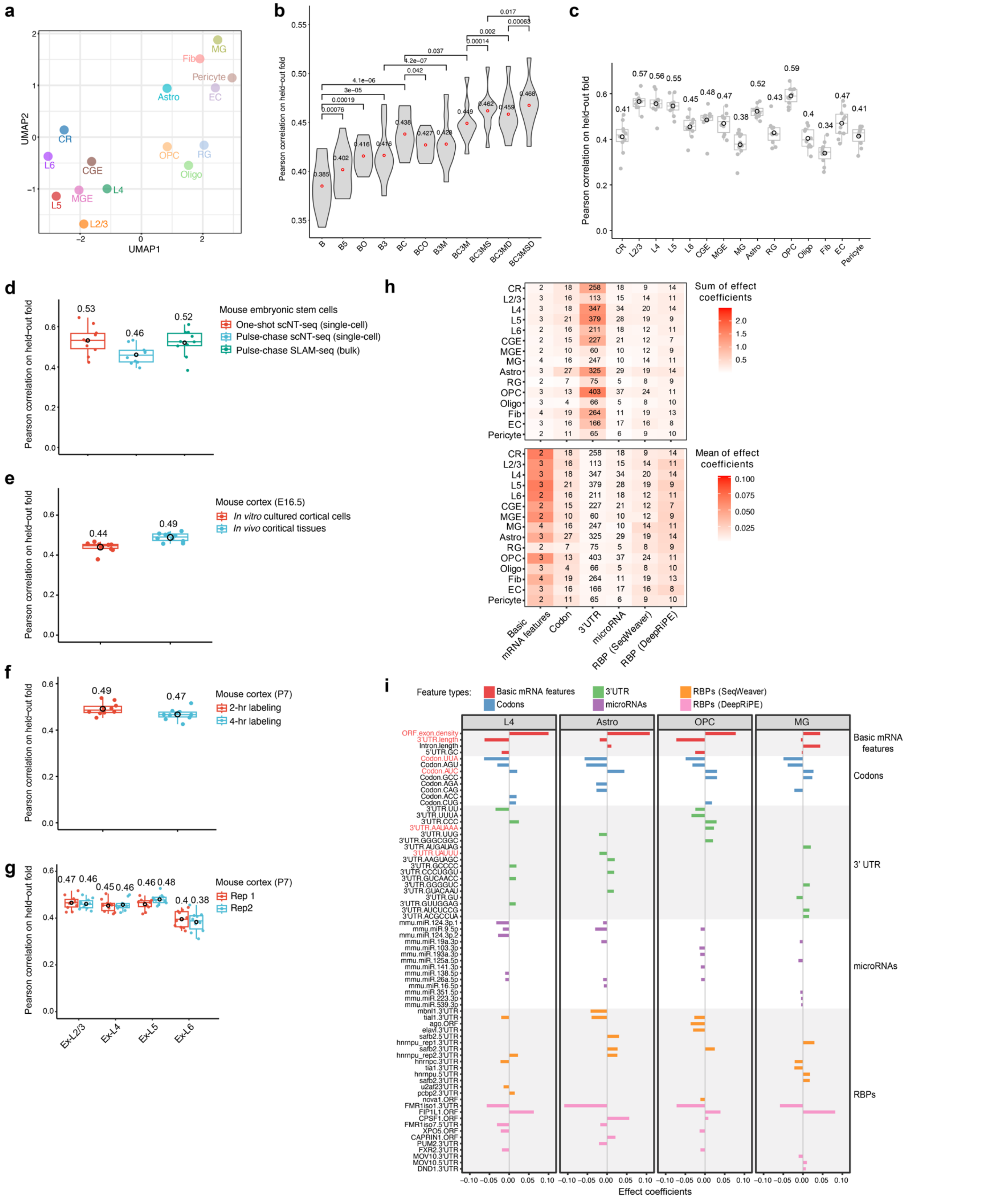


**Supplementary Fig. 15 | Evaluation of machine learning regression models for mRNA half-life prediction using scNT-seq2 datasets.**

**a.** UMAP visualization of annotated cell types based on the mRNA half-lives of top 3000 highly variable genes. Cell type abbreviations: CR, Cajal–Retzius cells; Ex-L2/3/4/5/6, layer specific excitatory neurons; Inh-CGE, caudal ganglionic eminence (CGE)-derived inhibitory neurons; Inh-MGE, medial ganglionic eminence (MGE)-derived inhibitory neurons; MG, microglia; Astro, astrocytes; RG, radial glial cells; OPC, oligodendrocyte precursor cells; Oligo, oligodendrocytes; Fib, fibroblast; EC, endothelial cells.

**b.** Violin plots showing the performance of trained models in excitatory neurons (Ex) on each of 10 held-out folds of data. Number of the input genes: 15,388. Sequence-derived feature groups: (1) “B”: basic mRNA features such as the length and G/C content of different functional regions (5’ UTR, ORF, and 3’ UTR), intron length, ORF exon junction density; (2) “5”, “O”, “3”: *k*-mer frequencies of length 1-7 in the 5’ UTR, ORF, and 3’ UTR; (3) “C”: codon frequencies; (4) “M”: predicted target scores of mammalian-conserved miRNA families by TargetScan [19]; (5) “S”: predicted binding of RBPs to the mRNA sequence by SeqWeaver [20]; (6) “D”: predicted binding of RBPs to the mRNA sequence by DeepRiPE [21]. RBP binding was predicted separately for different functional regions (5’ UTR, ORF, and 3’ UTR). To determine which of these feature sets would be informative for predicting RNA half-life, we analyzed a series of nested models which iteratively incorporated additional groups of features.

**c.** Box plots showing the performance of predictive modeling analysis across different cell types using common genes (n=2,853) on each of 10-fold held-out data.

**d.** Box plots showing the performance of predictive modeling analysis of mESC datasets on each of 10-fold held-out data. One-shot labeling and pulse-chase scNT-seq datasets [2] and bulk SLAM-seq data [13] are derived from previous studies. Number of the input genes: 1,889.

**e.** Box plots showing the performance of predictive modeling analysis of *in vitro* and *in vivo* 4sU-labeled E16.5 cortical neurons. Number of the input genes: *in vitro* Ex neurons - 14,416; *in vivo* Ex neurons - 14,823.

**f.** Box plots showing the performance of predictive modeling analysis of 2hr and 4hr labeling cortical datasets. Number of the input genes: 2hr labeling, 15,308; 4hr labeling, 15,388.

**g.** Box plots showing the performance of predictive modeling analysis of different cortical neuronal subtypes between two replicates. Number of the input genes: Ex-L2/3, 11,413; Ex-L4, 11,028; Ex-L5, 10,855; Ex-L6, 7,701.

**h.** Numbers and coefficients of each feature category across different cell types. Numbers of feature used in the model are listed, while the color indicates the summed coefficients (top) or average coefficients (bottom).

**i.** The top 5 features in each category associated with post-transcriptional regulation of RNA half-lives in four representative cell-types. Features are colored according to the categories.

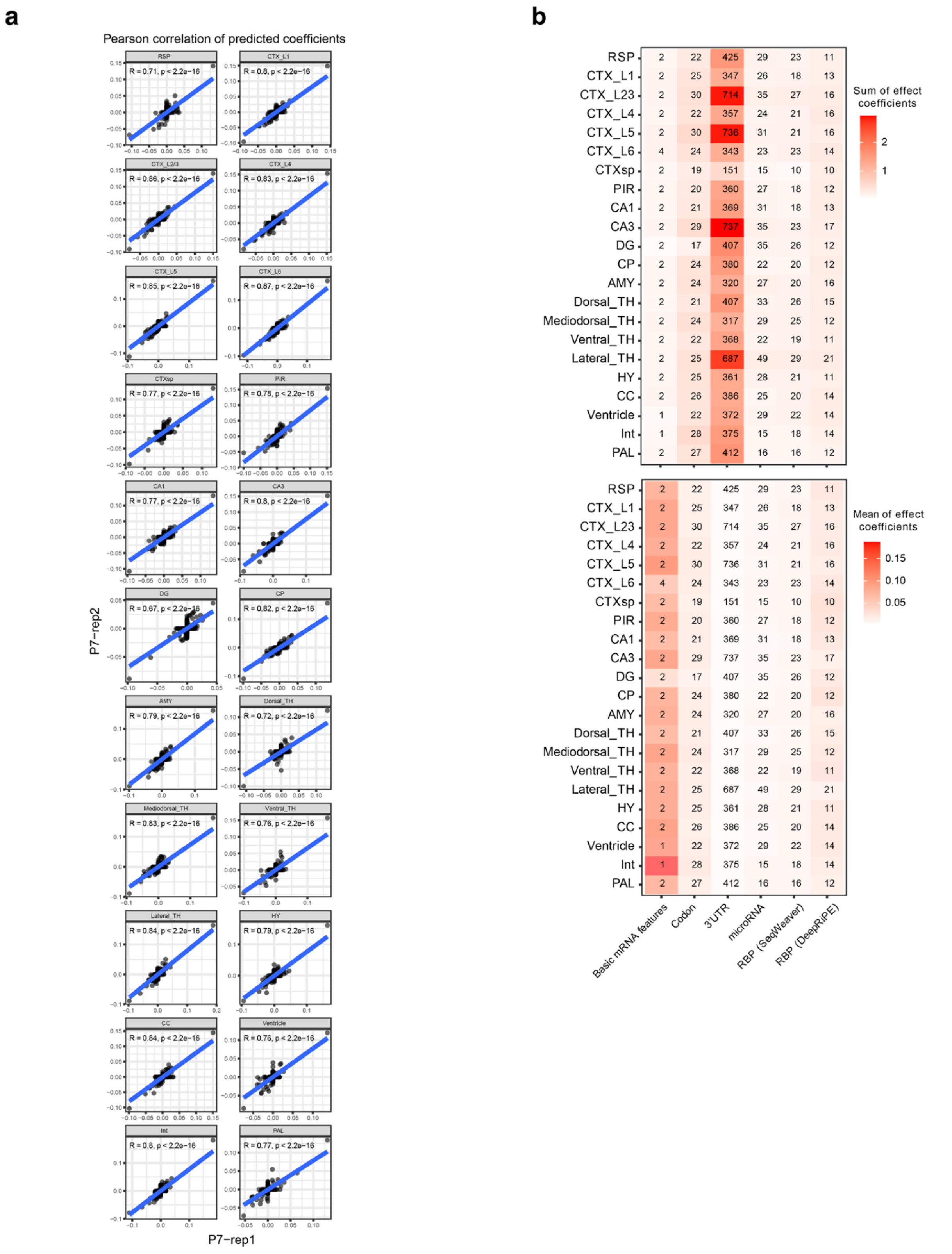


**Supplementary Fig. 16 | Evaluation of machine learning regression models for spatially resolved mRNA half-life prediction in mouse brains.**

- **a.** Scatter plots showing the Pearson's correlation between two replicates for predicted coefficients across 22 brain regions from P7 mouse brain sections. All features with non-zero predictd coefficients were plotted.
- **b.** Numbers and coefficients of each feature category across different spatial clusters. Numbers of feature used in the model are listed, while the color indicates the summed coefficients (top) or average coefficients (bottom).

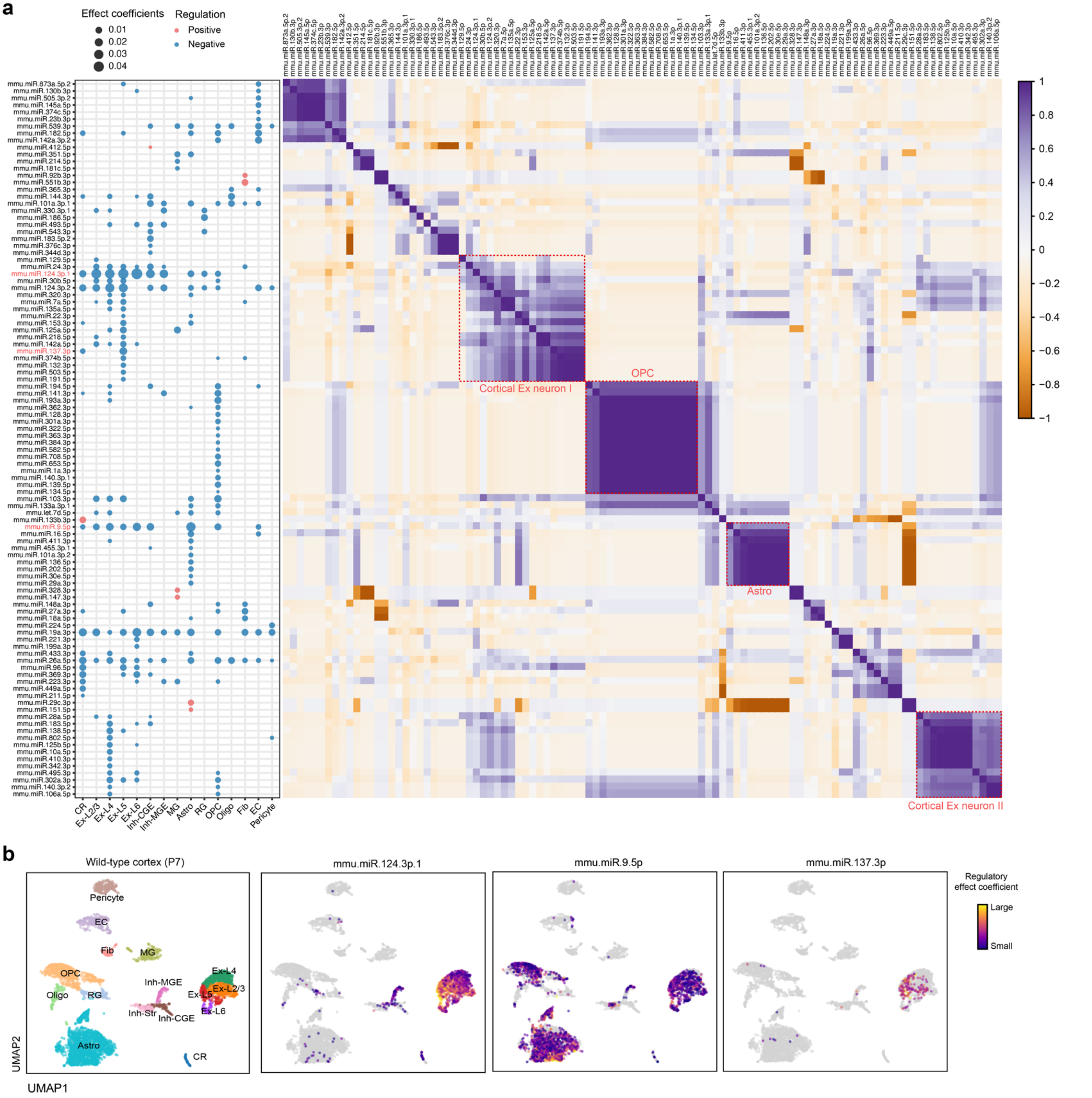


**Supplementary Fig. 17 | LASSO modeling of cell-type specific RNA half-life profiles reveals regulatory effects of miRNAs in regulating *in vivo* mRNA stability across diverse cortical cell types.**

**a.** The dot plot (left) and correlation matrix (right) of 102 miRNA features with non-zero coefficients across 15 cortical cell types. Dot plots showing their regulatory effect coefficients predicted by the optimized machine learning model (left). The magnitude and directionality of effect coefficients are indicated by the dot sizes and colors (red: positive regulators of mRNA stability; blue: negative regulators), respectively. Heat map showing the Pearson correlation matrix between these miRNA features (right). Representative clusters of miRNAs whose regulation is primarily cell-type specific are highlighted by red boxes.

**b.** UMAP visualization of the regulatory effect coefficients for three representative miRNAs in cortical cells derived from P7 mouse brains. In UMAP, the magnitude of regulatory effect coefficients is color-coded for each cell.

**a**

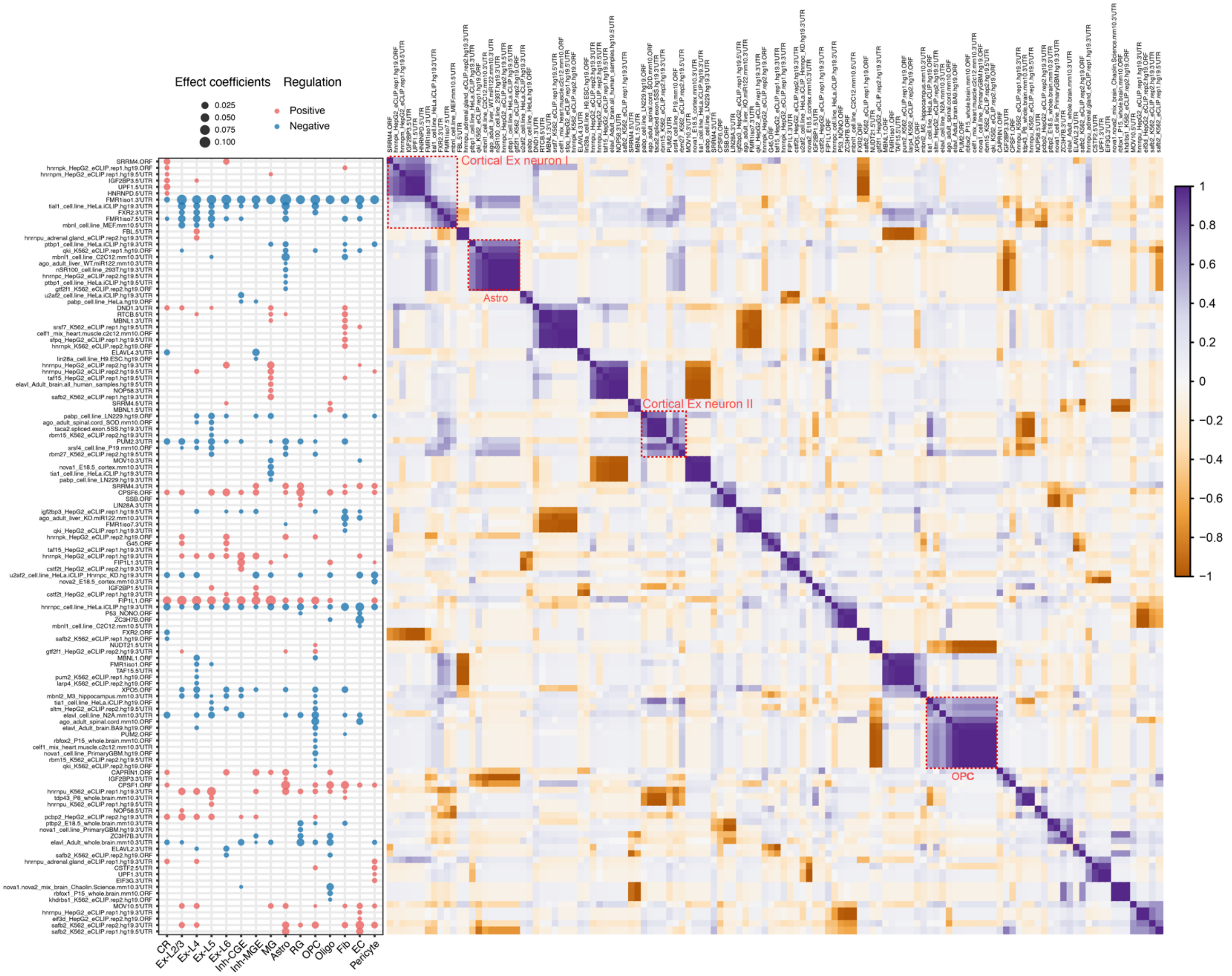


**b**

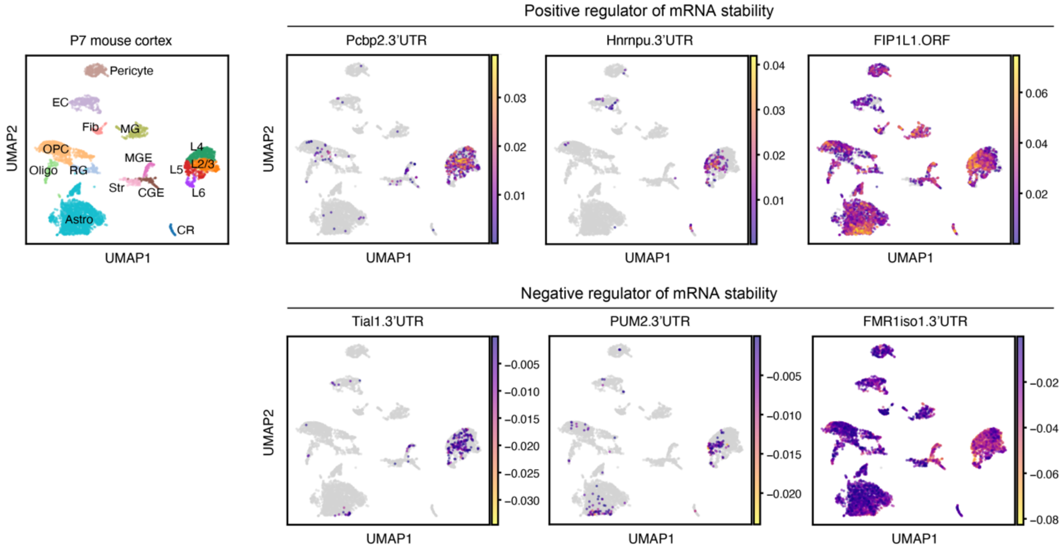


**Supplementary Fig. 18 | LASSO modeling of cell-type specific RNA half-life profiles reveals regulatory effects of RBPs in regulating *in vivo* mRNA stability across diverse cortical cell types.**

- **a.** The dot plot (left) and correlation matrix (right) of 122 RBP features with non-zero coefficients across 15 cortical cell types. Dot plots showing their regulatory effect coefficients predicted by the optimized machine learning model (left). The magnitude and directionality of effect coefficients are indicated by the dot sizes and colors (red: positive regulators of mRNA stability; blue: negative regulators), respectively. Heat map showing the Pearson correlation matrix between these RBP features (right).
- **b.** UMAP visualization of regulatory effect coefficients for six representative RBP features at single-cell resolution in P7 mouse cortex. Notably, three RBP features show stabilizing effects (Pcbp2.3UTR, Hnrnpu.3UTR, and FIP1L1.ORF), while other RBP features show destabilizing effects (Tial1.3UTR, PUM2.3UTR, and FMR1-iso1.3UTR).

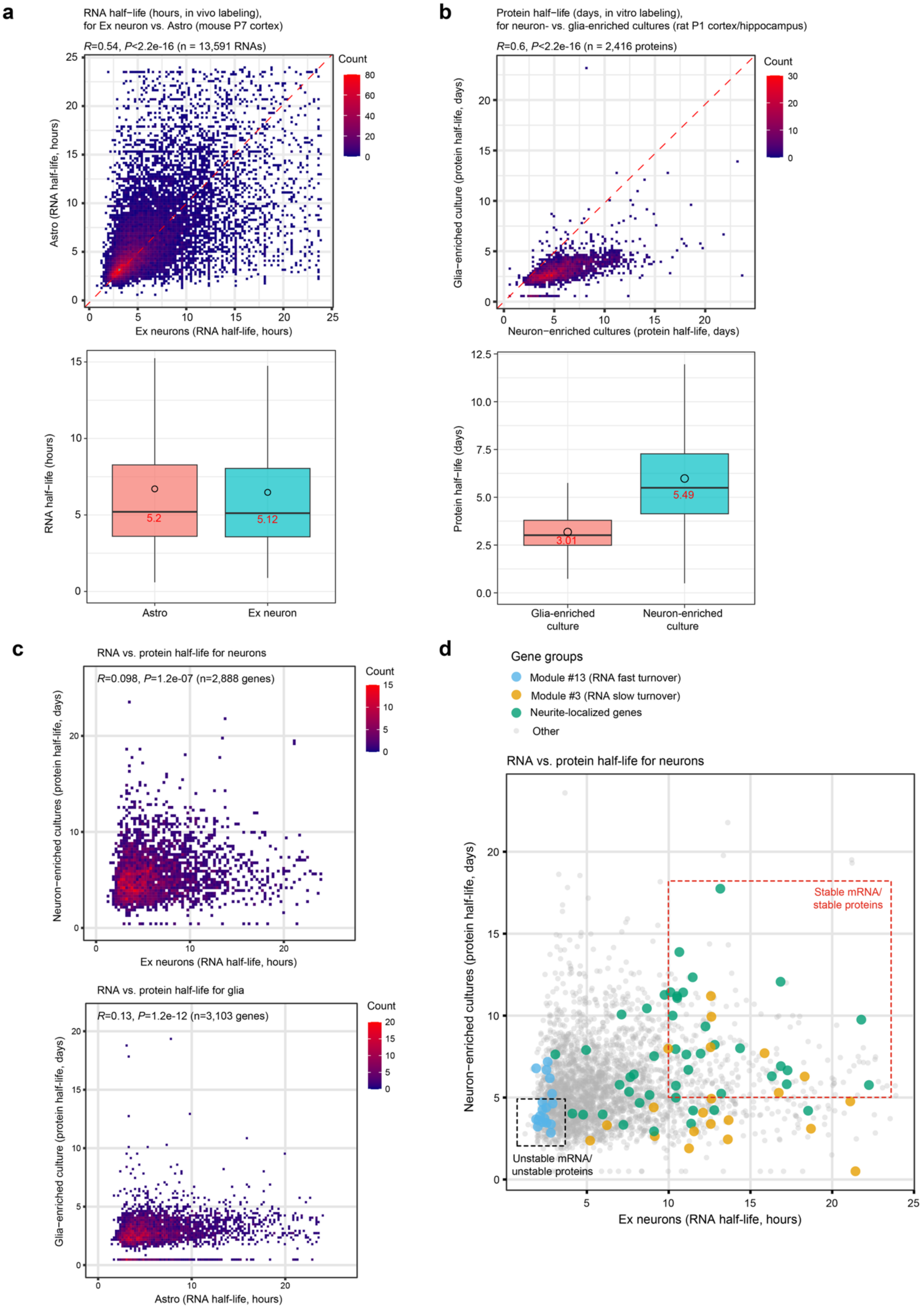


**Supplementary Fig. 19 | Comparison of RNA and protein half-lives in neurons and glia.**

- **a.** Scatterplots (top) and boxplots (bottom) comparing RNA half-lives (hours) of 13,591 unique transcripts between excitatory neurons (Ex) and astrocytes (Astro) in postnatal day 7 (P7) mouse cortex (scNT-seq2, this study). The Pearson correlation coefficient (R) and P-value are shown in the scatterplot; median half-lives for each cell type are indicated in the boxplot.
- **b.** Scatterplots (top) and boxplots (bottom) comparing protein half-lives (days) of 2,416 proteins between neuron- and glia-enriched cultures derived from postnatal day 1 (P1) rat cortex/hippocampus (cultured 18–19 days in vitro; data from LC-MS/MS with stable isotope labeling by amino acids in cell culture, SILAC) [22]. Median half-lives are shown in the boxplots.
- **c.** Scatterplots comparing RNA and protein half-lives in neurons (top; n=2,888 genes) and glia (bottom; n=3,103 genes). Pearson correlation coefficients (R) and P-values are reported.
- **d.** Scatterplots comparing RNA and protein half-lives in neurons for distinct gene groups: Module #13 (blue), Module #3 (yellow), neurite-localized genes (green), and all other genes (grey). A subset of genes associated with mRNA and protein stability are highlighted in the red box, whereas genes associated with mRNA and protein instability are highlighted in the black box.

**REFERENCES (for Supplementary Figures)**


1 Hughes, T. K. *et al.* Second-Strand Synthesis-Based Massively Parallel scRNA-Seq Reveals Cellular States and Molecular Features of Human Inflammatory Skin Pathologies. *Immunity* **53**, 878-894 e877, doi:10.1016/j.immuni.2020.09.015 (2020).

2 Qiu, Q. *et al.* Massively parallel and time-resolved RNA sequencing in single cells with scNT-seq. *Nature methods* **17**, 991-1001, doi:10.1038/s41592-020-0935-4 (2020).

3 Kleshchevnikov, V. *et al.* Cell2location maps fine-grained cell types in spatial transcriptomics. *Nat Biotechnol* **40**, 661-671, doi:10.1038/s41587-021-01139-4 (2022).

4 Okesli, A., Khosla, C. & Bassik, M. C. Human pyrimidine nucleotide biosynthesis as a target for antiviral chemotherapy. *Curr Opin Biotechnol* **48**, 127-134, doi:10.1016/j.copbio.2017.03.010 (2017).

5 Jaffrey, S. R. & Wilkinson, M. F. Nonsense-mediated RNA decay in the brain: emerging modulator of neural development and disease. *Nat Rev Neurosci* **19**, 715-728, doi:10.1038/s41583-018-0079-z (2018).

6 Zhao, B. S., Roundtree, I. A. & He, C. Post-transcriptional gene regulation by mRNA modifications. *Nature reviews. Molecular cell biology* **18**, 31-42, doi:10.1038/nrm.2016.132 (2017).

7 Zaccara, S., Ries, R. J. & Jaffrey, S. R. Reading, writing and erasing mRNA methylation. *Nature reviews. Molecular cell biology* **20**, 608-624, doi:10.1038/s41580-019-0168-5 (2019).

8 Bartel, D. P. Metazoan MicroRNAs. *Cell* **173**, 20-51, doi:10.1016/j.cell.2018.03.006 (2018).

9 Hochgerner, H. *et al.* Neuronal types in the mouse amygdala and their transcriptional response to fear conditioning. *Nat Neurosci* **26**, 2237-2249, doi:10.1038/s41593-023-01469-3 (2023).

10 Schwanhausser, B. *et al.* Global quantification of mammalian gene expression control. *Nature* **473**, 337-342, doi:10.1038/nature10098 (2011).

11 Dolken, L. *et al.* High-resolution gene expression profiling for simultaneous kinetic parameter analysis of RNA synthesis and decay. *RNA* **14**, 1959-1972, doi:10.1261/rna.1136108 (2008).

12 Schofield, J. A., Duffy, E. E., Kiefer, L., Sullivan, M. C. & Simon, M. D. TimeLapse-seq: adding a temporal dimension to RNA sequencing through nucleoside recoding. *Nature methods* **15**, 221-225, doi:10.1038/nmeth.4582 (2018).

13 Herzog, V. A. *et al.* Thiol-linked alkylation of RNA to assess expression dynamics. *Nature methods* **14**, 1198-1204, doi:10.1038/nmeth.4435 (2017).

14 Friedel, C. C., Dolken, L., Ruzsics, Z., Koszinowski, U. H. & Zimmer, R. Conserved principles of mammalian transcriptional regulation revealed by RNA half-life. *Nucleic Acids Res* **37**, e115, doi:10.1093/nar/gkp542 (2009).

15 Loedige, I. *et al.* mRNA stability and m(6)A are major determinants of subcellular mRNA localization in neurons. *Mol Cell* **83**, 2709-2725 e2710, doi:10.1016/j.molcel.2023.06.021 (2023).

16 Bravo Gonzalez-Blas, C. *et al.* SCENIC+: single-cell multiomic inference of enhancers and gene regulatory networks. *Nature methods* **20**, 1355-1367, doi:10.1038/s41592-023-01938-4 (2023).

17 von Kugelgen, N. & Chekulaeva, M. Conservation of a core neurite transcriptome across neuronal types and species. *Wiley Interdiscip Rev RNA* **11**, e1590, doi:10.1002/wrna.1590 (2020).

18 Battich, N. *et al.* Sequencing metabolically labeled transcripts in single cells reveals mRNA turnover strategies. *Science* **367**, 1151-1156, doi:10.1126/science.aax3072 (2020).

19 Agarwal, V., Bell, G. W., Nam, J. W. & Bartel, D. P. Predicting effective microRNA target sites in mammalian mRNAs. *Elife* **4**, doi:10.7554/eLife.05005 (2015).

20 Park, C. Y. *et al.* Genome-wide landscape of RNA-binding protein target site dysregulation reveals a major impact on psychiatric disorder risk. *Nat Genet* **53**, 166-173, doi:10.1038/s41588-020-00761-3 (2021).

21 Ghanbari, M. & Ohler, U. Deep neural networks for interpreting RNA-binding protein target preferences. *Genome Res* **30**, 214-226, doi:10.1101/gr.247494.118 (2020).

22 Dorrbaum, A. R., Kochen, L., Langer, J. D. & Schuman, E. M. Local and global influences on protein turnover in neurons and glia. *Elife* **7**, doi:10.7554/eLife.34202 (2018).

### Supplementary Note 1. Development and benchmark of scNT-seq2

Metabolic RNA labeling enables time-resolved, transcriptome-wide analysis of RNA dynamics at single-cell resolution by distinguishing newly synthesized ("new") and pre-existing ("old") RNAs using cell-permeable, nucleoside analogs such as 5-ethynyluridine (5-EU) or 4-thiouridine (4sU). Newly synthesized RNAs, incorporated with these analogs, can then be selectively identified through biochemical enrichment [1] or enrichment-free chemical conversion (e.g. T-to-C substitutions for 4sU labeling) [2-4]. Combined with scRNA-seq platforms – plate-based [scSLAM-seq [5]; NASC-seq [6]; scEU-seq [1]], combinatorial indexing [sci-fate [7,8]] or droplet-based [scNT-seq [9]] – this approach enables simultaneous profiling of new and old RNAs in the same cell. By allowing precise control over the duration and timing of metabolic RNA labeling, these methods capture more comprehensively key genes (e.g. TFs), enable more accurate inference on RNA velocity to inform on the future trajectory of a cell [8-10], and can quantitatively determining absolute RNA kinetics rates in a specific cell type/state [1,9]. Thus, it offers a powerful strategy for quantifying cell-type-specific RNA kinetics and tracking cellular dynamics [11,12].

To benchmark *in vivo* metabolic RNA labeling of intact mouse brains, we employed scNT-seq [9], a droplet-based, high-throughput platform optimized for capturing diverse brain cell types [13]. This method enables efficient on-bead chemical conversion of 4sU-labeled new transcripts, allowing joint profiling of "new" (marked by T-to-C substitutions) and "old" RNAs in whole cells by efficiently capturing both nuclear and cytoplasmic RNAs. Given the lower RNA content and labeling efficiency in *in vivo* brain cells compared to cultured cell lines, we developed and bscNT-seq2 to improve data quality and sensitivity for *in vivo* applications.

To overcome these limitations, scNT-seq2 incorporates key optimizations in second strand cDNA synthesis (2$^{nd}$ SS) reaction, including substitution of Klenow polymerase with Bst3, and redesigned random primers (N3G2N4B in place of N9) (**Supplementary Fig. 1a-e** and **Table 1-2** for Supplementary Note 1 below). Compared to original scNT-seq (TimeLapse + 2$^{nd}$ SS/Klenow with N9 random primers), these enhancements substantially improved read alignment rates (**Extended Data Fig. 1a** and **Supplementary Fig. 1e**), library complexity (**Extended Data Fig. 1b** and **Supplementary Fig. 1d**), chemical conversion of 4sU to cytosine analogs (T-to-C mutations in **Extended Data Fig. 1c** and **Supplementary Fig. 1f**), and reduced background mutations (non T-to-C mutations in **Extended Data Fig. 1c** and **Supplementary Fig. 1f**).

To enable quantitative analysis of RNA turnover in the mouse brain, we integrated *in vivo* metabolic RNA labeling with scNT-seq2 (**Fig. 1a**). Briefly, wild-type or UPRT transgenic mice were injected intraperitoneally with 4sU (wild-type) or 4tU (as a positive control, requiring transgenic expression of UPRT) for an experimentally controlled period (~2-4 hrs). Labeled cells were isolated from primary tissues, cryopreserved, and subjected to scNT-seq2 analysis.

**Table 1** (for Supplementary Note 1)

| Reaction buffer | Reaction mix |
|---|---|
| Klenow buffer A (SeqWell S3 condition) | 200 µl of reaction mixture (1× maxima RT buffer (ThermoFisher), 12% PEG-8000, 1 mM dNTPs (Clontech), 5 µM Template Switch Oligo-GAATG (TSO-GAATG: /5SpC3/AAGCAGTGGTATCAACGCAGAGTGAATG), 10 µM Template Switch Oligo-Random primer, and 1.25 U/ µl Klenow exo- (Enzymatics) |
| Klenow buffer B (scNT-seq condition) | 200 µl of reaction mixture (1× Blue buffer (Enzymatics), 4% Ficoll PM-400, 1 mM dNTPs (Clontech), 10 µM Template Switch Oligo-random primer, and 1.25 U/ µl Klenow exo- (Enzymatics) |
| Bst3-based reaction mixture | 200 µl of reaction mixture (1× Isothermal Amplification Buffer II (NEB), 6 mM MgSO4, 4% Ficoll PM-400, 1.4 mM dNTPs (Clontech), 10 µM Template Switch Oligo-random primer, and 0.4 U/µl Bst 3.0 DNA polymerase (NEB, M0374) |

**Table 2** (for Supplementary Note 1)

| Primer Name | Reference | Sequence of the oligo |
|---|---|---|
| TSO_N3GGN3B (SeqWell) | SeqWell S3 | AAG CAG TGG TAT CAA CGC AGA GTG A(N1:25252525)(N1) (N1)GG (N1)(N1)(N1) B |
| TSO-ATGGG | scNT-seq | AAG CAG TGG TAT CAA CGC AGA GTG AATGGG |
| TSO-N9 | scNT-seq | AAG CAG TGG TAT CAA CGC AGA GTG AAT (N1:25252525)(N1)(N1) (N1)(N1)(N1) (N1)(N1)(N1) |
| TSO-N3GN5 | This study | AAG CAG TGG TAT CAA CGC AGA GTG A(N1:25252525)(N1) (N1)G(N1) (N1)(N1)(N1) (N1) |
| TSO-N3GN4B | This study | AAG CAG TGG TAT CAA CGC AGA GTG A(N1:25252525)(N1) (N1)G(N1) (N1)(N1)(N1) (N2:00333433) |
| TSO-N4GN3B | This study | AAG CAG TGG TAT CAA CGC AGA GTG A(N1:25252525)(N1) (N1)(N1)G (N1)(N1)(N1) (N2:00333433) |
| TSO-N3GGN4B | This study | AAG CAG TGG TAT CAA CGC AGA GTG A(N1:25252525)(N1) (N1)GG (N1)(N1)(N1) (N1)(N2:00333433) |
| TSO-N3GGN5B | This study | AAG CAG TGG TAT CAA CGC AGA GTG A(N1:25252525)(N1) (N1)GG (N1)(N1)(N1) (N1)(N1)(N2:00333433) |

N1: 25%A, 25%C, 25%G, 25%T.
N2: 0%A, 33%C, 34%G, 33%T.

**Supplementary Note 2. Benchmarking the performance of the LASSO regression modeling of *in vivo* cell-type specific RNA half-life and regulome of RNA stability**

Recent advances in machine learning have enabled the predictive modeling analysis of mRNA stability regulatory landscapes using transcriptome-wide RNA half-life data [14,15] (**Fig. 5a**), but previous studies have been limited to *in vitro* cultured cells. To identify molecular features regulating *in vivo* cell-type-specific RNA stability, we trained and benchmarked a Least Absolute Shrinkage and Selection Operator (LASSO) regression model [14,15]

First, we first established the best performing model for collectively explaining *in vivo* cell-type-specific RNA half-life data with feature sets consisting of basic mRNA features, codon frequencies, 3' UTR *k*-mer motifs, miRNA target scores [16], and RBP binding scores [17,18]. This analysis showed the "BC3MSD" feature set as the optimal model for collectively explaining *in vivo* RNA half-life, consisting of basic mRNA features ("B", n=8 features), codon frequencies ("C", n=61), 3' UTR *k*-mer motifs ("3", n=21,844 features), predicted target scores of miRNA ("M", n=315 features) [16], and RBP binding scores predicted by SeqWeaver ("S", n=780 features) [17] or DeepRiPE ("D", n=177 features) [18] (**Supplementary Fig. 15b**). The choice of feature sets in our final model achieved optimal balance between model performance and feature complexity, aligning with previous ML modeling analysis of "ensemble" RNA half-life values from *in vitro* cultured mouse cells [14].

Second, the performance of trained models with various feature sets (i.e. measured by Pearson's correlation coefficients between predicted and observed RNA half-life) was systematically validated using held-out data and the 10-fold cross-validation strategy in both single-cell (**Supplementary Fig. 15b**) and spatial NT-seq (**Extended Data Fig. 9a**) datasets. Specifically, we applied the model to cell-type-specific measurements of mRNA half-life across 15 major cortical neuronal and non-neuronal cell-types (Pearson's correlation coefficients: mean ± s.d. = 0.46 ± 0.08 in **Supplementary Fig. 15c**). In addition, we evaluated the performance of the model on predicting *in vivo* RNA half-life of 22 spatial brain regions (mean ± s.d. = 0.44 ± 0.02 in **Extended Data Fig. 9b**), which is comparable to that of single-cell analysis. Importantly, the regulatory effects of top-ranked features are concordant between two replicates across 22 spatial brain regions (**Supplementary Fig. 16a**).

Finally, to assess the robustness of our model in RNA half-life prediction, we systematically evaluated it across various metabolic labeling-based RNA half-life analysis experiments (**Supplementary Fig. 15d-g**). Specifically, the trained model performs comparably across labeling strategies (one-shot vs. pulse-chase in **Supplementary Fig. 15d**), sequencing methods (single-cell vs. bulk RNA-seq in **Supplementary Fig. 15d**), sample origin (*in vitro* vs. *in vivo* 4sU labeling in **Supplementary Fig. 15e**), labeling durations (2-hr vs. 4-hr in **Supplementary Fig. 15f**), and is highly reproducible across biological replicates (**Supplementary Fig. 15g**).

**Supplementary Note 3. *In vivo* single-cell and spatial mapping of RBP-mediated regulation of RNA stability**

RBPs are a diverse class of proteins with the ability to recognize specific RNA sequences or structures, exerting regulatory control over various aspects of mRNA metabolism [19], including stability [20,21], yet their cell-type specific roles in modulating *in vivo* RNA stability remain poorly understood.

Utilizing LASSO regression-based predictions to assess the regulatory effects of a diverse array of RBPs on mRNA stability, we uncovered a total of 122 distinct RBP binding features, exhibiting cell-type specific patterns (**Supplementary Fig. 18a**). Specifically, we examined the effect coefficients pertaining to context-specific RBP bindings localized in the 5' UTR, ORF, and 3' UTR regions, as curated by deep learning-based computational models trained on experimental RBP binding datasets [17,18]. In contrast to miRNAs, which predominantly exert a negative influence on mRNA stability, our analysis revealed that 55 RBP binding features (constituting 45% of context-specific RBP binding features) exhibited positive effects on mRNA half-life (indicated by red dots in the left panel in **Supplementary Fig. 18a**), while 67 features (55%) showed negative impacts on mRNA half-life (depicted by blue dots in the left panel in **Supplementary Fig. 18a**). This is consistent with the notion that RBPs can elicit both stabilization and destabilization of target mRNAs, with these specific interactions influenced by various signaling cascades [14,19,22]. Moreover, spatially resolved RNA half-life analysis showed that, among 167 RBP binding features, 46% are associated with mRNA stabilization (red dots in **Extended Data Fig. 10a**), while the remainder exhibited negative impacts on mRNA stability (blue dots in **Extended Data Fig. 10a**).

To validate the predicted regulatory effects of RBPs, we examined RBPs with known roles in regulation of mRNA stability. Predicted effects of well-characterized RBPs aligned with findings from *in vitro* cultured cells, including mRNA stabilizing RBPs such as PCBP2 [23], hnRNP-K [24], hnRNP-U [25], and destabilizing RBPs including PUM2 [26], QKI [27], and TIA1/TIAL1 [20]. Among top-ranked predicted RBP features, 8 out of 9 features – encompassing either destabilizing (FMRiso1-3'UTR, PUM2-3'UTR, srsf4-ORF, rbm15-3'UTR, mbnl-5'UTR, and FXR2-3'UTR) or stabilizing (safb2-5'UTR, IGF2BP3-3'UTR) RBPs – were consistent with prior data from cultured mouse cells [14]. Furthermore, predicted *in vivo* regulatory effects of 5 out of 7 overlapping RBPs (destabilizing: DDX6, LARP4, and RBM15; stabilizing: CPSF6 and IGF2BP3) aligned with results derived from RBP knockdown experiments in cultured HepG2 and K562 cells [19]. The discrepancies likely reflect cell-type-specific regulation or methodological differences (bulk RNA-seq analysis of knockdown cell lines versus single-cell metabolic labeling RNA-seq). Thus, these results demonstrate the validity of our integrated experimental and computational approach in mapping the landscape of RBP-mediated post-transcriptional regulation of *in vivo* mRNA stability.

To enhance the resolution and sensitivity in detecting regulatory effects of RBP features on mRNA stability, we employed single-cell or spatial spot level RNA half-life measurements derived from *RNAkinetoScope* to compute their effects coefficients. The single-cell level analysis not only confirmed the predicted effects derived from pseudo-bulk/cell-type level mRNA half-life data, but also revealed both widespread (FMR1iso1.3'UTR and FIP1L1.ORF) and cell-type restricted regulatory effects (e.g. in cortical Ex neurons: Tial1.3'UTR, Pcbp2.3'UTR, Hnrnpu.3'UTR, and PUM2.3'UTR) of representative RBPs at the single-cell level (**Supplementary Fig. 18b**). Predictive modeling of spatial RNA half-life at the single spatial spot level identified both stabilizing and destabilizing RBPs with their regulatory effects enhanced in specific brain regions (e.g. Hnrnpu.3'UTR and PUM2.3'UTR enriched in DG/CA1 and specific cortical layers in **Extended Data Fig. 10b**).

**Supplementary Note 4. Comparison of RNA and protein turnover in neurons and glia**

To better understand whether and how RNA and protein turnover are coordinated in specific cell-types, we performed additional analyses comparing RNA and protein half-lives in neurons and glia using data from a published study [28].

First, we demonstrated that RNA life-life in Ex neurons and astrocytes (from P7 mouse cortex) are moderately correlated (Pearson correlation coefficient $R$=0.54, $P$<2.2e-16, n=13,591 common RNAs), and their median values are comparable (Astro: 5.20-hrs vs. Ex neuron: 5.12-hrs) when two biological replicates are combined (**Supplementary Fig. 19a**). Reanalysis of published protein half-life data from neuron- and glia-enriched cultures (derived from P1 rat cortex/hippocampus, cultured 18-19 days in vitro) confirmed that a similar moderate correlation (Pearson $R$=0.6, $P$ <2.2e-16, n=2,416 common proteins), but revealed that median protein half-lives in glia are ~45% shorter than in neurons (glia: 3.01-days vs. neuron: 5.49-days) (**Supplementary Fig. 19b**). These results suggest that glial proteins turn over faster than neuronal proteins, whereas RNA turnover rates are largely comparable. Nonetheless, differences in sample source (mouse cortex vs. rat cortex/hippocampus), developmental stage (P7 vs. P1), and experimental conditions (in vivo vs. in vitro) could also contribute to these discrepancies.

Next, we directly compared RNA and protein half-lives in both neuron and glia (**Supplementary Fig. 19c**). This analysis showed that RNA and proteins are only weakly correlated (in neuron: Pearson $R$=0.098, $P$=1.2e-7, n=2,888 genes; in glia: Pearson $R$=0.13, $P$=1.2e-12, n=3,103 genes), consistent with prior reports showing minimal concordance between RNA and protein stability when measured in parallel using metabolic labeling of RNA and protein [29]. Interestingly, subsets of genes showed coordinated stability: many long-lived neurite-enriched RNAs or genes in Module #3 (e.g. encoding pre-synaptic proteins) are associated with stable proteins (red box in **Supplementary Fig. 19d**), whereas many genes in module #13 (e.g. encoding neuron-specific TFs) exhibited both RNA and protein instability (black box in **Supplementary Fig. 19d**). These findings suggest that genes with matched mRNA and protein stability may share common cellular functions or subcellular localizations.

**References (for supplementary notes)**

17 Park, C. Y. *et al.* Genome-wide landscape of RNA-binding protein target site dysregulation reveals a major impact on psychiatric disorder risk. *Nat Genet* **53**, 166-173, doi:10.1038/s41588-020-00761-3 (2021).

18 Ghanbari, M. & Ohler, U. Deep neural networks for interpreting RNA-binding protein target preferences. *Genome Res* **30**, 214-226, doi:10.1101/gr.247494.118 (2020).

19 Van Nostrand, E. L. *et al.* A large-scale binding and functional map of human RNA-binding proteins. *Nature* **583**, 711-719, doi:10.1038/s41586-020-2077-3 (2020).

20 Akira, S. & Maeda, K. Control of RNA Stability in Immunity. *Annu Rev Immunol* **39**, 481-509, doi:10.1146/annurev-immunol-101819-075147 (2021).

21 Li, W., Deng, X. & Chen, J. RNA-binding proteins in regulating mRNA stability and translation: roles and mechanisms in cancer. *Semin Cancer Biol* **86**, 664-677, doi:10.1016/j.semcancer.2022.03.025 (2022).

22 Schoenberg, D. R. & Maquat, L. E. Regulation of cytoplasmic mRNA decay. *Nat Rev Genet* **13**, 246-259, doi:10.1038/nrg3160 (2012).

23 Ji, X., Kong, J. & Liebhaber, S. A. An RNA-protein complex links enhanced nuclear 3' processing with cytoplasmic mRNA stabilization. *EMBO J* **30**, 2622-2633, doi:10.1038/emboj.2011.171 (2011).

24 Thiele, B. J. *et al.* RNA-binding proteins heterogeneous nuclear ribonucleoprotein A1, E1, and K are involved in post-transcriptional control of collagen I and III synthesis. *Circ Res* **95**, 1058-1066, doi:10.1161/01.RES.0000149166.33833.08 (2004).

25 Yugami, M., Kabe, Y., Yamaguchi, Y., Wada, T. & Handa, H. hnRNP-U enhances the expression of specific genes by stabilizing mRNA. *FEBS Lett* **581**, 1-7, doi:10.1016/j.febslet.2006.11.062 (2007).

26 Goldstrohm, A. C., Hall, T. M. T. & McKenney, K. M. Post-transcriptional Regulatory Functions of Mammalian Pumilio Proteins. *Trends Genet* **34**, 972-990, doi:10.1016/j.tig.2018.09.006 (2018).

27 Doukhanine, E., Gavino, C., Haines, J. D., Almazan, G. & Richard, S. The QKI-6 RNA binding protein regulates actin-interacting protein-1 mRNA stability during oligodendrocyte differentiation. *Mol Biol Cell* **21**, 3029-3040, doi:10.1091/mbc.E10-04-0305 (2010).

28 Dorrbaum, A. R., Kochen, L., Langer, J. D. & Schuman, E. M. Local and global influences on protein turnover in neurons and glia. *Elife* **7**, doi:10.7554/eLife.34202 (2018).

29 Schwanhausser, B. *et al.* Global quantification of mammalian gene expression control. *Nature* **473**, 337-342, doi:10.1038/nature10098 (2011).
